# Nicotinic acetylcholine receptor partial antagonist polyamides from tunicates and their predatory sea slugs

**DOI:** 10.1101/2021.05.17.444536

**Authors:** Noemi D. Paguigan, Jortan O. Tun, Lee S. Leavitt, Zhenjian Lin, Kevin Chase, Cheryl Dowell, Cassandra E. Deering-Rice, Albebson L. Lim, Manju Karthikeyan, Ronald W. Hughen, Jie Zhang, Randall T. Peterson, Christopher A. Reilly, Alan R. Light, Shrinivasan Raghuraman, J. Michael McIntosh, Baldomero M. Olivera, Eric W. Schmidt

## Abstract

In our efforts to discover new drugs to treat pain, we identified molleamines A-E (**1**-**5**) as major neuroactive components of the sea slug, *Pleurobranchus forskalii* and their prey, *Didemnum molle* tunicates. The chemical structures of molleamines were elucidated by spectroscopy and confirmed by the total synthesis of molleamines A (**1**) and C (**3**). Synthetic **3** completely blocked acetylcholine-induced calcium flux in peptidergic nociceptors (PNs) in the somatosensory nervous system. Compound **3** affected neither the α7 nAChR nor the muscarinic acetylcholine receptors in calcium flux assays. In addition to nociceptors, **3** partially blocked the acetylcholine-induced calcium flux in the sympathetic nervous system, including neurons from the superior cervical ganglion. Electrophysiology revealed a block of α3β4 (mouse) and α6/α3β4 (rat) nicotinic acetylcholine receptors (nAChRs), with IC_50_ values of 1.4 and 3.1 µM, respectively. Molleamine C (**3**) is a partial antagonist, reaching a maximum block of 76-82% of the acetylcholine signal and showing no partial agonist response. Molleamine C (**3**) may thus provide a lead compound for the development of neuroactive compounds with unique biological properties.

## INTRODUCTION

Mollusks often use small molecules or peptides in defense, predation, and signaling. Cone snails, for example, inject into prey animals a potent venom that has led to many drug candidates and an FDA-approved pain therapy *(Bjørn-Yoshimoto, et al., 2020)*. Beyond cone snails, the neurochemical diversity of mollusks has been relatively unexplored, prompting us to conduct a screening campaign that led to the discovery of mollusk compounds with potential to combat the opioid crisis *(Volkow and McLellan, 2016)*. We performed phenotypic screen using the mouse dorsal root ganglion (DRG), where peripheral neurons detecting pain, heat, cold, touch, limb position, and other sensory modalities are bundled. Each of these sensations is signaled by several cell types each expressing complex constellation of receptors and ion channels. We applied constellation pharmacology to determine the cell-type selectivity of compounds in a single assay and to rapidly focus on compounds that selectively target nociceptors or other desired sensory cell types. Constellation pharmacology also enables the identification of potential molecular targets that are relevant to disease progression *(Raghuraman, et al., 2020; Teichert, et al., 2014; Teichert, et al., 2015; Teichert, et al., 2012)*. Pleurobranchs are a family of mollusks that are well known for neuroactive compounds: tetrodotoxin was isolated from *Pleurobranchus maculata (Wood, et al., 2012)*, while an ergot alkaloid was isolated from *Pleurobranchus forskalii (Wakimoto, et al., 2013)*. Here, we describe a class of potential antinociceptive compounds isolated from the sea slug *P. forskalii* from the Solomon Islands. Like other mollusks from Family Pleurobranchidae, *P. forskalii* has a greatly reduced shell and copious unprotected tissues that may require a chemical defense. It is a nocturnal animal that preys on tunicates, especially *Didemnum molle. P. forskalii* accumulates typical *D. molle* metabolites, including macrocyclic cyanobactin peptides and diterpenes, presumably from the diet *(Tan, et al., 2013; Wesson and Hamann, 1996)*. The compounds are toxic, leading to a proposed role in chemical defense. While defensive metabolites are well studied in many species of shell-less mollusks *(Cimino and Ghiselin, 2009)*, to the best of our knowledge the ecological roles of natural products in pleurobranchs have not been experimentally tested. Here, we show that *P. forskalii* obtained from the Solomon Islands contains molleamines, small molecules that are structurally related to the previously described mollecarbamate/molleurea class of alkaloid natural products *(Issac, et al., 2017; Lu, et al., 2012)*. Using metabolomics, we show that molleamines and their structural relatives are ubiquitously found in *D. molle* tunicates throughout the region where the *P. forskalii* sample was collected, while they were absent from other tunicates from the Family Didemnidae, reinforcing the dietary origin of the compounds.

In constellation pharmacology assays, molleamine C (**3**) displayed selective activity in blocking the acetylcholine induced calcium influx in a subset of peptidergic nociceptors (PNs), as well as an additional activity in Aδ-low threshold mechanoreceptors (LTMRs) at higher doses. These neurons are directly relevant to pain sensation. Mechanistic investigations show that the major activity of **3** is due to partial antagonism of the major α3 and α6 nicotinic acetylcholine receptors (nAChRs) that are specifically present in those cells. While there are many nAChR agonists and antagonists, the only other selective partial antagonist of α3 nAChRs (AT-1001) is also a partial agonist. Molleamine C (**3**) may thus provide a new lead for neuroactive drug discovery.

## RESULTS

### Isolation and characterization of neuroactive molleamines

*P. forskalii* was collected at night at Lomousa Reef, Solomon Islands (S 09° 08’42.33” E 159° 06’01.50). *D. molle* specimens SI-074U, -075W, -123K, -124L, -128T, -222K, -223L, -226S, -228U, -382U, -386H, -389MA, -389MB, -457S, -462Y, -463H, -464K, -465L, -485H, and -486K were collected near the Russell Islands and Honiara Island, Solomon Islands. The cytochrome oxidase I gene sequence confirmed the field identification (deposited in GenBank, MW663488). The *P. forskalii* ethanolic extract was potently active in a DRG assay, in particular blocking the effects of ATP on a subset of neurons. Assay-guided fractionation afforded five compounds, the molleamines, that were responsible for the observed activity.

Molleamine A (**1**) was isolated as an amorphous, colorless solid with a molecular formula of C_17_H_20_N_2_O, indicating nine degrees of unsaturation, as determined by HRMS and NMR data. The ^1^H NMR spectrum of **1** indicated the presence of an amide group (δ_H_ 8.55), an amine (δ_H_ 7.84), four methylenes, and nine aromatic protons. ^13^C and HSQC NMR data were consistent with the above units, which together accounted for all unsaturations. COSY and HMBC correlations from H-2 to C-8 and C-3 (Subunit A), H-4 to C-2 (Subunit A), and H-2 to C-3, C-4 (Subunit C) and H-8 to C-2 (Subunit C) enabled assembly of phenethylamine and 2-(2-aminoethyl)benzoic acid (AEBA) moieties (**Figure 1, Figure 1-figure supplement 1**). MS/MS data provided evidence that these two moieties were connected (**Figure 1-figure supplement 5**), and chemical synthesis of **1** from commercially available starting units confirmed the assignment. Molleamine B (**2**) differed from **1** by an oxygenation. This difference could be explained by a pair of *ortho*-coupled aromatic doublets (δ_H_ 7.84 and δ_H_ 6.70) and a phenolic OH signal at δ_H_ 9.21, indicating that the phenyl group in **1** was oxygenated in **2**, forming a *p*-hydroxyphenyl moiety. All NMR and MS data were consistent with this assignment.

**Figure 1.**
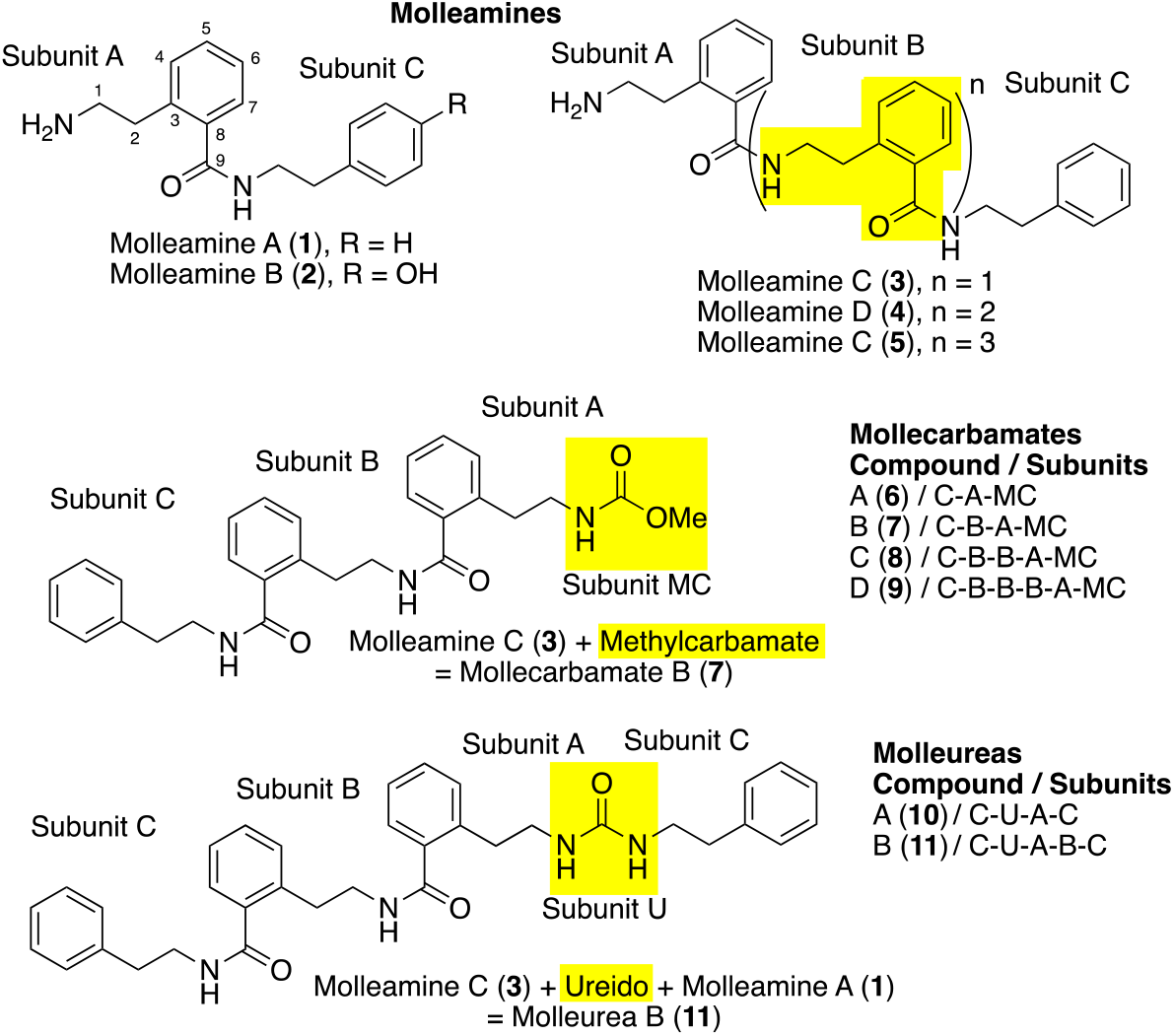
Molleamines isolated from the shell-less mollusk *P. forskalii*, compared with previously identified mollecar-bamates and molleureas from *P. forskalii*’s prey organism, the tunicate *D. molle*. All compounds contain a repeating 2-(2-aminoethyl)benzoic acid (AEBA) moiety (subunits A and B), capped with phenylethylamine or hydroxylphenylethylamine (subunit C). Highlighting shows characteristic features that define each compound class. With 34 supplements (**Figure supplement 1-33, Table supplement 1**).

Molleamine C (**3**) was isolated as an amorphous colorless solid with a molecular formula of C_26_H_29_N_3_O_2_, indicating 14 degrees of unsaturation. In comparison to **1, 3** incorporated nine more carbon atoms, nine more hydrogen atoms, one more nitrogen atom, and one more oxygen atom. The ^1^H NMR spectrum of **3** revealed two amide protons (δ_H_ 8.68 and δ_H_ 8.46), an amine (δ_H_ 7.88), 13 overlapping aromatic protons (δ_H_ 7.2-7.4), and six methylene groups (δ_H_ 2.84 - δ_H_ 3.47). The ^13^C NMR and HSQC data of compound **3** indicated the presence of two amide carbonyls, 18 aromatic carbons, and six methylenes. COSY and HMBC correlations from the above structural units supported one phenethylamine and two AEBA units. Key HMBC correlations from H-7 (subunit A) and H-1 (subunit B1) to amide carbonyl C-9 (subunit A), and from H-7 (subunit B1) and H-1 (subunit C) to amide carbonyl C-9 (subunit B1) allowed connection of the subunits as shown (**Figure 1-figure supplement 1**). The MS/MS fragment ions *m/z* 105, *m/z* 148, *m/z* 252, which were comparable to those of **1**, and new fragments *m/z* 269 and *m/z* 399 (**Figure 1-figure supplement 13**) supported the link between these moieties. Finally, because of the limited quantities available for pharmacological testing, we performed a gram-scale total synthesis of **3** from commercially available starting material. Spectroscopic data and coelution of the natural and synthetic products under several different conditions proved the identity of the synthetic and natural materials, and thus confirmed the structure of **3** as assigned (**Figure 1-figure supplement 26-29**).

Molleamines C (**4**) and D (**5**) were similarly elucidated, the differences being extension by one and two additional AEBA units, respectively. The assignments of **4** and **5** were supported by extensive NMR and MS data. The only exception was that the limited quantity available for **5** made it difficult to obtain a clear ^13^C NMR spectrum. However, the remaining data and close similarity to the spectrum of **4** strongly supported the structure as shown.

Compounds **1**-**5** all feature the unusual AEBA repeating units in their structures, which to the best of our knowledge have not been described in nature previously except in a series of molleureas isolated from the tunicate, *D. mole (Issac, et al., 2017; Lu, et al., 2012)*. Molleamines are structurally identical to a subset of the molleureas, except that they lack the ureido linkage.

In the same field trip in 2018 in which we obtained the *P. forskalii* sample, we also collected *D. molle* specimens spanning the known color morphs *(Hirose, et al., 2010; Hirose, et al., 2009)*, and hundreds of specimens of other tunicates. *D. molle* color morphs are genetically very distinct, and thus *D. molle* is likely comprised of many different cryptic species. A metabolomics analysis of the sample set revealed that of 20 *D. molle* specimens sampled, all contained diverse compounds in the AEBA-containing natural product family, including compounds **1** and **3**-**5** isolated from *P. forskalii* in this study (**Figure 2**). These results suggest that *P. forskalii* consumes *D. molle*, from which it obtains and concentrates the bioactive molleamines. The molleamine potency in assays (see below) is several orders of magnitude higher than the previously reported activity for molleureas and mollecarbamates *(Issac, et al., 2017)*, although the assays previously performed on the latter compounds are quite different than what we have done here.

**Figure 2.**
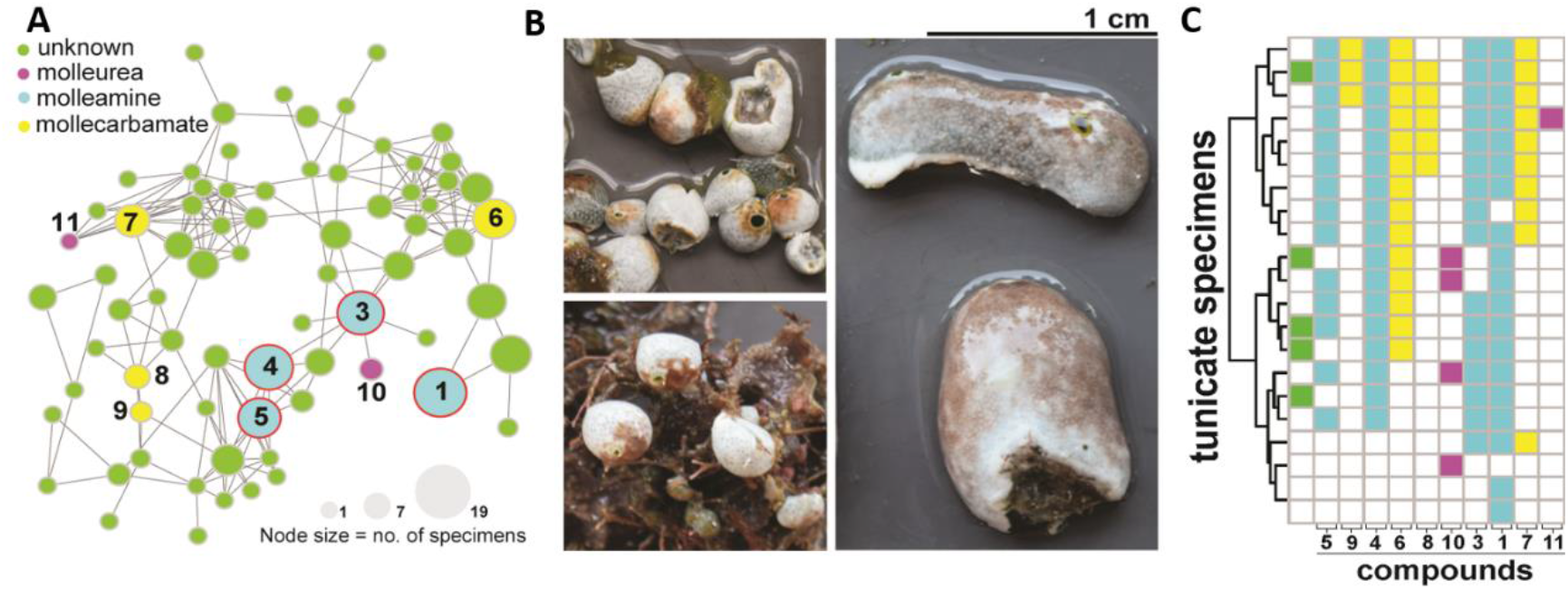
AEBA-containing metabolites are widely distributed in *D. molle* tunicates and their predatory slug, *P. forskalii*. **A**) A molecular network of AEBA-containing metabolites from *D. molle* from the Solomon Islands. Nodes are colored according to type of compound identified. Compounds **1**-**5** (blue circles) are also found in *P. forskalii*. The size of the nodes is proportional to the number of tunicate specimens in which the compound was identified. **B**) Representative morphotypes of *D. molle* tunicates from the Solomon Islands, from which the compounds shown in panel A were identified. **C**) Distribution of AEBA-containing compounds shown in the network in panel A. The coloring scheme is identical to that used in panel A. Molleamine compounds (blue) were first identified in the *P. forskalii* specimen described here, and they are widespread in the slug’s tunicate prey. Other mollureas (pink) and mollecarbamates (yellow) are also found in many of the animals. **Figure supplement 1**. Chemical structure and MS/MS fragmentation of compounds structurally-related to molleamines identified by metabolomics analysis of *D. molle* specimens. **Figure supplement 2**. Chemical structure and MS/MS fragmentation of compounds structurally-related to molleamines identified by metabolomics analysis of *D. molle* specimens.

### Molleamines selectively target specific cell types in the somatosensory nervous system

Using mouse DRG neurons and glia, we performed a calcium imaging-based high-content phenotypic screening assay *(Teichert, et al., 2014; Teichert, et al., 2015; Teichert, et al., 2012)*. Primary cultures from mouse DRG neurons were plated to obtain ∼2,000 cells/well, and the intracellular calcium levels were simultaneously monitored in all the cells using Fura-2-AM dye. Calcium traces were extracted for individual cells at the end of the experiment. Initial screening used periodic depolarizations of the DRG neurons with the application of extracellular ATP (20 µM), KCl (30 mM), or acetylcholine (ACh, 1 mM), which were expected to function primarily by activating purinergic receptors, voltage-gated Ca^2+^ channels, and acetylcholine receptors, respectively. At the end of each experiment, a series of pharmacological differentiators (κM-conopeptide RIIIJ (κM-RIIIJ), allylisothiocyanate (AITC), menthol, and capsaicin) aided in the identification of specific cell types. Thus, by interrogating cells with different pharmacological agents, the effects of extracts and pure compounds could be readily identified. In addition, cell size determination, fluorescent labeling with IB4, and the use of transgenic fluorescent markers to label calcitonin gene-related peptide (CGRP) expressing PNs, served to further differentiate cell types into at least 16 reproducible and distinct cell classes *(Giacobassi, et al., 2020)*.

Bioactivity screening of purified **1**-**5** at 20 µM revealed molleamine C (**3**) as a cell-type selective ligand (**Figure 3, Figure 3-table supplement 1-2**). Compound **3** potently and completely blocked the response of PN cells to ACh and modestly blocked the response of Aδ-LTMRs to ATP, but it had virtually no observable impact on DRG cells in response to KCl. This was a unique and promising phenotype in comparison to other compounds we have screened. Moreover, because PNs are therapeutic targets in pain, while Aδ-LTMRs are potential pain targets implicated in mechanical allodynia under neuropathic conditions *(Dhandapani, et al., 2018)*, we chose to focus on **3**.

**Figure 3.**
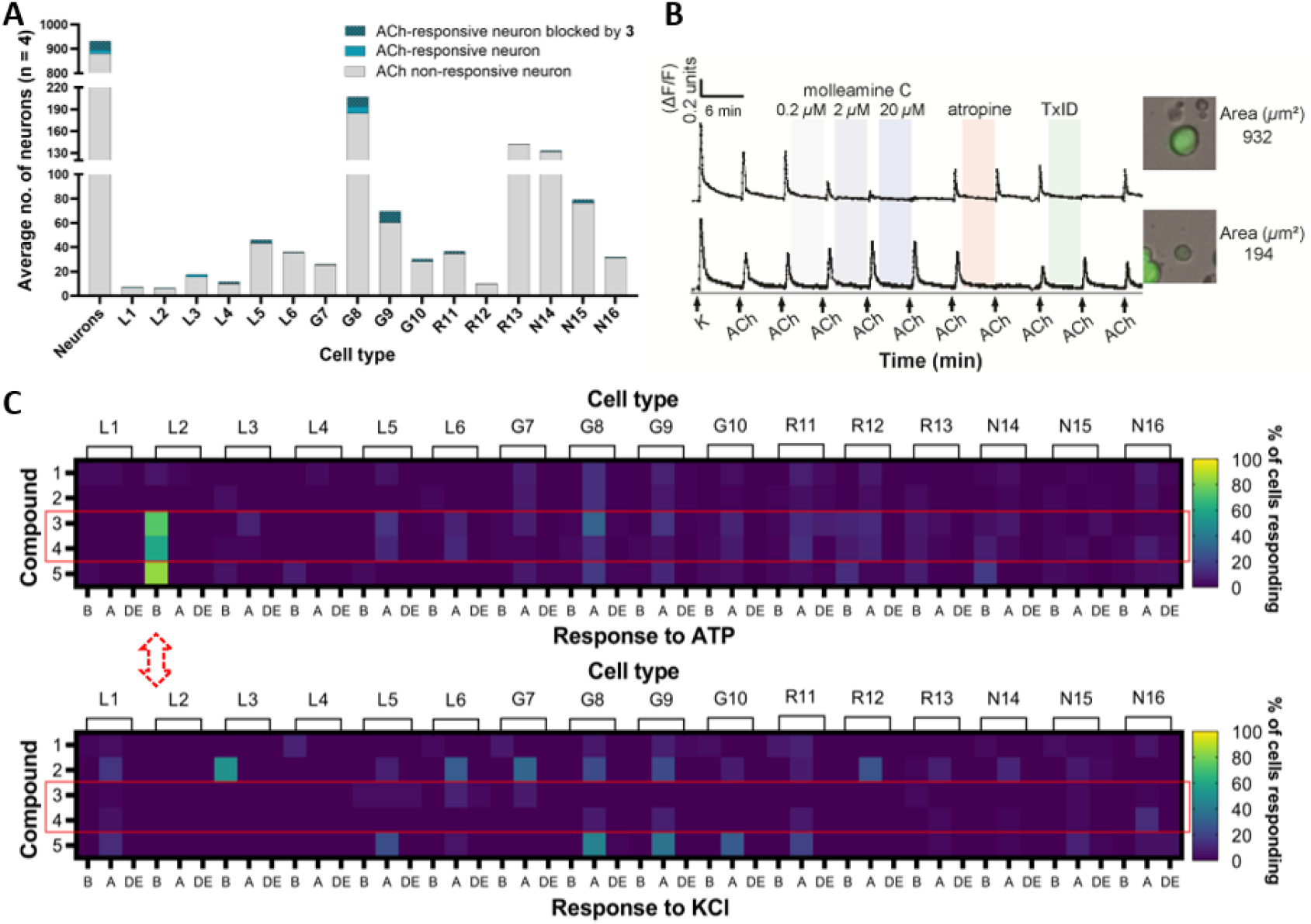
Constellation pharmacology of compounds **1**-**5** in DRG neurons. Descriptors L1-6, G7-10, R11-13, and N14-16 refer to neuronal cell types found in the DRG,*(Bosse, et al., 2021; Giacobassi, et al., 2020)* but importantly for this figure, G8 and G9 cells comprise pain sensors (PNs), while L2 cells are light touch-responsive Aδ-LTMRs. **A)** and **B)** Molleamine C (**3**) is a nAChR antagonist. Compound **3** (0.2-20 µM) was applied in experiments (n = 4) using ACh (1 mM), with an average of ∼900 neurons observed per experiment. **A)** Average number of neurons with ACh response blocked by **3** (20 µM). The *y*-axis shows the average number of neurons, while the *x*-axis indicates neuronal cell type. **B)** Representative individual DRG neurons responding to ACh (1 mM). The *y*-axis indicates intra-cellular [Ca^2+^], reflected in the normalized min/max fluorescence ratio of 340 nM/380 nM from the Fura-2-AM Ca^2+^ indicator. The *x*-axis is time (min), where ACh (1 mM) is repeatedly pulsed (arrows), with incubation of increasing concentrations of compound **3**, muscarinic receptor agonist atropine, and peptide nAChR antagonist TxID. Inset figures show the bright-field image of the corresponding cell (cross-sectional area in µm^2^). **C)** Compounds **3**-**4** selectively block ATP activation in Aδ-LTMRs, while response to KCl is unaffected. Compounds (applied at 20 µM) are in the *y*-axis, while the *x*-axis is activity observed in each cell type: amplification (A) of the response to ATP or KCl; blocking (B) of the response to ATP or KCl; and direct effects (DE), in which compounds directly depolarize cells. **Figure supplement 1**. Constellation pharmacology indicates that compounds **3**-**5** are selective against L2 DRG neurons, which are Aδ-low threshold mechanoreceptors (LTMRs). **Figure supplement 2**. Constellation pharmacology of molleamine C and transcriptomics analysis of neurons with responses to acetylcholine (ACh) blocked by molleamine C (20 µM) and TxID (1 µM). **Figure supplement 3**. Molleamine C does not affect α7-nAChRs. **Table supplement 1**. Census of effects elicited by compounds **1**-**5** on ATP-induced depolarization in 16 DRG neuronal subtypes screened in calcium imaging experiments. **Table supplement 2**. Census of effects elicited by compounds **1**-**5** on K^+^-induced depolarization in 16 DRG neuronal subtypes screened in calcium imaging experiments.

### Molleamine C (3) is an antagonist of nAChRs in peptidergic nociceptors

When we treated DRG neurons with ACh (1 mM), only a relatively small subset of cells responded (∼5% of neurons, **Figure 3A**). The strongest responses were observed in PNs, although only <10% of PNs responded. When molleamine C (**3**) was applied at 20 µM, the responses to ACh in many cells were almost completely blocked (**Figure 3B**). Block was dose dependent and could be observed at concentrations as low as 0.2 µM (**Figure 3B**). To differentiate nAChR (ion channel) versus muscarinic (GPCR) activity, either atropine (muscarinic antagonist)*(Zwart and Vijverberg, 1997)* or the α-conotoxin TxID (nicotinic antagonist)*(Luo, et al., 2013)* was applied in a calcium imaging experiment. In all cases, ACh responses that were blocked by TxID were also blocked by **3**. On the contrary, ACh responses that were blocked by atropine (in glia and some neurons) were not blocked by **3**. Taken together, these results indicate that **3** is an nAChR antagonist.

Agonists and positive allosteric modulators of nAChR subtypes α4β2, α6β4, and α7 are analgesic in various animal models, as are antagonists of α9α10 *(Christensen, et al., 2017; Cucchiaro, et al., 2008; Hone, et al., 2018; Limapichat, et al., 2014; Loram, et al., 2012; Romero, et al., 2017; Zheng, et al., 2020)*. To identify the specific nAChR subtype composition modulated by **3**, we performed calcium imaging experiments, and individual cells whose responses to ACh were blocked by **3** were picked for single-cell transcriptomic analysis. We found that nAChR genes encoding subunits α3 (Chrna3), α6 (Chrna6), β3 (Chrnb3), β4 (Chrnb4), and β2 (Chrnb2) are significantly expressed in neurons that respond to **3** (**Figure 3-figure supplement 2**). At 1 µM, TxID should completely block both the α3β4 and the α6β4 nAChR subtypes, supporting the activity of **3** as an antagonist of one or both receptors.

Previous studies have shown that DRGs functionally express the α7 nAChR *(Hone, et al., 2012; Smith, et al., 2013)* To observe the calcium transients elicited by the opening of α7 receptors, we applied the positive allosteric modulator PNU 120596 before depolarization with ACh. Molleamine C (**3**) was inactive against the elicited α7 nAChR activity (**Figure 3-figure supplement 3**).

To assess the functional effects of **3** on both α3β4 and α6β4 nicotinic acetylcholine receptors (nAChR), the compound was applied to *Xenopus laevis* oocytes expressing α3- and α6-containing nAChRs, measuring the responses to acetylcholine (ACh) at 100 µM. The compound exhibited partial antagonism of ACh-evoked currents mediated by mouse α3β4 nAChR and rat α6/α3β4 nAChR, with IC_50_ values (and 95% confidence intervals) of 1.43 (0.4 – 5.1) µM and 3.10 (1.9 – 5.0) µM, respectively (**Figure 4**). The partial block of α3β4 and α6/α3β4 plateaued at approximately 76% and 82%, respectively. Although **3** is a partial antagonist of the receptors, it exhibits complete block of calcium flux in PNs because the cells are unable to achieve the ion concentration necessary to induce depolarization in the assay conditions.

**Figure 4.**
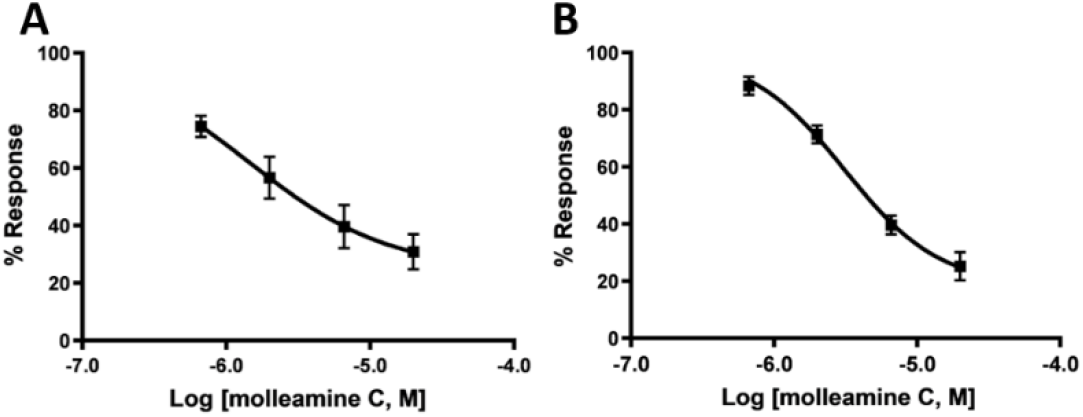
Compound **3** is a partial antagonist of nAChRs expressed in *X. laevis* oocytes, as determined by electrophysiology. The *x*-axis is log [**3**], while the *y*-axis is the response to ACh at varying [**3**], in comparison to ACh without **3**, given as the mean ± SEM from at least four separate oocytes. Ranges in parentheses are 95% confidence intervals. **A)** Mouse α3β4 nAChR, IC_50_ 1.43 (0.4 – 5.1) µM, Hill slope 0.89 (0.20 -2.0). **B)** Rat α6/α3β4 nAChR, IC_50_ 1.4 (1.9 – 5.0) µM, Hill slope 1.3 (0.67 - 1.90).

### Molleamine C (3) blocks ACh signaling in the sympathetic nervous system

Because α3- and α6-containing neurons are found in several important cell types in the peripheral nervous system, we aimed to determine the impact of **3** in those neurons. We performed constellation pharmacology assay using primary cultures of superior cervical ganglion (SCG) cells, which are involved in the fight-or-flight response. SCGs contain abundant α3β4-containing nAChRs, with a lower number of other nAChR types *(Simeone, et al., 2019)*. Consistent with this observation, in our hands almost all SCGs responded to ACh (300 µM) by depolarization and influx of calcium (**Figure 5**). When treated with **3** (20 µM), the influx of calcium was blocked in essentially all SCG neurons. This block was only complete in a subset of neurons, and partial block was observed in 95 % of ACh-responding neurons, consistent with a mixture of nAChR subtypes, not all of which respond to **3**, in this cell population. This data reinforce the primary role of **3** as a partial antagonist of α3β4 and α6/α3β4 nAChRs. This also represents the first application of constellation pharmacology to the sympathetic nervous system, showing the broad utility of the method in differentiating cells and investigating mechanism of action of neuroactive drugs.

**Figure 5.**
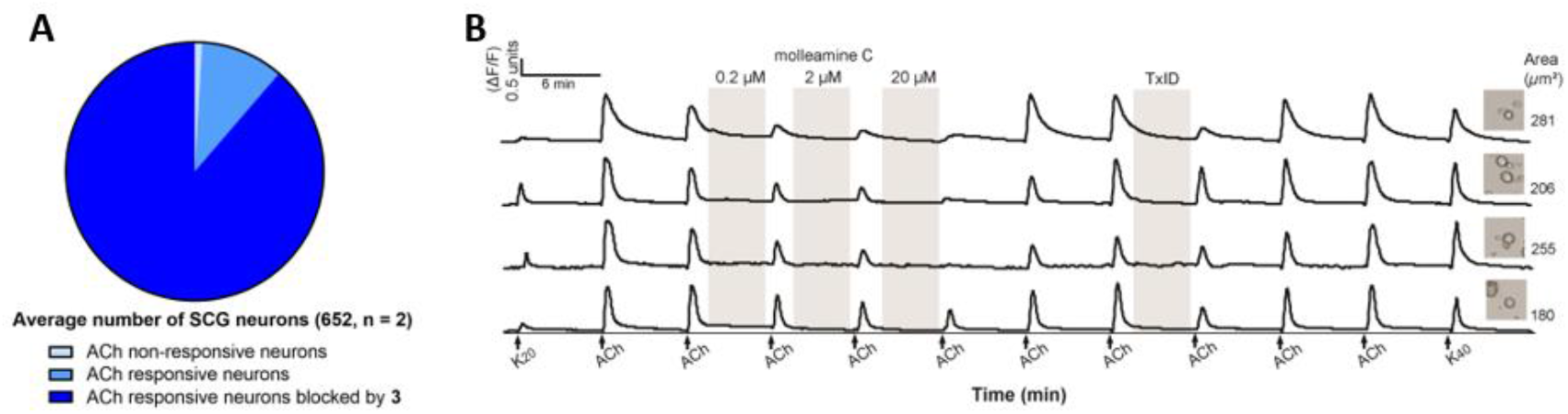
Constellation pharmacology indicates that **3** blocks ACh signaling in primary cultures of SCG neurons. SCG cells were treated with ACh (300 µM), **3** (0.2, 2 and 20 µM, and TxID (1 µM). **A)** Average number of neurons with ACh response blocked by **3** (20 µM) (n = 2). **B)** Representative individual SCG neurons responding to ACh. The *y*-axis indicates intracellular [Ca^2+^], reflected in the normalized min/max fluorescence ratio of 340 nM/380 nM from the Fura-2-AM Ca^2+^ indicator. The *x*-axis is time (min), where ACh (1 mM) is repeatedly pulsed (arrows), with incubation of **3** and TxID. Inset figures show the bright-field image of the corresponding cell (cross-sectional area in µm^2^).

### Molleamine C (3) indirectly blocks P2Y1 signaling in Aδ-LTMRs

At concentrations between 6.7 µM and 180 µM, **3** modestly but selectively blocked ATP signaling in Aδ-LTMRs (**Figure 6A**). Single-cell transcriptomics revealed the purinergic receptor P2Y1 as the major potential target in Aδ-LTMRs *(Zheng, et al., 2019)*. A selective P2Y1 agonist, MRS 2365 *(Lu, et al., 2007)*, was applied to DRG neurons, leading to selective, robust depolarization of Aδ-LTMRs (**Figure 6-table supplement 1**.). Molleamine C (**3**) at 20 and 180 µM selectively blocked the effect of MRS 2365 (**Figure 6B**), indicating selective activity against P2Y1-mediated signaling in mouse Aδ-LTMR neurons. However, **3** did not inhibit human P2Y1 in HEK293 cells, and P2Y1 is distributed in several cells in the DRG that were not blocked by **3**, indicating that P2Y1 itself is not blocked in Aδ-LTMRs, and instead the inhibition is due to an indirect effect provoked by **3**.

**Figure 6.**
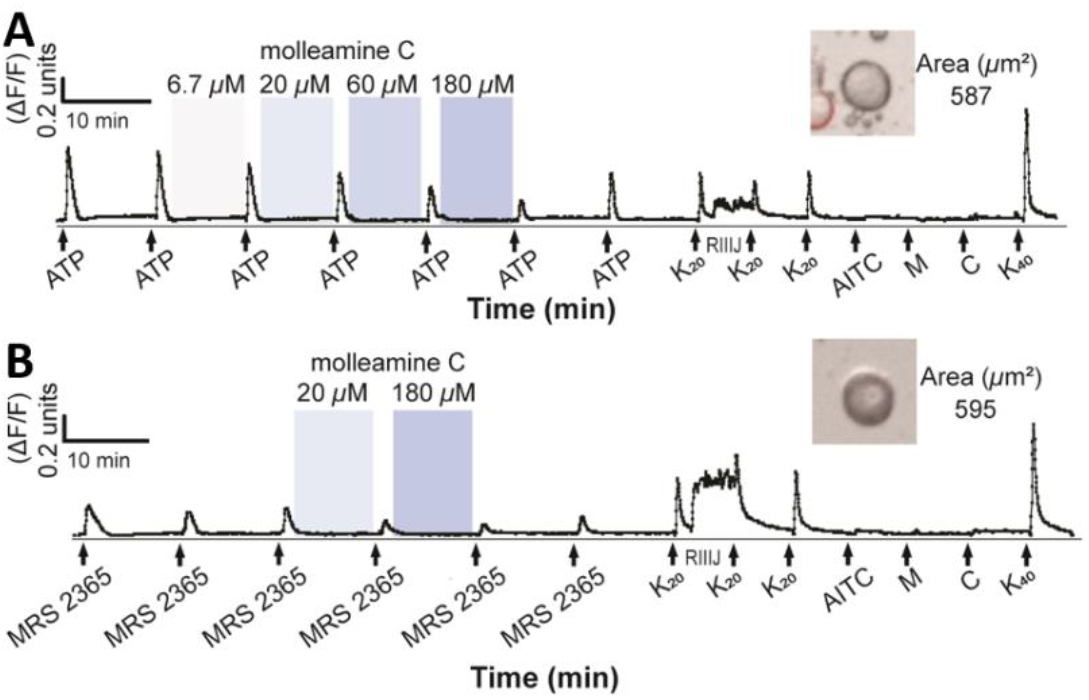
Compound **3** inhibits P2Y1-mediated responses in Aδ-LTMR DRG neurons. The *y*-axis indicates intracellular [Ca^2+^], reflected in the normalized min/max fluorescence ratio of 340 nM/380 nM from the Fura-2-AM Ca^2+^ indicator. The *x*-axis is time (min), where **A)** ATP (20 µM) or **B)** MRS 2365 (100 nM) activation is opposed by addition of (from 6.7 to 180 µM). Ligands used to differentiate cell types (K_20_, potassium chloride (20 mM); RIIIJ (1 µM); AITC (100 µM); M, menthol (400 µM); C, capsaicin (500 nM); K_40_, potassium chloride (40 mM)) are added at the end of the experiment. Inset figures show the bright-field image of the corresponding cell (cross-sectional area in µm^2^). **Figure supplement 1**. Compound **3** inhibits responses from L2 neurons after depolarization with a P2Y1 agonist MRS 2365 (100 nM). **Figure supplement 2**. Compound **3** does not inhibit P2Y1 activity in human embryonic kidney (HEK-293) overexpressing GCaMP6s. **Table supplement 1**. Census of effects elicited by **3** on MRS2365-induced depolarization in 16 DRG neuronal subtypes screened in two calcium imaging experiments.

### Preliminary *in vivo* evaluation of molleamine C (3)

Because of the effects of **3** dampening signals in pain relevant neurons PNs and Aδ-LTMRs, we assessed the impact of **3** on both acute and persistent pain perception in mice using a formalin assay *(Fu, et al., 2001)*. Compared with the vehicle, 5 mg/kg and 20 mg/kg doses of **3** decreased the pain response in Phase I (*p*<0.05). The highest dose of **3** at 30 mg/kg modestly attenuated the pain responses in both Phase I and II at *p* = 0.06 (**Figure 7**). The biphasic response in the formalin test is believed to reflect the direct activation of primary afferent sensory neurons (Phase I) and sensitization of the central nervous system in combination with inflammatory factors (Phase II) *(Hunskaar and Hole, 1987; McNamara, et al., 2007; Tjølsen, et al., 1992)*. Therefore, the effects of **3**, if supported by further studies, might be caused by both peripheral and central mechanisms. These results should be viewed as preliminary, as much more work is required to determine whether **3** has analgesic potential in therapy and to connect our observed molecular mechanisms to any *in vivo* activity.

**Figure 7.**
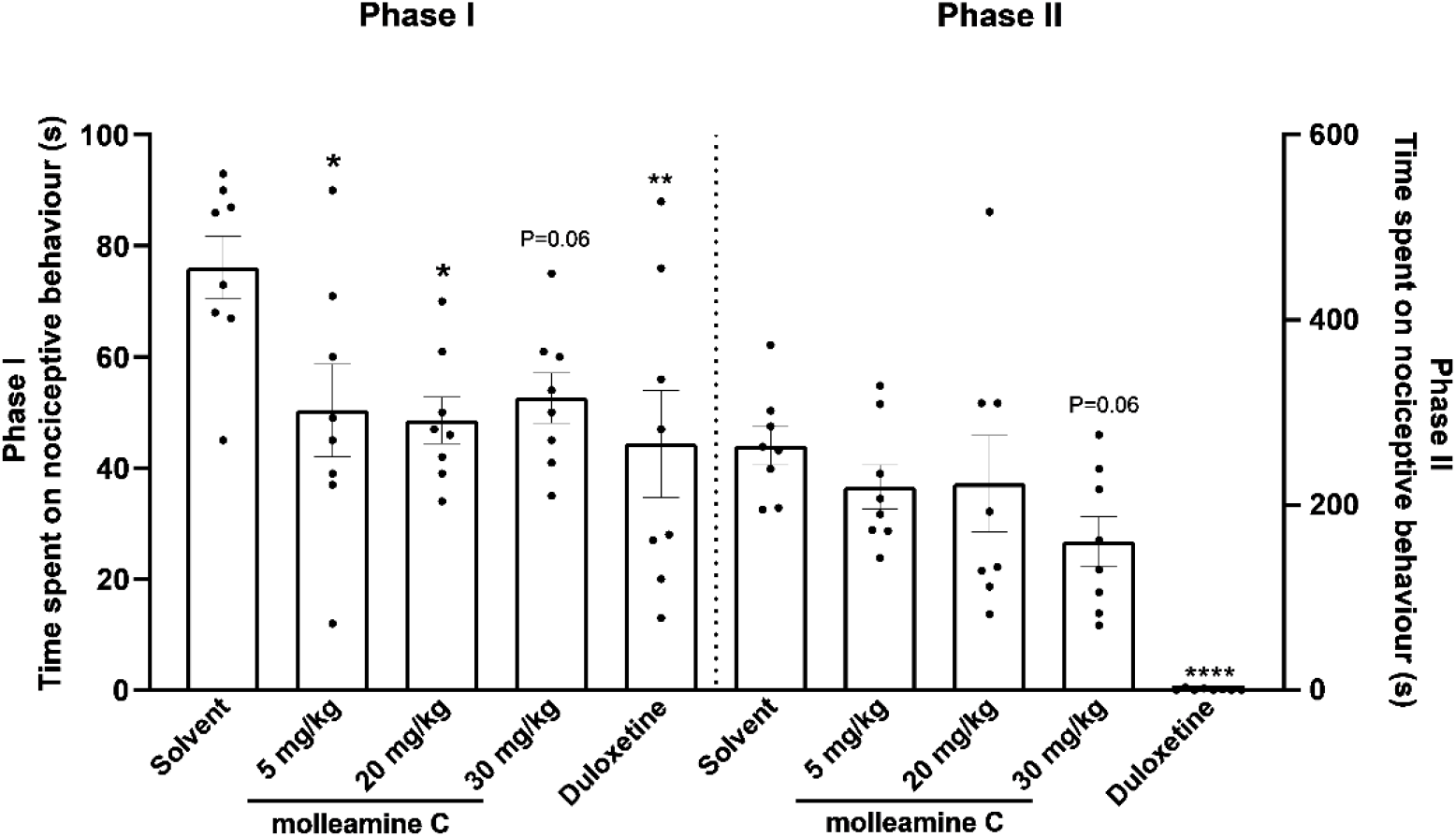
Compound **3** is antinociceptive in the formalin test. Male adult mice (n = 8 per group) were pretreated with **3** (5 mg/kg, 20 mg/kg, 30 mg/kg, i.p), vehicle (5 mL/kg, i.p.) or duloxetine (10 mg/kg, 10 mL/kg, i.p.). Formalin (5% in saline) was injected into the left hind paw pad after dosing; nociceptive behaviors were recorded for 0-35 min. Phase I represents the sum of time spent on nociceptive behavior in the first 5 min immediately after formalin injection while Phase II denotes the response period 20-35 min after injection. Each animal is represented by a circle, and the average SEM is shown with error bars. Significance in comparison to vehicle: \**p*<0.05, \*\**p*<0.01, \*\*\*\**p*<0.0001. Compound **3** inhibited nociceptive behaviors in Phase I at doses 5 mg/kg (*p*<0.05), 20 mg/kg (*p*<0.05), and 30 mg/kg (*p* = 0.06) and in Phase II at 30 mg/kg (*p* = 0.06). **Figure supplement 1**. Stability of compound **3** in mouse plasma over 24 h. **Figure supplement 2**. Cytotoxicity evaluation of compound **3** in *in vitro* MTT assay with human embryonic kidney (HEK-293) cells. **Figure supplement 3**. Zebrafish photomotor-response after exposure to increasing concentrations of compound **3**.

Stability of **3** was monitored in mouse blood plasma over 24 hours. No degradation was detected within an hour, but after 16 hours significant degradation of **3** (>50%) was observed (**Figure 7-figure supplement 1**). These data suggest that the majority of **3** remained intact over the course of the formalin experiment. No acute toxicity was observed at the chosen highest dose. However, mortality was observed in mice at doses above 45 mg/kg. The compound was found to have a cytotoxic effect in human embryonic kidney (HEK-293) cells with IC_50_ = 54 µM (**Figure 7-figure supplement 2**). Moreover, **3** did not cause toxicity to juvenile zebrafish at doses up to 100 µM, although a hyperactive phenotype was observed at doses as low as 30 µM (**Figure 7-figure supplement 3**).

## DISCUSSION

Our previous work with DRG neurons led to the identification of several families of active marine natural products that we have investigated for antinociceptive efficacy *(Lin, et al., 2010; Lin, et al., 2011; Lin, et al., 2017)*. In the course of this work, we identified several hits that seemed to target only a small subset of cells, but we had no framework to identify those cells, leading us to develop constellation pharmacology. Here, we show the power of constellation pharmacology to rapidly identify cell-type selective agents with therapeutic promise. Although Aδ-LTMRs constituted < 2% of DRG neurons, we could immediately assess the selective impact of **3**-**5** in blocking ATP-triggered, P2Y1-based depolarization of that neuronal subclass. Although ACh-responsive cells were <4% of the DRG neurons, selective block of ACh in peptidergic nociceptors was immediately apparent. Because both neuronal subclasses are important in pain conditions, we are currently assessing the potential of **3** as a neuroactive agent in analgesia and anesthesia.

The effects of **3** as an α3β4 and α6/α3β4 nAChR partial antagonist were clearly elucidated using a combination of electrophysiology and constellation pharmacology. Where those receptors were dominant in neurons, almost complete block of calcium flux could be observed, but when those receptors were absent or other receptors were dominant partial or no block was seen. Although there are countless nAChR-acting drugs and ligands, there are few close analogs of **3** in the literature in terms of biological activity. For example, nAChR agonist AT-1001 has been extensively studied *(Toll, et al., 2012; Yuan, et al., 2017)*. Although it was initially described as a selective α3β4 partial antagonist *(Toll, et al., 2012)*, more recent data reveal that it is a weak partial agonist that is competitive with ACh, while our data definitively rule out that possibility for **3**. Its physiological actions on animals are quite different than what we have observed for **3**. There is also a growing list of marine natural products targeting nAChRs, including compounds from algae, tunicates, sponges, mollusks, dinoflagellates, bryozoans, and corals *(Aráoz, et al., 2015; Culver, et al., 1984; Hamouda, et al., 2015; Kasheverov, et al., 2015; Kudryavtsev, et al., 2014; Tsuneki, et al., 2005; Wonnacott and Gallagher, 2006)*. However, none of these agents exhibit the selectivity shown by **3**. Most of them target neuromuscular nAChRs primarily and thus cause acute toxicity in vertebrates, whereas **3** only showed lethality in mice at doses greatly elevated above the α3β4 and α6/α3β4 nAChR partial antagonist activities, and no evidence of lethality in fish despite provoking a strong hyperactive response even at low doses. These results demonstrate that **3** has a different set of biological actions than previously characterized nAChR-targeted small molecules from the ocean.

In this study, we show that previously unknown compounds molleamines are widely distributed in *D. molle* tunicates, and that the most potent compounds are likely concentrated in the diet of *P. forskalii*, where they may serve as defensive metabolites. Our ability to focus on neuronal cell-type selectivity led to identification of molleamines as novel neuroactive compounds. The chemical simplicity of these polymeric compounds, as well as our development of a robust synthesis, makes them accessible for future structure-activity relationship studies to assess the pharmaceutical potential of this new chemical class of neuroactive compounds more fully.

## METHODS

### General experimental procedures

The UV data were acquired using a Thermo Scientific Evolution 201 UV-VIS Spectrophotometer. IR spectra were recorded using a Nicolet iS50 FT-IR spectrometer. NMR data were collected using either a Varian INOVA 500 spectrometer operating at 500 MHz for ^1^H and 125 MHz for ^13^C, and equipped with a 3mm Nalorac MDBG probe with a *z*-axis gradient; or a Varian INOVA 600 spectrometer operating at 600 MHz for ^1^H and 150 MHz for ^13^C NMR, and equipped with a 5 mm ^1^H[^13^C,^15^N] triple resonance cold probe with a *z*-axis gradient. NMR shift values were referenced to the residual solvent signals (DMSO-*d*_*6*_: δ_H_ 2.50, δ_C_ 39.5). UPLC-qTOFMS-MS/MS analysis of the compounds were performed on a Waters Acquity UPLC system coupled to a Waters Xevo G2-XS qTOF equipped with an ESI source. HPLC separations were performed using a Thermo Scientific Dionex WPS-3000 HPLC system equipped with a Photodiode array detector. Unless stated otherwise, all reagents and solvents were purchased from commercial suppliers and were used without further purification.

### Biological material and gene sequencing

*P. forskalii* was collected by hand using SCUBA in April 2018 (specimen SI-223L) in Solomon Islands (S 09° 22.891’ E 159° 52.428’). The freshly collected sample was kept frozen at -20 °C until use. The DNA was extracted from a small portion of the sample (∼25 mg) using the Qiagen DNeasy kit (Qiagen, Germantown, MD). The mitochondrial COXI genes were amplified using Folmer’s universal COXI primers LCO1490 (5’-GGT CAA CAA ATC ATA AAG ATA TTG G) and HCO2198 (5’-TAA ACT TCA GGG TGA CCA AAA AAT CA) *(Folmer, et al., 1994)*. The polymerase chain reaction was performed using a master mix consisting of 38 µL H_2_O, 5 µL 10× PCR buffer (High Fidelity Buffer; Invitrogen, Waltham, MA), 1.5 µL 50 mM MgCl_2_, 1 µL 10 mM LCO1490 primer, 1 µL 10 mM HCO2198 primer, 1 µL 10 mM dNTP mix, 0.5 µL 5U/µL Platinum Taq DNA Polymerase High Fidelity (Invitrogen) and 2 µL 0.25 ng/µL template DNA. PCR conditions were as follows: hot start (94 °C/2 min), 39 cycles of [94 °C/30 s, 45 °C/30 s, 72 °C/2 min], and extension (72 °C/10 min). The PCR product was gel purified using the Qiagen QIAquick kit and Sanger sequenced (Genewiz, Boston, MA). The COX1 gene sequence was submitted to GenBank (accession number MW663488).

### Extraction and isolation

The frozen sample (50 g wet weight) was thawed, diced, and exhaustively extracted with ethanol. The extract was dried *in vacuo* and partitioned between H_2_O (100 mL) and CHCl_3_ (100 mL × 3). The CHCl_3_-soluble extract was purified using reversed-phase HPLC with a Phenomenex Luna C_18_ column (250 × 10 mm) and a linear gradient from 20% to 100% CH_3_CN in H_2_O (0.1% TFA) over 20 min at 3mL/min flow rate. Resulting fractions were further purified using reversed-phase HPLC with a Phenomenex Luna C_18_ column (250 × 4 mm) with a linear gradient from 20% to 50% CH_3_CN in H_2_O (0.1% TFA) over 40 min at 1 mL/min flow rate to give compounds **1** (1.5 mg), **3** (4.0 mg), **4** (1.8 mg) and **5** (0.7 mg). One further fraction was purified using the same C_18_ column with a linear gradient from 5% to 100% CH_3_CN in H_2_O (0.1% TFA) over 40 min at 1 mL/min to yield compound **2** (0.6 mg). Overall yield: 0.017% of wet weight.

#### *Molleamine A* (**1**)

colorless amorphous solid; UV (CH_3_OH) λ_max_ (log ε) 207 (4.2) nm; IR ν_max_ 3290, 3061, 2919, 2849, 1678, 1532, 1445, 1207, 1138 cm^-1^; ^1^H and ^13^C NMR, Table 1; HRESIMS *m/z* 269.1656 [M+H]^+^ (calcd for C_17_H_21_N_2_O^+^, 269.1649).

#### *Molleamine B* (**2**)

colorless amorphous solid; UV (CH_3_OH) λ_max_ (log ε) 206 (3.6) nm; IR ν_max_ 3280, 3061, 2919, 2848, 1678, 1537, 1445, 1207, 1140 cm^-11^H and ^13^C NMR, Table 1; HRESIMS *m/z* 285.1603 [M+H]^+^ (calcd for C_17_H_21_N_2_O_2_^+^, 285.1598).

#### *Molleamine C* (**3**)

colorless amorphous solid; UV (CH_3_CN) λ_max_ (log ε) 205 (4.7) nm; IR ν_max_ 3272, 3063, 2910, 2846, 1682, 1538, 1446, 1206, 1139 cm^-11^H and ^13^C NMR, Table 1; HRESIMS *m/z* 416.2345 [M+H]^+^ (calcd for C_26_H_30_N_3_O_2_^+^, 416.2333).

#### *Molleamine D* (**4**)

colorless amorphous solid; UV (CH_3_CN) λ_max_ (log ε) 206 (3.9) nm; IR ν_max_ 3305, 2918, 2849, 1682, 1539, 1447, 1211, 1143 cm^-1 1^H and ^13^C NMR, Table 1; HRESIMS *m/z* 563.3022 [M+H]^+^ (calcd for C_35_H_39_N_4_O_2_^+^, 563.3017).

#### *Molleamine E* (**5**)

colorless amorphous solid; UV (CH_3_CN) λ_max_ 209 nm; IR ν_max_ 3308, 2918, 2846, 1682, 1580, 1446, 1210, 1143 cm^-1 1^H and ^13^C NMR, Table 1; HRESIMS *m/z* 710.3719 [M+H]^+^ (calcd for C_44_H_48_N_5_O_2_^+^, 710.3701).

### Synthesis of molleamine A (1)

To an ice-cold solution of 2-(2-aminoethyl)benzoic acid (190 mg, 0.94 mmol) dissolved in 10% Na_2_CO_3_ (10 mL) was added *N*-(9-fluorenylmethoxycarbonyloxy)succinimide (250 mg, 0.75 mmol) in acetone (10 mL). The mixture was stirred overnight at rt. The solution was dried *in vacuo* to remove the acetone, and the remaining aqueous portion was acidified to pH=2 with addition of 6N HCl and stirred for 1 h. The precipitate obtained was repeatedly washed with deionized H_2_O and air dried. To the reaction product (240 mg) dissolved in dichloromethane (20 mL, cooled to 0 °C) was added in order HOBt (120 mg, 0.91 mmol), DIEA (120 mg, 0.91 mmol), phenethylamine (88 mg, 0.73 mmol), and a solution of EDC·HCl (180 mg, 0.91 mmol) in dichloromethane (20 mL). The mixture was stirred overnight at rt. Water (20 mL) was then added, and then the dichloromethane layer was collected, dried with Na_2_SO_4_, and concentrated *in vacuo*. The resulting residue was stirred in 20% piperidine in DMF (5 mL) for 5 min, dried, and purified via reversed-phase HPLC using Phenomenex Luna C_18_ column (250 × 10 mm) with a linear gradient elution from 20% to 25% CH_3_CN in H_2_O (0.1% TFA) over 20 min at 3.5 mL/min to yield **1** (117 mg, 46% yield): ^1^H NMR (500 MHz, DMSO-*d*_*6*_) δ_H_ 8.56 (1H, t, *J* = 5.4 Hz), 8.03 (3H, brs), 7.42 (1H, m), 7.30-7.33 (5H, m), 7.26 (2H, d, *J* = 7.4 Hz), 7.22 (1H, t, *J* = 7.1 Hz), 3.49 (2H, td, *J* = 7.3, 5.4 Hz), 3.04 (2H, m), 2.93 (2H, m), 2.86 (2H, t, *J* = 7.3 Hz); ^13^C NMR (125 MHz, DMSO-*d*_*6*_) δ_C_ 168.9, 139.4, 137.0, 135.3, 130.3, 129.9, 128.7(×2), 128.4(×2), 127.4, 126.8, 126.2, 40.6, 40.2, 34.9, 31.0; HRESIMS *m/z* 269.1650 [M+H]^+^ calcd for C_17_H_21_N_2_O^+^, 269.1649). Chromatographic co-elution of the natural with the synthetic compound showed a uniform peak (**Figure 1-figure supplement 25**).

### Synthesis of molleamine C (3)

To an ice-cold solution of 2-(2-aminoethyl)benzoic acid (150 mg, 0.74 mmol) dissolved in 10% Na_2_CO_3_ (10 mL) was added *N*-(9-fluorenylmethoxycarbonyloxy)succinimide (209 mg, 0.62 mmol) in acetone (10 mL). The mixture was stirred overnight at rt. The solution was dried *in vacuo* to remove the acetone, and the remaining aqueous portion was acidified to pH=2 with addition of 6N HCl and stirred for 1 h. The precipitate obtained was repeatedly washed with deionized H_2_O and air dried. To the reaction product (48 mg) dissolved in dichloromethane (4 mL, cooled to 0 °C) was added in order HOBt (25 mg, 0.19 mmol), DIEA (24 mg, 0.19 mmol), molleamine A (40 mg, 0.15 mmol) in dichloromethane (1 mL), and a solution of EDC·HCl (36 mg, 0.19 mmol) in dichloromethane (5 mL). The mixture was stirred overnight at rt. Water (10 mL) was then added, and the dichloromethane layer was collected, dried with Na_2_SO_4_, and concentrated *in vacuo*. The resulting residue was stirred in 20% piperidine in DMF (1 mL) for 5 min, dried, and purified via reversed-phase HPLC using *Phenomenex* Luna C_18_ column (250 × 10 mm) with an isocratic elution at 32% CH_3_CN in H_2_O (0.1% TFA) for 17 min at 3.5 mL/min to yield **3** (43 mg, 69% yield). The residue obtained was then subjected to reversed-phase HPLC to yield **3**: ^1^H NMR (500 MHz, DMSO-*d*_*6*_) δ_H_ 8.67 (1H, t, *J* = 5.2 Hz), 8.48 (1H, t, *J* = 5.6 Hz), 8.01 (3H, brs), 7.41 (1H, m), 7.40 (1H, m), 7.35-7.26 (9H, m), 7.21 (1H, t, *J* = 7.1 Hz), 3.49 (4H, m), 3.04 (2H, m), 2.93 (2H, t, *J* = 7.3 Hz), 2.92 (2H, t, *J* = 7.1 Hz), 2.86 (2H, t, *J* = 7.3 Hz) ; ^13^C NMR (125 MHz, DMSO-*d*_*6*_) δ_C_ 169.4, 169.0, 139.5, 137.3(×2), 136.9, 135.6, 130.5, 130.2, 130.0, 129.5, 128.8(×2), 128.4(×2), 127.5, 127.2, 126.8, 126.2(×2), 41.0, 40.6, 40.3, 35.0, 32.1, 31.1; HRESIMS m/z 416.2332 [M+H]^+^ calcd for C_26_H_30_N_3_O_2_^+^, 416.2333). Chromatographic co-elution of the natural with the synthetic compound showed a uniform peak (**Figure 1-figure supplement 29**).

### Metabolomics analysis

UPLC-MS and MS/MS analyses of 21 tunicate and single mollusk specimens were done using an Agilent 6530 Q-TOF mass spectrometer with a Kinetex C_18_ column (2.6 µ, 100 A, 100 × 4.6 mm, 1 mL/min) and a gradient from 5 to 100 % MeCN in 20 min. The raw LC-MS/MS data were converted to mgf format using MassHunter. The mgf version of the data was then submitted to molecular networking analysis using the GNPS web site *(Wang, et al., 2016)* with the standard parameter and MSCluster option turned off. The output result was visualized using Cytoscape v3.7 *(Shannon, et al., 2003)*.

### Animals

All experiments involving the care and use of animals were conducted in accordance with ethical guidelines that were approved by the Institutional Animal Care and Use Committee (IACUC) of the University of Utah, Charles River Laboratories-Montreal IACUC, and the USA National Research Council and the Canadian Council on Animal Care (CCAC). In the formalin test male C57BL/6 mice (20-30 g, 6-8 weeks old, Charles River Laboratories, Canada) were acclimated for 5 days in the laboratory environment before the start of treatment.

### Cell culture and calcium imaging of cultured DRG and SCG cells

Descriptions of DRG and SCG cell preparation and calcium imaging protocols have been described in detail previously *(Hone, et al., 2020; Jackson and Tourtellotte, 2014; Light, et al., 2008; Memon, et al., 2017; Memon, et al., 2019)*. Briefly, lumbar DRG neurons were harvested from a CGRP-green fluorescent protein (GFP) mouse in a CD-1 genetic background. In these transgenic mice, PNs in the sensory neuronal population can be tracked through GFP expression. DRG neurons were dissociated by trypsinization and mechanical trituration and were subsequently plated onto a 24-well poly-D-lysine-coated plate. DRG neurons were cultured and incubated overnight at 37 °C, in a 5% CO_2_ tissue culture incubator and with 0.7 mL minimum essential medium (MEM, pH = 7.4) supplemented with 10% fetal bovine serum, penicillin (100 U/mL), streptomycin (100 µg/mL), 10 mM HEPES, and 0.4% (w/v) glucose.

The cultured cells were incubated with Fura-2-acetoxymethyl ester (Fura-2-AM; Molecular Probes; 2.5 µM) in MEM (0.7 mL) at 37 °C for 1 h and equilibrated at room temperature 0.5 hour prior to imaging. The MEM solution was then removed and the cells were washed three times (0.7 mL each) with observation solution (145 mM NaCl, 5mM KCl, 2mM CaCl_2_, 1 mM MgCl_2_, 1 mM sodium citrate, 10 mM HEPES, and 10 mM glucose). Experiments were performed at rt using fluorescence microscopy. In each calcium imaging experiment, >1000 cells were imaged simultaneously with individual cells treated as individual samples and their individual responses analyzed. Changes in intracellular calcium level ([Ca^2+^]_i_) while applying different pharmacological agents were measured by taking the relative ratio of emissions at 510 nm resulting from the excitation of the Fura-2-AM dye at 340 and 380 nm.

In general, calcium transients were elicited by a ∼15-s application of a pharmacological agent/depolarizing stimulus as follows: the observation solution was aspirated from the well via a peristaltic pump controlled by a microfluidic system, and then a 700 μL solution of a depolarizing stimulus was added (either via pipet or the microfluidics system). After ∼15-s incubation, the solution was replaced with observation solution in the same manner. The observation solution was typically replaced three more times over the next ∼50 s. This washing procedure was repeated as needed at intervals ranging from 3 to 8 min.

Each DRG experiment was followed by sequential application of pharmacological agents to identify neuronal cell classes. Pharmacological identifiers present in each experiment included KCl (20 mM), κM-RIIIJ (1 µM), AITC (100 µM), menthol (400 µM), and capsaicin (300 nM). At the end of the experiment, KCl (40 mM) was applied to assess viability of neurons. All solutions used in the experiment were prepared with DRG observation solution. Test compounds were dissolved in DMSO and diluted to the desired concentrations with DRG observation solution (DMSO final concentration was kept at no more than 0.1 % (v/v)).

After each DRG calcium imaging experiment, the cells were incubated with 0.7 mL Hoechst stain (1000 µg/mL) for 5 min and then washed 3x with the observation solution. After this, the cells were incubated with 0.7 mL Alexa-Fluor 647 Isolectin (2.5 µg/mL) for 5 min and then washed 3× with the observation solution. Finally, a bright-field image, a GFP image to visualize CGRP positive cells, and a Cy5 image to identify IB4-positive cells, were acquired using a rhodamine filter set. Nis elements and CellProfiler *(Jones, et al., 2008)* were used to acquire and create ROIs and to extract cellular information, respectively. Video information and trace data were extracted using in-house software built in Python and R language. Responses to the pharmacological identifiers and characteristic IB4 and CGRP labeling were used to group cells into different subclasses *(Bosse, et al., 2021; Giacobassi, et al., 2020)*.

Work with SCG cells was performed similarly, except that pharmacological differentiating agents were not used to classify the cell types. Our functional data shows that neurons in SCG are predominantly cholinergic, which is consistent with previous literature reports.

### Functional classification of DRG neuronal subclasses

Cell types are differentiated into 16 discrete types as follows. Cells with a cross-sectional area >500 µm^2^ are classified as large (L), and cells with <500 µm^2^ cross-sectional area are divided into those that are CGRP-GFP positive (G), IB4 positive (R), or negative to both (N). L cells that are CGRP-GFP positive are further divided into those that do (L5) and do not (L6) respond to conotoxin RIIIJ. L cells that are CGRP-GFP negative are divided in those that do not (L4) or do respond to conotoxin RIIIJ; the latter are further divided into those without a direct effect to RIIIJ (L3) and those that do have a direct effect (L1 and L2); the shape of the direct effect differentiates L1 (proprioceptors) and L2 (Aδ-LTMRs), as previously reported *(Giacobassi, et al., 2020)*.

G cells are divided by their response to menthol: positive (G7), or negative; the negatives are further divided by whether they are capsaicin responsive (G8; peptidergic nociceptors), AITC responsive (G10), or respond to both (G9).

R cells are differentiated based upon their response to capsaicin (R11), AITC (R13, nonpeptidergic nociceptors), or both (R12). N cells are recognized based upon their response to capsaicin (N16), menthol (N15, thermosensors), or neither (N14, C-low threshold mechanoreceptors).

### Dose-dependent effect of 3 on ATP-induced depolarization in DRG neurons

The cells were depolarized by two ∼15-s applications of ATP (20 µM). After the second depolarization, the cells were incubated for 8 min with 6.7 µM, 20 µM, 60 µM, and 180 µM of **3** at times 13, 23, 33, and 43 min, respectively. Each incubation with **3** was followed by application of ATP to determine the effect of **3** on the responses of the neurons to depolarization. A final application of ATP at 61 min was done to determine the reversibility of the responses of the cells to depolarization in the presence of **3**. This experiment was repeated twice (36-day old male and female mice) with a total of about 3,600 DRG neurons.

### Dose-dependent effect of 3 on MRS-2365-induced depolarization in DRG neurons

The cells were depolarized by two ∼15-s applications of MRS-2365 (100 nM). After the second depolarization, the cells were incubated for 8 min with 6.7 µM, 20 µM, 60 µM, and 180 µM of **3** at times 13, 23, 33, and 43 min, respectively. Each incubation with **3** was followed by ∼15s application of MRS-2365 to determine the effects of **3** on the responses of the neurons to depolarization. A final application of MRS-2365 at 61 min was done to determine the reversibility of the responses of the cells to depolarization in the presence of **3**. This experiment was repeated twice (36-day old male and female mice) with a total of about 3,100 DRG neurons.

### Constellation pharmacology of 3 on acetylcholine-induced depolarization in DRG neurons

A ∼15-s application of 30 mM KCl was done at the start of the calcium imaging experiment. This was followed by two consecutive ∼15-s applications of acetylcholine (ACh, 1 mM). After the second application of ACh, the cells were incubated for 4 min with 0.2 µM, 2 µM, and 20 µM of **3** at times 15, 21, and 27 min, respectively and with 500 nM atropine and 1 μM TxID at times 39 and 51 min, respectively. Each incubation with the test compounds was preceded and followed by a ∼15-s application of ACh. This experiment was done twice on a 34-day old male mouse (2 technical replicates; 1 biological replicate) with a total of 2,400 DRG neurons and 5000 glia.

### Constellation pharmacology of compounds 1-5 with ATP-induced depolarization

The ∼15-s application of 20 μM ATP was used to induce depolarization in the cells. The cells were incubated for 8 min twice with 20 μM of the test compound at times 13 and 33 min. Compounds **1**-**5** were tested in different wells and repeated in two calcium imaging experiments on 29 to 34-day old male and female mice with at least 1,000 DRG neurons per well.

### Constellation pharmacology of compounds 1-5 with KCl-induced depolarization

A ∼15-s application 30 mM KCl was used to induce depolarization in the cells. The cells were incubated for 4 min twice with 10 μM of the test compound at times 9 and 27 min. Compounds **1**-**5** were tested in different wells, and the experiment was done once for each compound on 42 to 44-day old male and female mice with at least 1,000 cells per well.

### Constellation pharmacology of 3 targeting the α7 nAChR

The cells were depolarized by two ∼15-s applications of ACh (1 mM). After the second application of ACh, the cells were pre-incubated for 4 min with 1 μM PNU 120596 at times 9, 15, 33, 45, 51 min followed by a ∼15-s co-application of ACh and PNU 120596 at times 13, 19, 25, 31, 37, 43, 49, and 55 min. **3** (20 μM) co-applied with PNU 120596 (1 μM) was incubated with the cells for 4 min at times 21 and 27 min. The α-conotoxin ArIB[V11;V16D] (200 nM) co-applied with PNU 120596 (1 μM) was incubated with the cells for 4 min at 39 min. This experiment was repeated twice (31-day old male mouse) with a total of about 2,000 DRG neurons.

### Constellation pharmacology of 3 on acetylcholine-induced depolarization in SCG neurons

A ∼15-s application of 20 mM KCl was done at the start of the calcium imaging experiment. This was followed by two consecutive ∼15-s applications of acetylcholine (ACh, 300 µM). After the second application of ACh, the cells were incubated for 4 min with 0.2 µM, 2 µM, and 20 µM of **3** at times 15, 21, and 27 min, respectively and with 1 μM TxID at times 45 min, respectively. Each incubation with the test compounds was preceded and followed by a ∼15-s application of ACh. This experiment was done twice on a 25- and 42-day old male mice with a total of about 1,000 neurons.

### Single-cell transcriptomic analysis

Experiments were performed as described previously *(Giacobassi, et al., 2020)*. Briefly, individual cells were selected based upon their pharmacological response in the DRG assay, and then picked up with a fire-polished glass pipette. The cells were lysed, and messenger ribonucleic acid (mRNA) was reverse transcribed to generate complementary DNA (cDNA), which then underwent whole-transcriptome amplification, all using the QIAseq FX Single Cell RNA library kit according to the manufacturer’s protocol (Qiagen). The amplified cDNA was used to construct a sequencing library for the Illumina NGS platform also using the QIAseq FX kit. The amplified cDNA was fragmented to 300 bp in size, end repaired, and ligated to adapters. A final cleanup was performed with Agencourt AMPure XP magnetic beads (Beckman Coulter Life Sciences, Indianapolis, IN). The cDNA library was submitted to the Huntsman Cancer Institute High Throughput Genomics Shared Resource for library control and sequencing. Sequencing data were analyzed using in-house R scripts.

### Calcium imaging with HEK-293 overexpressing GCaMP6s

Calcium imaging experiments with HEK-293 (ATCC) overexpressing GCaMP6s (an ultrasensitive fluorescent protein calcium sensor) were used to assay P2Y1 activation. HEK-GCaMP6s cells were grown in DMEM:F12 (Invitrogen) containing 5% FBS, 0.3 mg/ml G418, and 1× penicillin/streptomycin (Invitrogen). For calcium imaging experiments, HEK-GCaMP6s cells were subcultured into 1% gelatin-coated 96-well cell culture plates and grown to 80-90% confluence. Before imaging, the medium was replaced with LHC-9 medium (Invitrogen) containing 0.75 mM trypan red and, for antagonist treatment wells, the P2Y1 antagonist MRS2179 or molleamine C, and incubated for 30 min before assessing P2Y1 activation. The agonist MRS2365, used to activate P2Y1, was prepared in LHC-9 at 3× concentration and added to cells at 37 °C as previously described *(Deering-Rice, et al., 2018; Lamb, et al., 2017)*. Calcium flux was detected using a NOVOstar fluorescent plate reader (BMG Labtech).

### Formalin test

Male adult (6-8 weeks old; n = 8 per treatment group) C57BL/6 mice were placed in an observation chamber for approximately 10 min for habituation before the start of the test. Animals were pre-treated via intraperitoneal (i.p.) injection with compound **3** (5, 20, and 30 mg/kg) or vehicle (DMSO-polyethylene glycol (PEG) 400-phosphate buffered saline (PBS)/1:6:13, 5 mL/kg) 10 min prior to formalin injection. Duloxetine (10 mg/kg, 10 mL/kg, i.p.) was used as a positive control drug and was administered 30 min before formalin injection. The mice then received intraplantar subcutaneous injection of 5 % formalin (in 30 μL phosphate buffered saline solution) into the left hind paw and were placed immediately back in the observation chamber. Formalin-evoked spontaneous nociceptive behaviors, including flinching, shaking, biting and licking of the injected paw in the mice, were then recorded for 0-60 min using a commercial camcorder. Nociceptive behaviors in the mice were scored using the recorded video files and assessed in the following bins: 0-5 minutes from early phase (Phase I) and 20-35 minutes from late phase (Phase II). Animals were euthanized immediately at the end of the study. Significance of drug effect versus vehicle were analyzed by one-way ANOVA and Dunnet’s multiple comparisons test with Graphpad Prism 9.0.0 (Graphpad Software).

### *In vitro* stability of 3 in mouse plasma

Fresh whole blood was collected from mice via cardiac puncture and transferred into tubes pre-coated with 0.12 M EDTA. Plasma was isolated from the whole blood by centrifugation at 1,500 × *g* for 10 min at 4 °C and transferred into 1.5 mL microcentrifuge tubes. The plasma was preheated to 37 °C prior to the start of the study. The reactions were initiated by the addition of **3** dissolved in DMSO to 400 μL of preheated plasma to yield final concentrations of **3** at 10, 30, and 100 μM (final DMSO concentration = 0.5%). The experiments were performed in a dry bath incubator set at 37 °C, and the reaction for each concentration was conducted in triplicate. Samples (50 μL) were taken at 0, 1, 4, 16, 24 h, added to 200 μL MeOH, mixed by vortexing for ∼1 min and then centrifuged at 19,000 × *g* for 10 min at 4 °C. The clear supernatants were analyzed by UPLC-qTOF-MS as follows: an aliquot of the supernatant was diluted in methanol containing internal standard (40 μL final volume), and 2 μL were loaded onto an Acquity UPLC HSS T3 (1.8 μm, 2.1 mm × 100 mm) column. A linear gradient of 5%-100% CH_3_CN in H_2_O (0.1% formic acid) over 7 min at 0.3 mL/min was used to elute the samples. The relative abundance of **3** at different time points was calculated as normalized area under the curve with respect to internal standard leucine enkephalin.

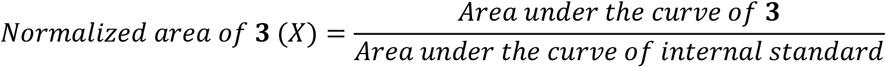

The percentage of **3** remaining in plasma at the individual time points relative to the 0 h sample was plotted versus incubation time (**Figure 7-figure supplement 1**). The approximate half-life for the compound was determined from the obtained graph, in which 50% of the compound remained.

### Oocyte receptor expression and electrophysiological recordings

Methods describing the preparation of cRNA encoding human, mouse, and rat nAChR subunits for expression of nAChRs in *X. laevis* oocytes have been described in detail previously *(Zheng, et al., 2020). X. laevis* oocytes were microinjected with cRNA encoding the selected nAChR subunits. Oocytes were incubated at 17 °C for 1–3 days in ND96 prior to use. Injected oocytes were placed in a 30 μL recording chamber and voltage clamped to a membrane potential of -70 mV. ND96 (96.0 mM NaCl, 2.0 mM KCl, 1.8 mM CaCl_2_, 1.0 mM MgCl_2_, 5 mM HEPES, pH 7.5) with 0.1 mg/mL BSA was gravity perfused through the recording chamber at ∼2 mL/min. A one second pulse of ACh was applied to measure the receptor response, with pulses occurring every minute. ACh was applied at a concentration of 100 μM for all subtypes. A baseline ACh response was established, and then the ND96 control solution was switched to a ND96 solution containing the various concentrations of compound **3** (0.67 µM, 2 µM, 6.7 µM, 20 µM). During perfusion of the compound-containing solutions, ACh pulses continued once per minute to assess for block of the ACh-induced response. ACh responses were measured in the presence of a compound concentration until the responses reached steady state; an average of three of these responses compared to the baseline response was used to determine percent response. To estimate the IC_50_ value for inhibition of the Ach responses by **3**, the normalized data were analyzed by nonlinear regression and fit using a four-parameter logistic equation in Graphpad Prism 9.0.0 (Graphpad Software).

### Zebrafish photomotor response assay

Zebrafish (*Danio rerio)* were obtained from the Centralized Zebrafish Animal Resource (CZAR) at the University of Utah. The zebrafish photomotor response assay to evaluate the effect of the compounds was performed as described previously with modifications *(Kokel, et al., 2010; Kokel and Peterson, 2011)*. Briefly, larvae were loaded onto a 96-well plate format at 168 hours post fertilization, 10 fish per well with 10 mM HEPES buffered E3 medium and transferred into a Zebrabox plate holder. Larval activity was tracked by using a Hamamatsu ORCA-ER camera mounted on a Nikon TE200 microscope with a 1× objective. A 300 W xenon bulb housed in a Sutter Lambda LS illuminator was used to deliver light stimuli and elicit PMR. The robotic stage, digital video camera, and stimulus presentation were all automated via the Metamorph Software (Universal Imaging). All experiments consisted of three minutes of acclimatization in white light (pre-white light) followed by sample addition. The larvae were treated with the compounds by spiking each well with the specified treatment and then thoroughly mixed with a pipet. After sample addition, this was followed by 3 min of incubation with white light (post-white light), then 7 min strobe light (in dark and white light), 3 min in the dark and 3 min white light and strobe light after 1 h. After 2 h, 7 min of strobe light, 3 min dark and 3 min white light.

### Mammalian cytotoxicity assay

HEK-293 (ATCC CRL-1573) cells were cultured in Dulbecco’s Modified Eagle Media (DMEM) supplemented with 10 % fetal bovine serum (FBS), 100 units of penicillin and 100 μg/mL of streptomycin under a humidified environment with 5% CO_2_, 95% air at 37°C. Cells were seeded in 96-well plates at a density of 10000 cells/well and treated after 24 h with varying concentrations of the test sample, positive control, and the solvent control. After 72h, the media was removed and 15μL of 5mg/mL MTT reagent was added to each well. This was then incubated for 3 h at 37°C, 5% CO_2_, and 95% air before addition of 100 μL DMSO. The absorbance was read at 570 nm using a microplate reader (Biotek Synergy HT). IC_50_ values were then calculated using GraphPad Prism 9.0.0 based on a four-point sigmoidal nonlinear regression analysis of cell viability vs log concentration of test sample.

## ACKNOWLEDGEMENTS

This work was funded by DOD W81XWH-17-1-0413, except for the tunicate metabolomic analysis, which was funded by NIH R35GM122521 to EWS, and electrophysiology experiments, which were funded by NIH R35GM136430 to JMM. Assistance from the Ministry of Environment, Climate Change, Disaster Management, and Meteorology (Solomon Islands) and Solomon Islands National University is gratefully acknowledged. We thank the ALSAM Foundation for supporting equipment acquisition.

## ADDITIONAL INFORMATION

### Funding

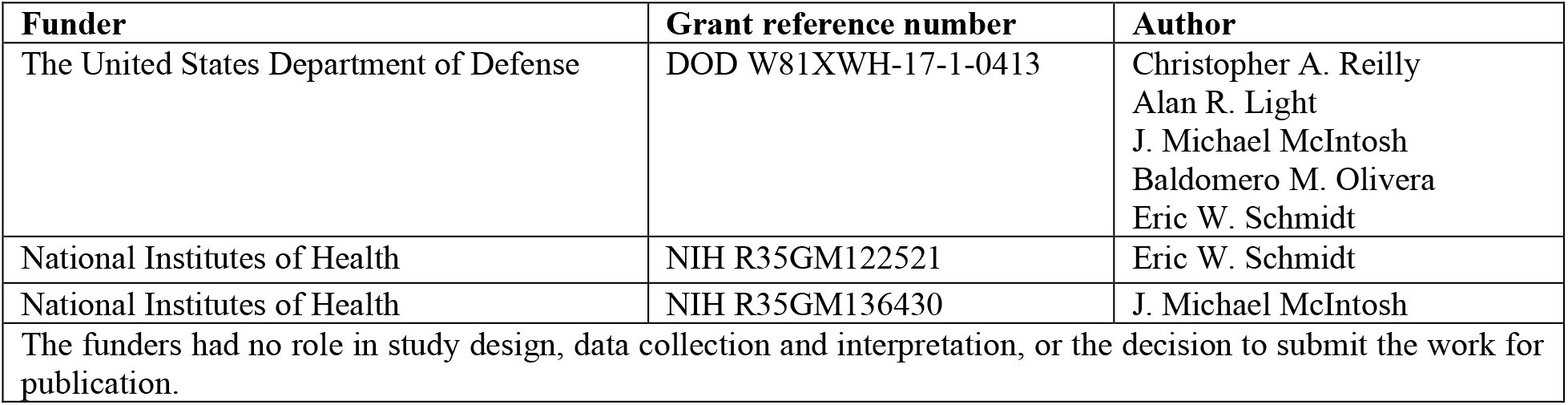

### Author Contributions

E.W.S. and B.M.O. designed the study; N.D.P., L.S.L, J.O.T., Z.L., K.C., C.D., C.E.D-R., A.L.L., M.K., R.W.H., J.Z. performed experiments and analyzed data. All authors contributed to manuscript preparation. The authors declare no conflicts of interest.

## ADDITIONAL FILES

Supplementary figures and tables.

## REFERENCES

Aráoz R, Ouanounou G, Iorga BI, Goudet A, Alili D, Amar M, Benoit E, Molgó J, & Servent D. 2015. The neurotoxic effect of 13,19-didesmethyl and 13-desmethyl spirolide C phycotoxins Is mainly mediated by nicotinic rather than muscarinic acetylcholine receptors. Toxicological Sciences 147: 156–167. DOI:10.1093/toxsci/kfv119

Bjørn-Yoshimoto WE, Ramiro IBL, Yandell M, McIntosh JM, Olivera BM, Ellgaard L, & Safavi-Hemami H. 2020. Curses or cures: A review of the numerous benefits versus the biosecurity concerns of conotoxin research. Biomedicines 8: 235. DOI:10.3390/biomedicines8080235

Bosse GD, Urcino C, Watkins M, Flórez Salcedo P, Kozel S, Chase K, Cabang A, Espino SS, Safavi-Hemami H, Raghuraman S, Olivera BM, Peterson RT, & Gajewiak J. 2021. Discovery of a potent conorfamide from Conus episcopatus using a novel zebrafish larvae assay. Journal of Natural Products. DOI:10.1021/acs.jnatprod.0c01297

Christensen SB, Hone AJ, Roux I, Kniazeff J, Pin J-P, Upert G, Servent D, Glowatzki E, & McIntosh JM. 2017. RgIA4 potently blocks mouse α9α10 nAChRs and provides long lasting protection against oxaliplatin-induced cold allodynia. Frontiers in Cellular Neuroscience 11: 219. DOI:10.3389/fncel.2017.00219

Cimino G, & Ghiselin MT. 2009. Chemical defense and the evolution of opisthobranch gastropods (Vol. 60). San Francisco, Ca: California Academy of Sciences.

Cucchiaro G, Xiao Y, Gonzalez-Sulser A, & Kellar Kenneth J. 2008. Analgesic effects of sazetidine-A, a new nicotinic cholinergic drug. Anesthesiology 109: 512–519. DOI:10.1097/ALN.0b013e3181834490

Culver P, Fenical W, & Taylor P. 1984. Lophotoxin irreversibly inactivates the nicotinic acetylcholine receptor by preferential association at one of the two primary agonist sites. Journal of Biological Chemistry 259: 3763–3770. DOI:10.1016/S0021-9258(17)43160-7

Deering-Rice CE, Nguyen N, Lu Z, Cox JE, Shapiro D, Romero EG, Mitchell VK, Burrell KL, Veranth JM, & Reilly CA. 2018. Activation of TRPV3 by wood smoke particles and roles in pneumotoxicity. Chemical Research in Toxicology 31: 291–301. DOI:10.1021/acs.chemrestox.7b00336

Dhandapani R, Arokiaraj CM, Taberner FJ, Pacifico P, Raja S, Nocchi L, Portulano C, Franciosa F, Maffei M, Hussain AF, de Castro Reis F, Reymond L, Perlas E, Garcovich S, Barth S, Johnsson K, Lechner SG, & Heppenstall PA. 2018. Control of mechanical pain hypersensitivity in mice through ligand-targeted photoablation of TrkB-positive sensory neurons. Nature Communications 9: 1640–1640. DOI:10.1038/s41467-018-04049-3

Folmer O, Black M, Hoeh W, Lutz R, & Vrijenhoek R. 1994. DNA primers for amplification of mitochondrial cytochrome c oxidase subunit I from diverse metazoan invertebrates. Molecular Marine Biology and Biotechnology 3: 294–299.

Fu K-Y, Light AR, & Maixner W. 2001. Long-lasting inflammation and long-term hyperalgesia after subcutaneous formalin injection into the rat hindpaw. The Journal of Pain 2: 2–11. DOI:10.1054/jpai.2001.9804

Giacobassi MJ, Leavitt LS, Raghuraman S, Alluri R, Chase K, Finol-Urdaneta RK, Terlau H, Teichert RW, & Olivera BM. 2020. An integrative approach to the facile functional classification of dorsal root ganglion neuronal subclasses. Proceedings of the National Academy of Sciences of the United States of America 117: 5494–5501. DOI:10.1073/pnas.1911382117

Hamouda AK, Wang Z-J, Stewart DS, Jain AD, Glennon RA, & Cohen JB. 2015. Desformylflustrabromine (dFBr) and [^3^H]dFBr-labeled binding sites in a nicotinic acetylcholine receptor. Molecular Pharmacology 88: 1–11. DOI:10.1124/mol.115.098913

Hirose M, Nozawa Y, & Hirose E. 2010. Genetic isolation among morphotypes in the photosymbiotic didemnid Didemnum molle (Ascidiacea, Tunicata) from the Ryukyus and Taiwan. Zoological Science 27: 959–964. DOI:10.2108/zsj.27.959

Hirose M, Yokobori S, & Hirose E. 2009. Potential speciation of morphotypes in the photosymbiotic ascidian Didemnum molle in the Ryukyu Archipelago, Japan. Coral Reefs 28: 119–126. DOI:10.1007/s00338-008-0425-0

Hone AJ, Meyer EL, McIntyre M, & McIntosh JM. 2012. Nicotinic acetylcholine receptors in dorsal root ganglion neurons include the α6β4* subtype. FASEB Journal : Official Publication of the Federation of American Societies for Experimental Biology 26: 917–926. DOI:10.1096/fj.11-195883

Hone AJ, Rueda-Ruzafa L, Gordon TJ, Gajewiak J, Christensen S, Dyhring T, Albillos A, & McIntosh JM. 2020. Expression of α3β2β4 nicotinic acetylcholine receptors by rat adrenal chromaffin cells determined using novel conopeptide antagonists. Journal of Neurochemistry 154: 158–176. DOI:10.1111/jnc.14966

Hone AJ, Talley TT, Bobango J, Huidobro Melo C, Hararah F, Gajewiak J, Christensen S, Harvey PJ, Craik DJ, & McIntosh JM. 2018. Molecular determinants of α-conotoxin potency for inhibition of human and rat α6β4 nicotinic acetylcholine receptors. Journal of Biological Chemistry 293: 17838–17852. DOI:10.1074/jbc.RA118.005649

Hunskaar S, & Hole K. 1987. The formalin test in mice: dissociation between inflammatory and non-inflammatory pain. Pain 30: 103–114. DOI:10.1016/0304-3959(87)90088-1

Issac M, Aknin M, Gauvin-Bialecki A, Pond CD, Barrows LR, Kashman Y, & Carmeli S. 2017. Mollecarbamates, molleureas, and molledihydroisoquinolone, o-carboxyphenethylamide metabolites of the ascidian Didemnum molle collected in Madagascar. Journal of Natural Products 80: 1844–1852. DOI:10.1021/acs.jnatprod.7b00123

Jackson M, & Tourtellotte W. 2014. Neuron culture from mouse superior cervical ganglion. Bio-protocol 4: e1035. DOI:10.21769/bioprotoc.1035

Jones TR, Kang IH, Wheeler DB, Lindquist RA, Papallo A, Sabatini DM, Golland P, & Carpenter AE. 2008. CellProfiler analyst: data exploration and analysis software for complex image-based screens. BMC Bioinformatics 9: 482. DOI:10.1186/1471-2105-9-482

Kasheverov IE, Shelukhina IV, Kudryavtsev DS, Makarieva TN, Spirova EN, Guzii AG, Stonik VA, & Tsetlin VI. 2015. 6-Bromohypaphorine from marine nudibranch mollusk Hermissenda crassicornis is an agonist of human alpha-7 nicotinic acetylcholine receptor. Marine Drugs 13: 1255–1266. DOI:10.3390/md13031255

Kokel D, Bryan J, Laggner C, White R, Cheung CYJ, Mateus R, Healey D, Kim S, Werdich AA, Haggarty SJ, MacRae CA, Shoichet B, & Peterson RT. 2010. Rapid behavior-based identification of neuroactive small molecules in the zebrafish. Nature Chemical Biology 6: 231–237. DOI:10.1038/nchembio.307

Kokel D, & Peterson RT. 2011. Using the zebrafish photomotor response for psychotropic drug screening. Methods in Cell Biology 105: 517–524. DOI:10.1016/B978-0-12-381320-6.00022-9

Kudryavtsev D, Makarieva T, Utkina N, Santalova E, Kryukova E, Methfessel C, Tsetlin V, Stonik V, & Kasheverov I. 2014. Marine natural products acting on the acetylcholine-binding protein and nicotinic receptors: from computer modeling to binding studies and electrophysiology. Marine Drugs 12: 1859–1875. DOI:10.3390/md12041859

Lamb JG, Romero EG, Lu Z, Marcus SK, Peterson HC, Veranth JM, Deering-Rice CE, & Reilly CA. 2017. Activation of human transient receptor potential melastatin-8 (TRPM8) by calcium-rich particulate materials and effects on human lung cells. Molecular Pharmacology 92: 653–664. DOI:10.1124/mol.117.109959

Light AR, Hughen RW, Zhang J, Rainier J, Liu Z, & Lee J. 2008. Dorsal root ganglion neurons innervating skeletal muscle respond to physiological combinations of protons, ATP, and lactate mediated by ASIC, P2X, and TRPV1. Journal of Neurophysiology 100: 1184–1201. DOI:10.1152/jn.01344.2007

Limapichat W, Dougherty DA, & Lester HA. 2014. Subtype-specific mechanisms for functional interaction between α6β4* nicotinic acetylcholine receptors and P2X receptors. Molecular Pharmacology 86: 263–274. DOI:10.1124/mol.114.093179

Lin Z, Antemano RR, Hughen RW, Tianero MDB, Peraud O, Haygood MG, Concepcion GP, Olivera BM, Light A, & Schmidt EW. 2010. Pulicatins A-E, neuroactive thiazoline metabolites from cone snail-associated bacteria. Journal of Natural Products 73: 1922–1926. DOI:10.1021/np100588c

Lin Z, Reilly CA, Antemano R, Hughen RW, Marett L, Concepcion GP, Haygood MG, Olivera BM, Light A, & Schmidt EW. 2011. Nobilamides A-H, long-acting transient receptor potential vanilloid-1 (TRPV1) antagonists from mollusk-associated bacteria. Journal of Medicinal Chemistry 54: 3746–3755. DOI:10.1021/jm101621u

Lin Z, Smith MD, Concepcion GP, Haygood MG, Olivera BM, Light A, & Schmidt EW. 2017. Modulating the serotonin receptor spectrum of pulicatin natural products. Journal of Natural Products 80: 2360–2370. DOI:10.1021/acs.jnatprod.7b00317

Loram LC, Taylor FR, Strand KA, Maier SF, Speake JD, Jordan KG, James JW, Wene SP, Pritchard RC, Green H, Van Dyke K, Mazarov A, Letchworth SR, & Watkins LR. 2012. Systemic administration of an alpha-7 nicotinic acetylcholine agonist reverses neuropathic pain in male Sprague Dawley rats. Journal of Pain 13: 1162–1171. DOI:10.1016/j.jpain.2012.08.009

Lu W, Reigada D, Sévigny J, & Mitchell CH. 2007. Stimulation of the P2Y1 receptor up-regulates nucleoside-triphosphate diphosphohydrolase-1 in human retinal pigment epithelial cells. Journal of Pharmacology and Experimental Therapeutics 323: 157–164. DOI:10.1124/jpet.107.124545

Lu Z, Harper MK, Pond CD, Barrows LR, Ireland CM, & Van Wagoner RM. 2012. Thiazoline peptides and a tris-phenethyl urea from Didemnum molle with anti-HIV activity. Journal of Natural Products 75: 1436–1440. DOI:10.1021/np300270p

Luo S, Zhangsun D, Zhu X, Wu Y, Hu Y, Christensen S, Harvey PJ, Akcan M, Craik DJ, & McIntosh JM. 2013. Characterization of a novel α-conotoxin TxID from Conus textile that potently blocks rat α3β4 nicotinic acetylcholine receptors. Journal of Medicinal Chemistry 56: 9655–9663. DOI:10.1021/jm401254c

McNamara CR, Mandel-Brehm J, Bautista DM, Siemens J, Deranian KL, Zhao M, Hayward NJ, Chong JA, Julius D, Moran MM, & Fanger CM. 2007. TRPA1 mediates formalin-induced pain. Proceedings of the National Academy of Sciences of the United States of America 104: 13525–13530. DOI:10.1073/pnas.0705924104

Memon T, Chase K, Leavitt LS, Olivera BM, & Teichert RW. 2017. TRPA1 expression levels and excitability brake by KV channels influence cold sensitivity of TRPA1-expressing neurons. Neuroscience 353: 76–86. DOI:10.1016/j.neuroscience.2017.04.001

Memon T, Yarishkin O, Reilly CA, Krizaj D, Olivera BM, & Teichert RW. 2019. Trans-anethole of fennel oil is a selective and non-electrophilic agonist of the TRPA1 ion channel. Molecular Pharmacology: 433–441. DOI:10.1124/mol.118.114561

Raghuraman S, Xie JY, Giacobassi MJ, Tun JO, Chase K, Lu D, Teichert RW, Porreca F, & Olivera BM. 2020. Chronicling changes in the somatosensory neurons after peripheral nerve injury. Proceedings of the National Academy of Sciences of the United States of America 117: 26414–26421. DOI:10.1073/pnas.1922618117

Romero HK, Christensen SB, Di Cesare Mannelli L, Gajewiak J, Ramachandra R, Elmslie KS, Vetter DE, Ghelardini C, Iadonato SP, Mercado JL, Olivera BM, & McIntosh JM. 2017. Inhibition of α9α10 nicotinic acetylcholine receptors prevents chemotherapy-induced neuropathic pain. Proceedings of the National Academy of Sciences of the United States of America 114: E1825–E1832. DOI:10.1073/pnas.1621433114

Shannon P, Markiel A, Ozier O, Baliga NS, Wang JT, Ramage D, Amin N, Schwikowski B, & Ideker T. 2003. Cytoscape: a software environment for integrated models of biomolecular interaction networks. Genome Research 13: 2498–2504. DOI:10.1101/gr.1239303

Simeone X, Karch R, Ciuraszkiewicz A, Orr-Urtreger A, Lemmens-Gruber R, Scholze P, & Huck S. 2019. The role of the nAChR subunits α5, β2, and β4 on synaptic transmission in the mouse superior cervical ganglion. Physiological reports 7: e14023–e14023. DOI:10.14814/phy2.14023

Smith NJ, Hone AJ, Memon T, Bossi S, Smith TE, McIntosh JM, Olivera BM, & Teichert RW. 2013. Comparative functional expression of nAChR subtypes in rodent DRG neurons. Frontiers in Cellular Neuroscience 7: 225–225. DOI:10.3389/fncel.2013.00225

Tan KC, Wakimoto T, Takada K, Ohtsuki T, Uchiyama N, Goda Y, & Abe I. 2013. Cycloforskamide, a cytotoxic macrocyclic peptide from the sea slug Pleurobranchus forskalii. Journal of Natural Products 76: 1388–1391. DOI:10.1021/np400404r

Teichert RW, Memon T, Aman JW, & Olivera BM. 2014. Using constellation pharmacology to define comprehensively a somatosensory neuronal subclass. Proceedings of the National Academy of Sciences of the United States of America 111: 2319–2324. DOI:10.1073/pnas.1324019111

Teichert RW, Schmidt EW, & Olivera BM. 2015. Constellation pharmacology: A new paradigm for drug discovery. Annual Review of Pharmacology and Toxicology 55: 573–589. DOI:10.1146/annurev-pharmtox-010814-124551

Teichert RW, Smith NJ, Raghuraman S, Yoshikami D, Light AR, & Olivera BM. 2012. Functional profiling of neurons through cellular neuropharmacology. Proceedings of the National Academy of Sciences of the United States of America 109: 1388–1395. DOI:10.1073/pnas.1118833109

Tjølsen A, Berge OG, Hunskaar S, Rosland JH, & Hole K. 1992. The formalin test: an evaluation of the method. Pain 51: 5–17. DOI:10.1016/0304-3959(92)90003-t

Toll L, Zaveri NT, Polgar WE, Jiang F, Khroyan TV, Zhou W, Xie XS, Stauber GB, Costello MR, & Leslie FM. 2012. AT-1001: a high affinity and selective α3β4 nicotinic acetylcholine receptor antagonist blocks nicotine self-administration in rats. Neuropsychopharmacology 37: 1367–1376. DOI:10.1038/npp.2011.322

Tsuneki H, You YR, Toyooka N, Sasaoka T, Nemoto H, Dani JA, & Kimura I. 2005. Marine alkaloids (-)-pictamine and (-)-lepadin B block neuronal nicotinic acetylcholine receptors. Biological and Pharmaceutical Bulletin 28: 611–614. DOI:10.1248/bpb.28.611

Volkow ND, & McLellan AT. 2016. Opioid abuse in chronic pain — Misconceptions and mitigation strategies. New England Journal of Medicine 374: 1253–1263. DOI:10.1056/NEJMra1507771

Wakimoto T, Tan KC, & Abe I. 2013. Ergot alkaloid from the sea slug Pleurobranchus forskalii. Toxicon 72: 1–4. DOI:10.1016/j.toxicon.2013.05.021

Wang M, Carver JJ, Phelan VV, Sanchez LM, Garg N, Peng Y, Nguyen DD, Watrous J, Kapono CA, Luzzatto-Knaan T, Porto C, Bouslimani A, Melnik AV, Meehan MJ, Liu W-T, Crüsemann M, Boudreau PD, Esquenazi E, Sandoval-Calderón M, Kersten RD, Pace LA, Quinn RA, Duncan KR, Hsu C-C, Floros DJ, Gavilan RG, Kleigrewe K, Northen T, Dutton RJ, Parrot D, Carlson EE, Aigle B, Michelsen CF, Jelsbak L, Sohlenkamp C, Pevzner P, Edlund A, McLean J, Piel J, Murphy BT, Gerwick L, Liaw C-C, Yang Y-L, Humpf H-U, Maansson M, Keyzers RA, Sims AC, Johnson AR, Sidebottom AM, Sedio BE, Klitgaard A, Larson CB, Boya P CA, Torres-Mendoza D, Gonzalez DJ, Silva DB, Marques LM, Demarque DP, Pociute E, O’Neill EC, Briand E, Helfrich EJN, Granatosky EA, Glukhov E, Ryffel F, Houson H, Mohimani H, Kharbush JJ, Zeng Y, Vorholt JA, Kurita KL, Charusanti P, McPhail KL, Nielsen KF, Vuong L, Elfeki M, Traxler MF, Engene N, Koyama N, Vining OB, Baric R, Silva RR, Mascuch SJ, Tomasi S, Jenkins S, Macherla V, Hoffman T, Agarwal V, Williams PG, Dai J, Neupane R, Gurr J, Rodríguez AMC, Lamsa A, Zhang C, Dorrestein K, Duggan BM, Almaliti J, Allard P-M, Phapale P, Nothias L-F, Alexandrov T, Litaudon M, Wolfender J-L, Kyle JE, Metz TO, Peryea T, Nguyen D-T, VanLeer D, Shinn P, Jadhav A, Müller R, Waters KM, Shi W, Liu X, Zhang L, Knight R, Jensen PR, Palsson BØ, Pogliano K, Linington RG, Gutiérrez M, Lopes NP, Gerwick WH, Moore BS, Dorrestein PC, & Bandeira N. 2016. Sharing and community curation of mass spectrometry data with Global Natural Products Social Molecular Networking. Nature Biotechnology 34: 828–837. DOI:10.1038/nbt.3597

Wesson KJ, & Hamann MT. 1996. Keenamide A, a bioactive cyclic peptide from the marine mollusk leurobranchus forskalii. Journal of Natural Products 59: 629–631. DOI:10.1021/np960153t

Wonnacott S, & Gallagher T. 2006. The chemistry and pharmacology of anatoxin-a and related homotropanes with respect to nicotinic acetylcholine receptors. Marine Drugs 4: 228–254. DOI:10.3390/md403228

Wood SA, Taylor DI, McNabb P, Walker J, Adamson J, & Cary SC. 2012. Tetrodotoxin concentrations in Pleurobranchaea maculata: Temporal, spatial and individual variability from New Zealand populations. Marine Drugs 10: 163–176. DOI:10.3390/md10010163

Yuan M, Malagon AM, Yasuda D, Belluzzi JD, Leslie FM, & Zaveri NT. 2017. The α3β4 nAChR partial agonist AT-1001 attenuates stress-induced reinstatement of nicotine seeking in a rat model of relapse and induces minimal withdrawal in dependent rats. Behavioural Brain Research 333: 251–257. DOI:10.1016/j.bbr.2017.07.004

Zheng N, Christensen SB, Blakely A, Dowell C, Purushottam L, McIntosh JM, & Chou DH-C. 2020. Development of conformationally constrained α-RgIA analogues as stable peptide antagonists of human α9α10 nicotinic acetylcholine receptors. Journal of Medicinal Chemistry 63: 8380–8387. DOI:10.1021/acs.jmedchem.0c00613

Zheng Y, Liu P, Bai L, Trimmer JS, Bean BP, & Ginty DD. 2019. Deep sequencing of somatosensory neurons reveals molecular determinants of intrinsic physiological properties. Neuron 103: 598-616.e597. DOI:10.1016/j.neuron.2019.05.039

Zwart R, & Vijverberg HPM. 1997. Potentiation and inhibition of neuronal nicotinic receptors by atropine: Competitive and noncompetitive effects. Molecular Pharmacology 52: 886–895. DOI:10.1124/mol.52.5.886

