## Supporting Information for "Nicotinic acetylcholine receptor partial antagonist polyamides from tunicates and their predatory sea slugs"

[1]Department of Medicinal Chemistry, University of Utah, Salt Lake City, Utah, 81112, USA; [2]School of Biological Sciences, University of Utah, Salt Lake City, Utah 84112, USA; [3]Department of Pharmacology and Toxicology, University of Utah, Salt Lake City, Utah, 81112, USA; [4]Department of Anesthesiology, School of Medicine, University of Utah, Salt Lake City, Utah 84112, USA; [5]Department of Psychiatry, University of Utah, Salt Lake City, Utah 84112, USA; [6]George E Whalen Veterans Affairs Medical Center, Salt Lake City, Utah 84148, USA

**Figure 1-figure supplement 1**. Structures of compounds **1-5** with COSY and HMBC correlations.
**Figure 1-figure supplement 2**. $^{1}$H NMR (500 MHz; top) and $^{13}$C NMR (125 MHz; bottom) spectra of **1** in DMSO-$d_6$.
**Figure 1-figure supplement 3**. gCOSY (top) and gHSQC (bottom) spectra of **1** in DMSO-d6.
**Figure 1-figure supplement 4**. gHMBC spectrum of **1** in DMSO-d6
**Figure 1-figure supplement 5**. (+)-HRESITOFMS of compound **1** (top) with predicted elemental composition.
**Figure 1-figure supplement 6**.$^{1}$H NMR (500 MHz; top) and $^{13}$C NMR (125 MHz; bottom) spectra of **2** in DMSO-$d_6$.
**Figure 1-figure supplement 7**. gCOSY (top) and gHSQC (bottom) spectra of **2** in DMSO-$d_6$.
**Figure 1-figure supplement 8**. gHMBC spectrum of **2** taken in DMSO-$d_6$.
**Figure 1-figure supplement 9**. (+)-HRESITOFMS of compound **2** (top) with predicted elemental composition.
**Figure 1-figure supplement 10**.$^{1}$H NMR (500 MHz; top) and $^{13}$C NMR (125 MHz; bottom) spectra of **3** in DMSO-$d_6$.
**Figure 1-figure supplement 11**. gCOSY (top) and gHSQC (bottom) spectra of **3** in DMSO-$d_6$.
**Figure 1-figure supplement 12**. gHMBC spectrum of **3** in DMSO-$d_6$.
**Figure 1-figure supplement 13**. (+)-HRESITOFMS of compound **3** (top) with predicted elemental composition.
**Figure 1-figure supplement 14**.$^{1}$H NMR (500 MHz; top) and $^{13}$C NMR (125 MHz; bottom) spectra of **4** in DMSO-$d_6$.
**Figure 1-figure supplement 15**. gCOSY (top) and gHSQC (bottom) spectra of **4** in DMSO-$d_6$.
**Figure 1-figure supplement 16**. gHMBC spectrum of **4** in DMSO-$d_6$.
**Figure 1-figure supplement 17**. (+)-HRESITOFMS of compound **4** (top) with predicted elemental composition.
**Figure 1-figure supplement 18**. $^{1}$H NMR (500 MHz; top) and $^{13}$C NMR (125 MHz; bottom) spectra of **5** in DMSO-$d_6$.
**Figure 1-figure supplement 19**. gCOSY (top) and gHSQC (bottom) spectra of **5** in DMSO-$d_6$.
**Figure 1-figure supplement 20**. gHMBC spectrum of **5** in DMSO-$d_6$.
**Figure 1-figure supplement 21**. (+)-HRESITOFMS of compound **5** (top) with predicted elemental composition.
**Figure 1-figure supplement 22**. $^{1}$H NMR (500 MHz) spectrum of **1** (natural; top) and synthetic **1** (bottom) in DMSO-$d_6$.
**Figure 1-figure supplement 23**. $^{13}$C NMR (125 MHz) spectrum of **1**(natural; top) and synthetic **1** (bottom) in DMSO-$d_6$.
**Figure 1-figure supplement 24**. (+)-HRESITOFMS of synthesized compound **1** with predicted elemental composition.
**Figure 1-figure supplement 25**. Chromatographic co-elution profile of natural and synthetic **1**.
**Figure 1-figure supplement 26**. $^{1}$H NMR (500 MHz) spectrum of **3** (natural; top) and synthetic **3** (bottom) in DMSO-$d_6$.
**Figure 1-figure supplement 27**. $^{13}$C NMR (125 MHz) spectrum of **3** (top) and synthetic **3** (bottom) in DMSO-$d_6$.
**Figure 1-figure supplement 28**. (+)-HRESITOFMS of synthesized compound 3 with predicted elemental composition.
**Figure 1-figure supplement 29**. Chromatographic co-elution profile of natural and synthetic compound **3**.
**Figure 1-figure supplement 30**. UV spectrum of **1** in $CH_3OH$.
**Figure 1-figure supplement 31**. UV spectrum of **2** in $CH_3OH$.
**Figure 1-figure supplement 32**. UV spectrum of **3** in $CH_3OH$.
**Figure 1-figure supplement 33**. UV spectrum of **4** in $CH_3OH$.

**Figure 2-figure supplement 1**. Chemical structure and MS/MS fragmentation of compounds structurally-related to molleamines identified by metabolomics analysis of *D. molle* specimens.
**Figure 2-figure supplement 2**. Chemical structure and MS/MS fragmentation of compounds structurally-related to molleamines identified by metabolomics analysis of *D. molle* specimens.
**Figure 3-figure supplement 1**. Constellation pharmacology indicates that compounds **3-5** are selective against L2 DRG neurons, which are Aδ-low threshold mechanoreceptors (LTMRs).
**Figure 3-figure supplement 2.** Constellation pharmacology of molleamine C and transcriptomics analysis of neurons with responses to acetylcholine (ACh) blocked by molleamine C (20 µM) and TxID (1 µM).
**Figure 3-figure supplement 3**. Molleamine C does not affect α7-nAChRs.
**Figure 6-figure supplement 1**. Compound **3** inhibits responses from L2 neurons after depolarization with a P2Y1 agonist MRS 2365 (100 nM).
**Figure 6-figure supplement 2**. Compound **3** does not inhibit P2Y1 activity in human embryonic kidney (HEK-293) overexpressing GCaMP6s.
**Figure 7-figure supplement 1**. Stability of compound **3** in mouse plasma over 24 h.
**Figure 7-figure supplement 2**. Cytotoxicity evaluation of compound **3** in *in vitro* MTT assay with human embryonic kidney (HEK-293) cells.
**Figure 7-figure supplement 3.** Zebrafish photomotor-response after exposure to increasing concentrations of compound **3**.
**Figure 1-table supplement 1.** $^{1}H$ NMR (400 MHz) and $^{13}C$ NMR (125 MHz) data for compounds **1-5** in DMSO-$d_6$.
**Figure 3-table supplement 1**. Census of effects elicited by compounds **1-5** on ATP-induced depolarization in 16 DRG neuronal subtypes screened in calcium imaging experiments.
**Figure 3-table supplement 2**. Census of effects elicited by compounds **1-5** on $K^+$-induced depolarization in 16 DRG neuronal subtypes screened in calcium imaging experiments.
**Figure 6-table supplement 1**. Census of effects elicited by **3** on MRS2365-induced depolarization in 16 DRG neuronal subtypes screened in two calcium imaging experiments.

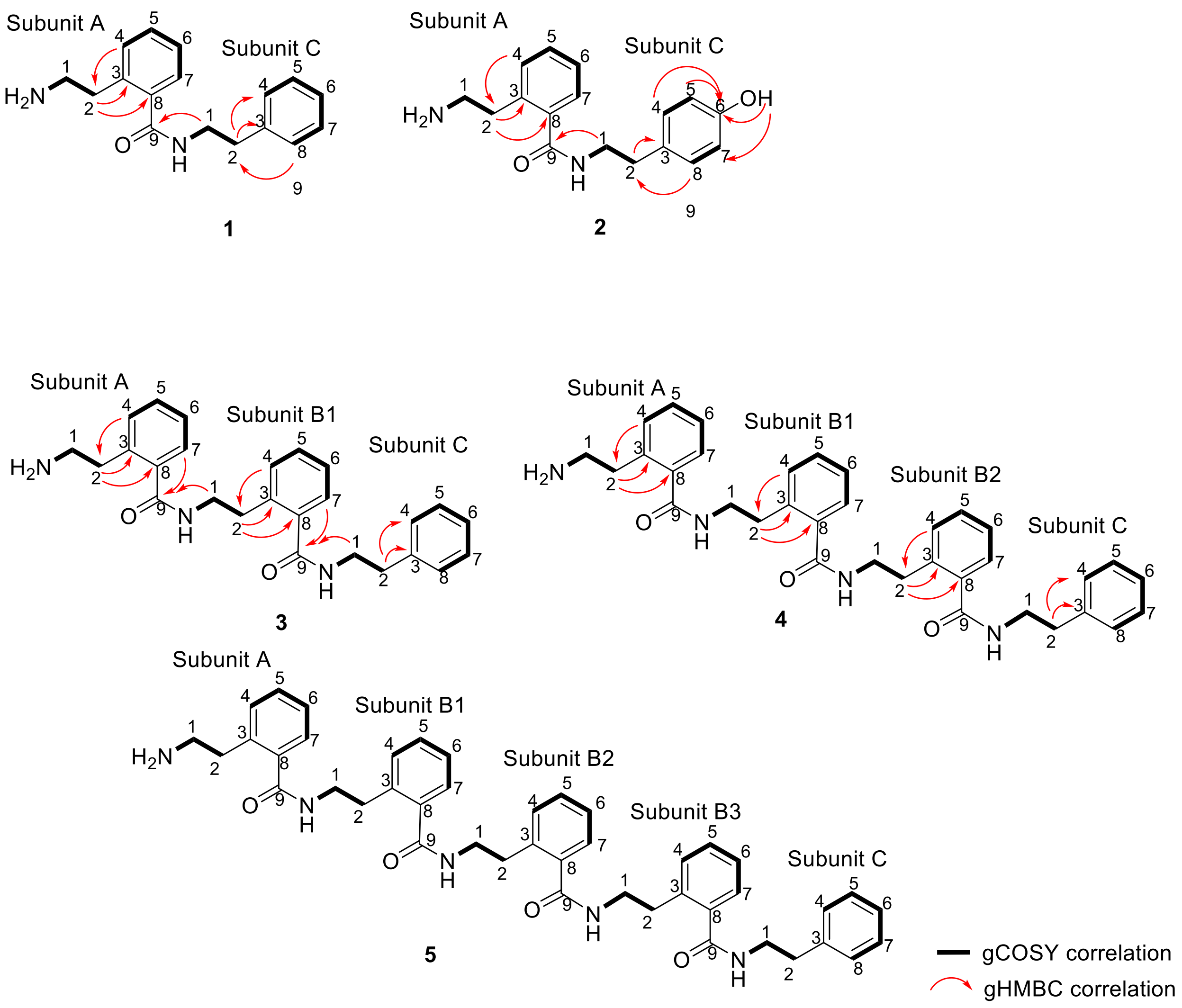


**Figure 1-figure supplement 1**. Structures of compounds **1-5** with COSY and HMBC correlations.

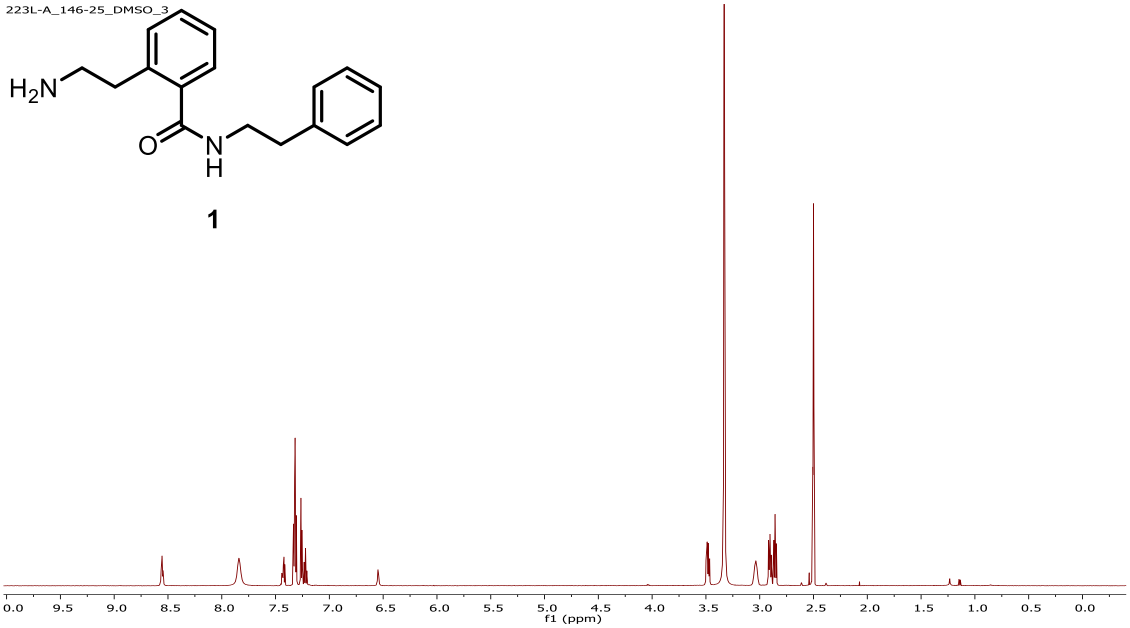


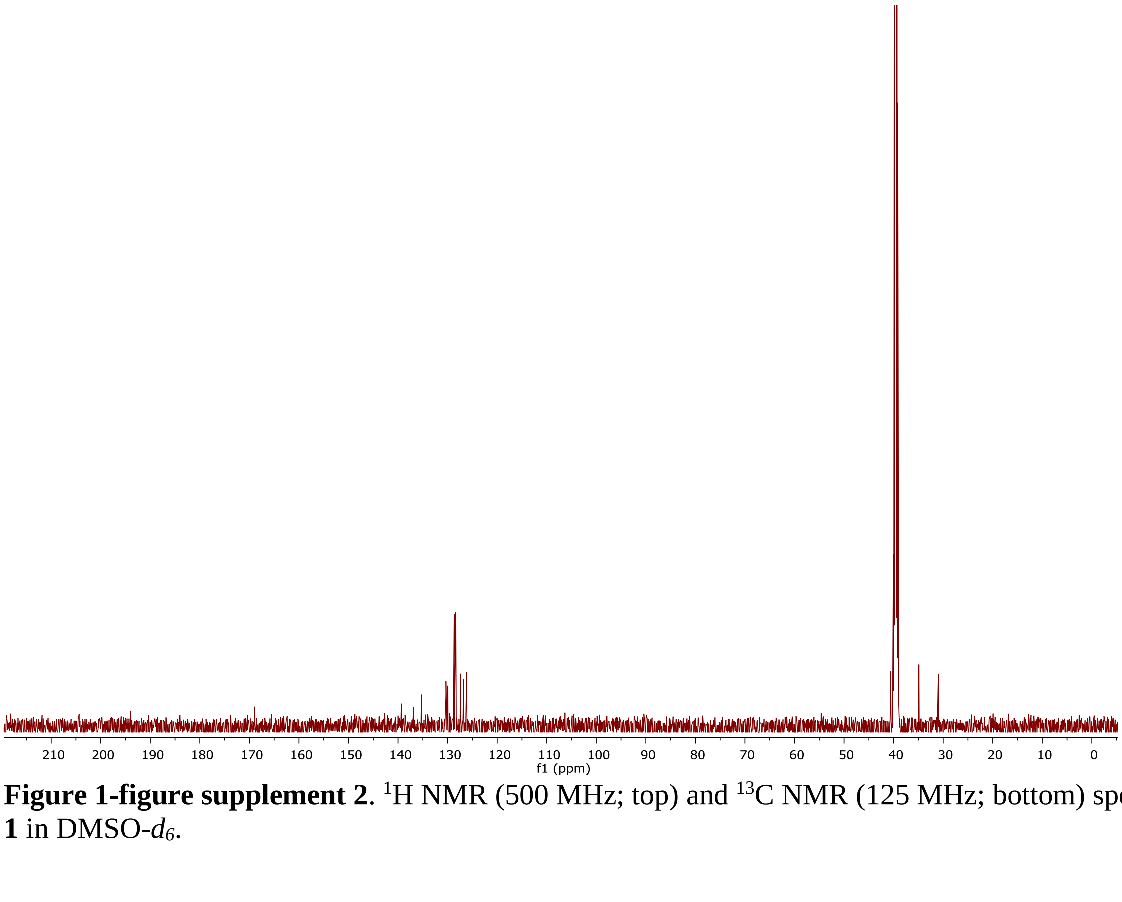


**Figure 1-figure supplement 2**. $^{1}$H NMR (500 MHz; top) and $^{13}$C NMR (125 MHz; bottom) spectra of **1** in DMSO-$d_6$.

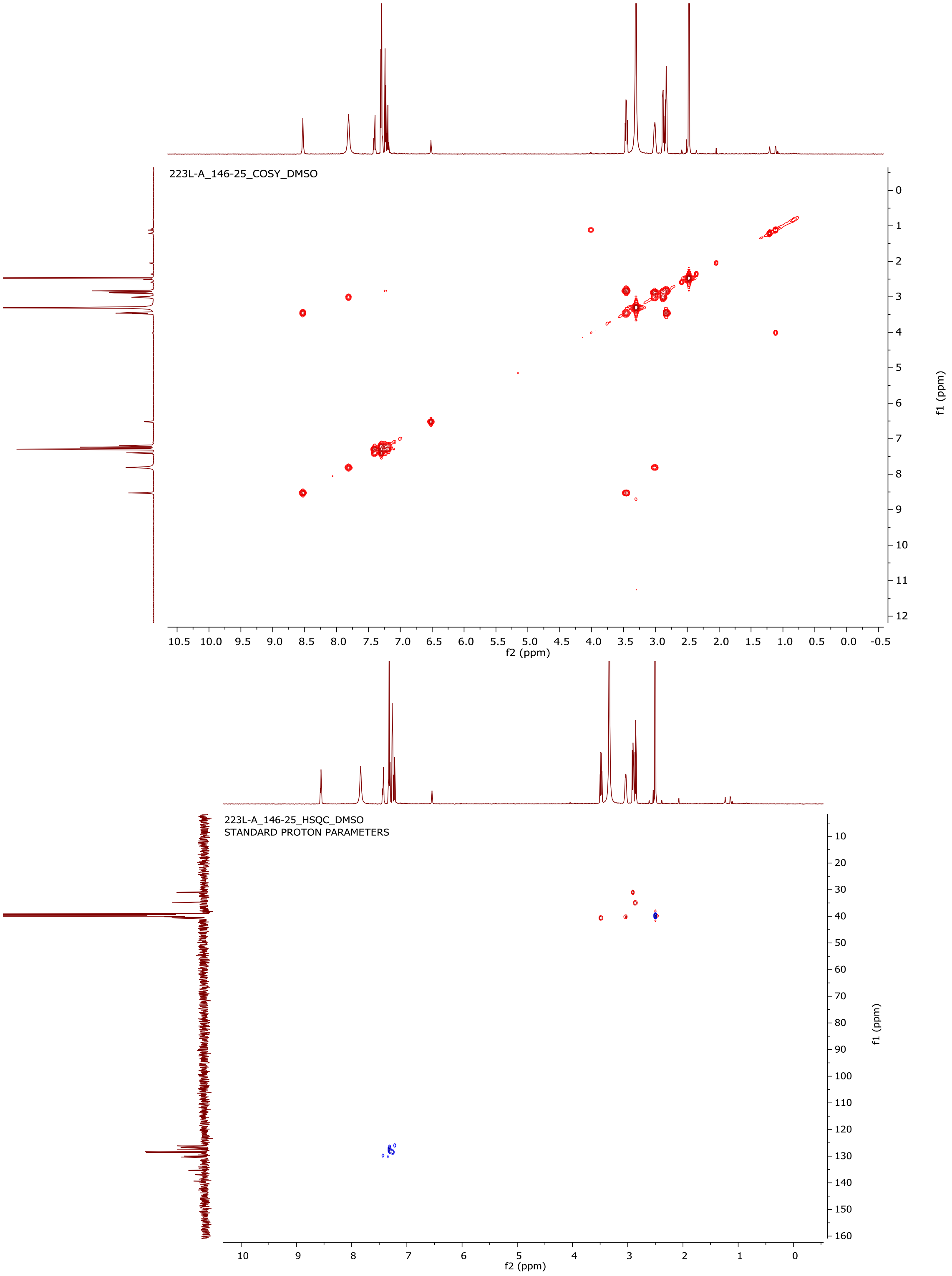


**Figure 1-figure supplement 3**. gCOSY (top) and gHSQC (bottom) spectra of **1** in DMSO-d6.

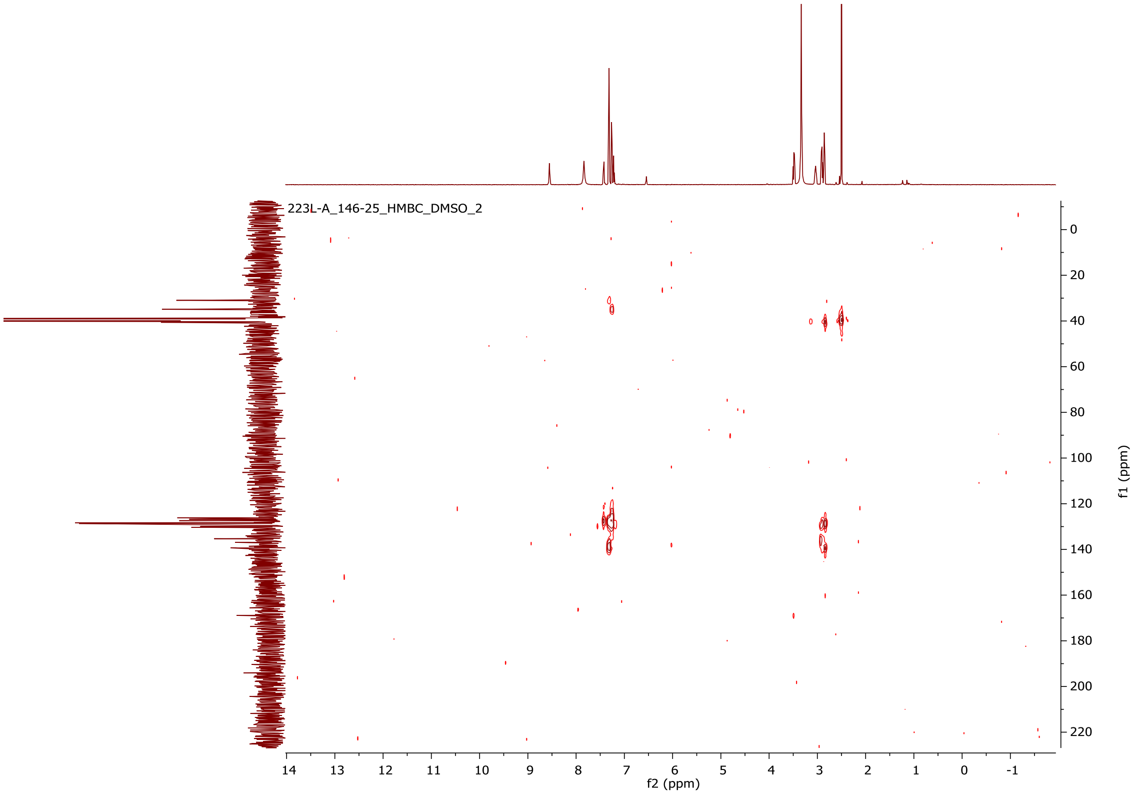


**Figure 1-figure supplement 4**. gHMBC spectrum of **1** in DMSO-d6

**Elemental Composition Report** **Page 1**

**Single Mass Analysis**
Tolerance = 5.0 PPM / DBE: min = -1.5, max = 50.0
Element prediction: Off
Number of isotope peaks used for i-FIT = 3

Monoisotopic Mass, Even Electron Ions
804 formula(e) evaluated with 2 results within limits (up to 50 best isotopic matches for each mass)
Elements Used:
C: 0-500 H: 0-1000 N: 0-200 O: 0-200 23Na: 0-1
1-146-25_7312019 CE = 6 Cone = 35
1-146-25_7312020 514 (4.573) AM2 (Ar,22000.0,0.00,0.00) 1: TOF MS ES+ 5.51e+006

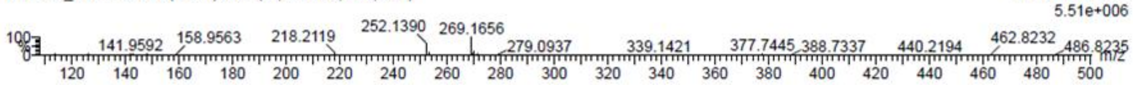


| | | | | | | | | |
|---|---|---|---|---|---|---|---|---|
| Minimum: | | | | -1.5 | | | | |
| Maximum: | | 5.0 | 5.0 | 50.0 | | | | |
| Mass | Calc. Mass | mDa | PPM | DBE | i-FIT | Norm | Conf(%) | Formula |
| 269.1656 | 269.1654 | 0.2 | 0.7 | 8.5 | 882.6 | 0.000 | 100.00 | C17 H21 N2 O |
| | 269.1659 | -0.3 | -1.1 | 1.5 | 902.7 | 20.081 | 0.00 | C2 H17 N14 O2 |

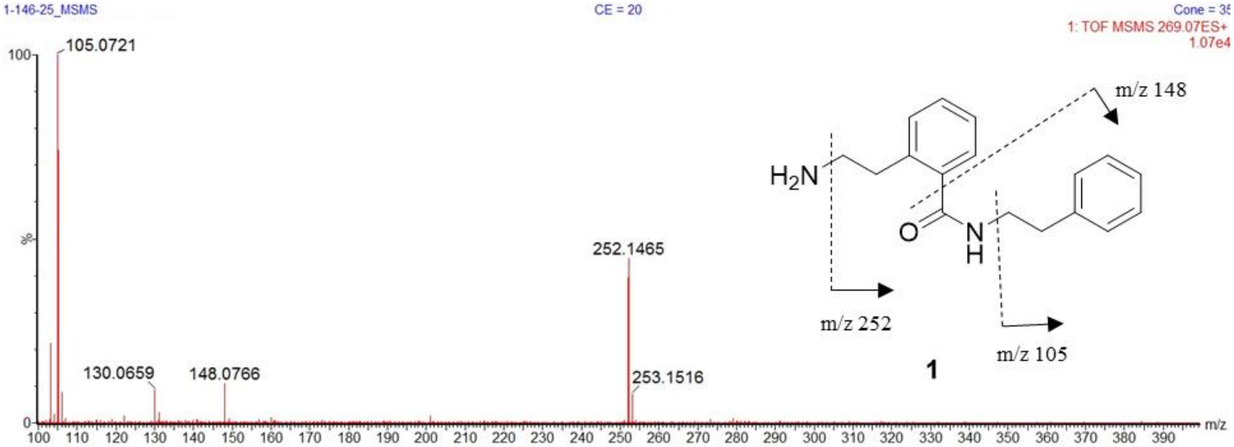


**Figure 1-figure supplement 5**. (+)-HRESITOFMS of compound **1** (top) with predicted elemental composition.

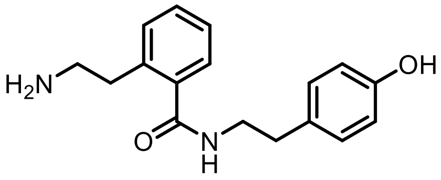


**2**

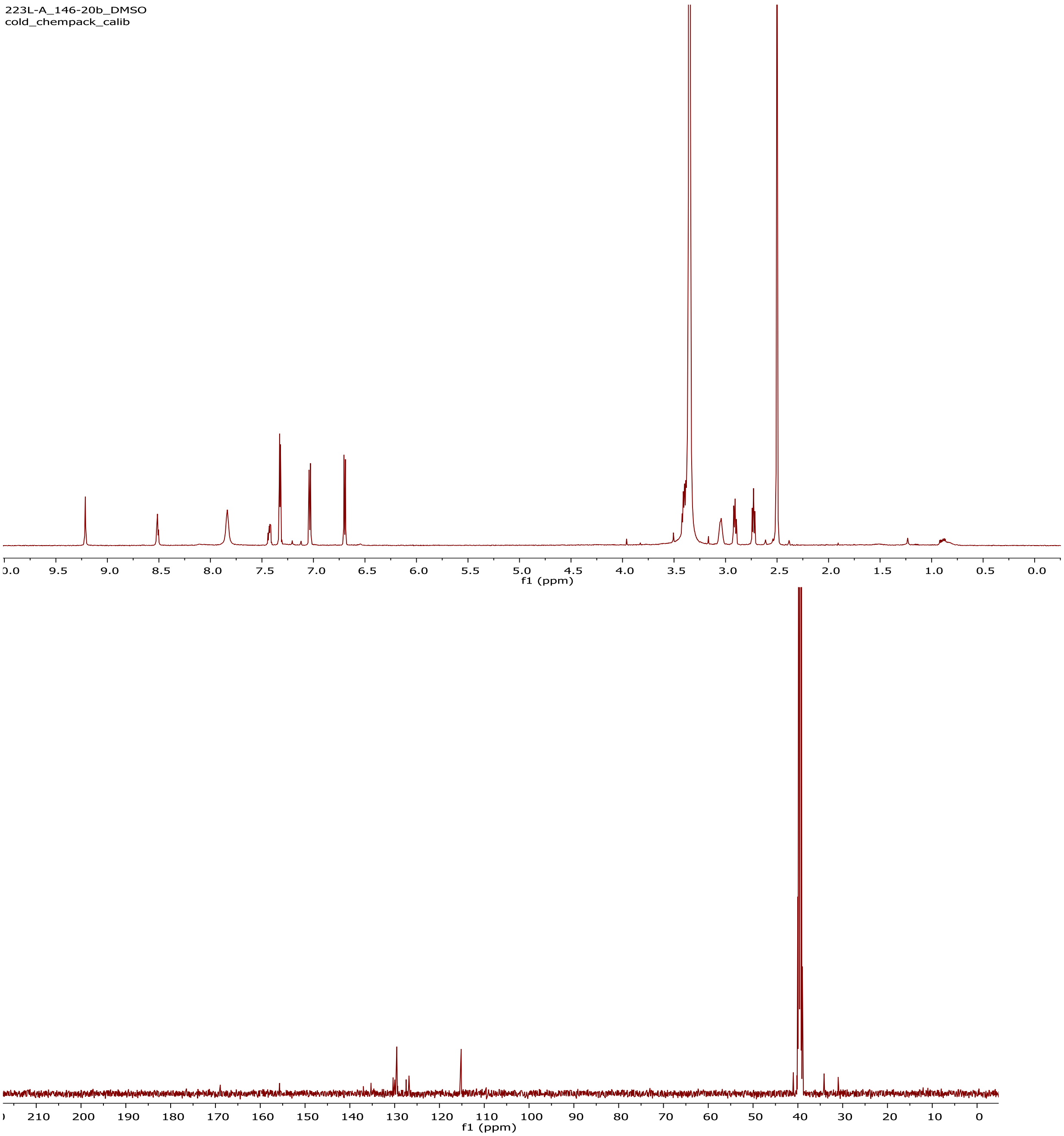


**Figure 1-figure supplement 6**.$^{1}$H NMR (500 MHz; top) and $^{13}$C NMR (125 MHz; bottom) spectra of **2** in DMSO-$d_6$.

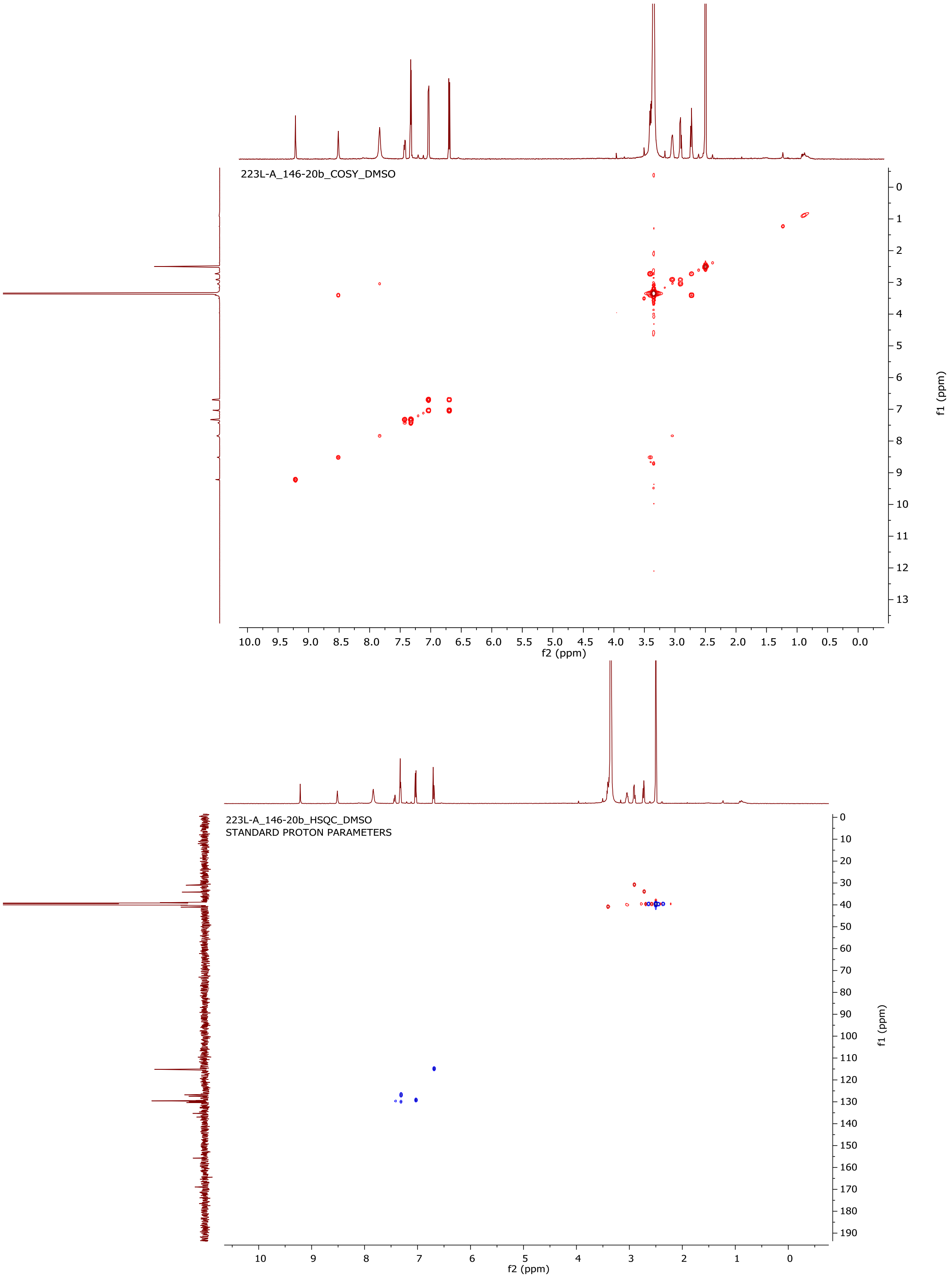


**Figure 1-figure supplement 7**. gCOSY (top) and gHSQC (bottom) spectra of **2** in DMSO-$d_6$.

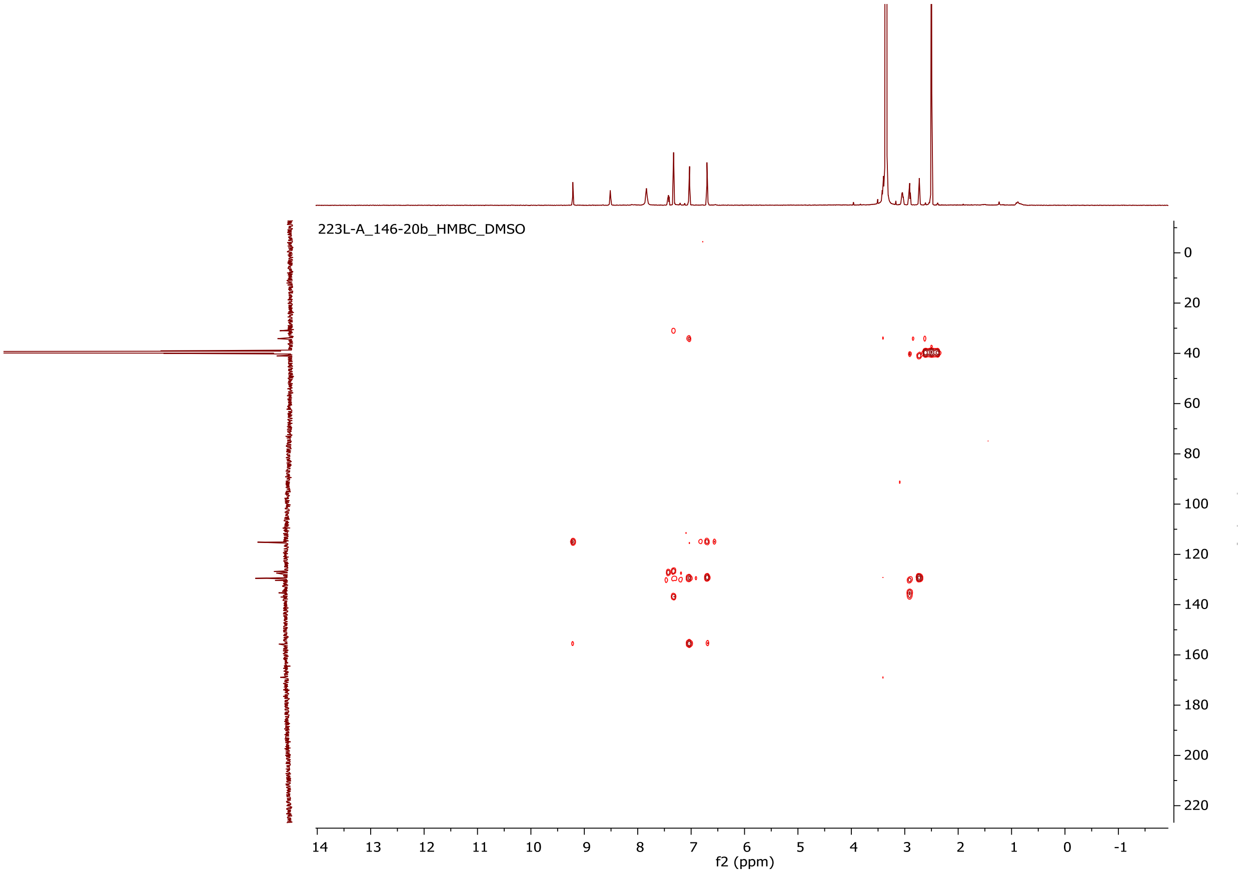


**Figure 1-figure supplement 8**. gHMBC spectrum of **2** taken in DMSO-$d_6$.

### Elemental Composition Report



#### Single Mass Analysis

Tolerance = 5.0 PPM / DBE: min = -1.5, max = 50.0
Element prediction: Off
Number of isotope peaks used for i-FIT = 3

Monoisotopic Mass, Even Electron Ions
934 formula(e) evaluated with 3 results within limits (up to 50 best isotopic matches for each mass)
Elements Used:
C: 0-500 H: 0-1000 N: 0-200 O: 0-200 23Na: 0-1

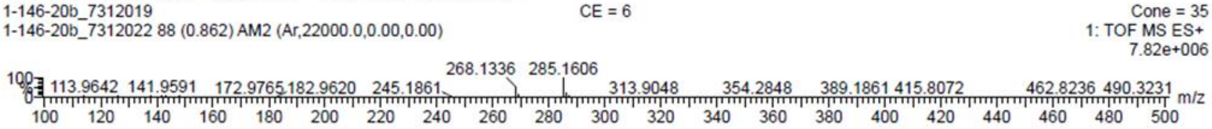


| | | | | | | | | |
|---|---|---|---|---|---|---|---|---|
| Minimum: | | | | -1.5 | | | | |
| Maximum: | | 5.0 | 5.0 | 50.0 | | | | |
| Mass | Calc. Mass | mDa | PPM | DBE | i-FIT | Norm | Conf(%) | Formula |
| 285.1606 | 285.1619 | -1.3 | -4.6 | 9.5 | 891.0 | 0.315 | 72.99 | C20 H22 23Na |
| | 285.1603 | 0.3 | 1.1 | 8.5 | 892.0 | 1.309 | 27.01 | C17 H21 N2 O2 |
| | 285.1608 | -0.2 | -0.7 | 1.5 | 907.7 | 16.961 | 0.00 | C2 H17 N14 O3 |

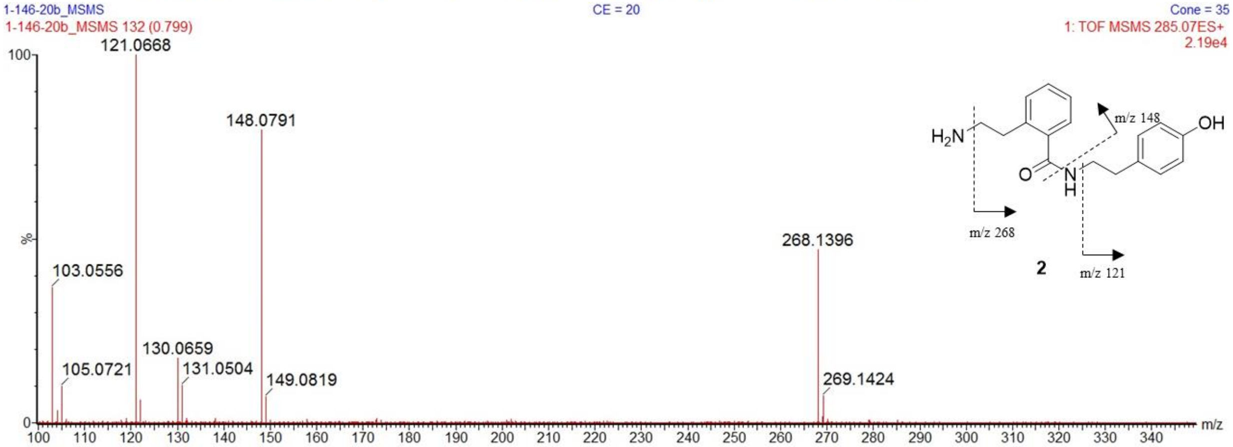


**Figure 1-figure supplement 9**. (+)-HRESITOFMS of compound **2** (top) with predicted elemental composition.

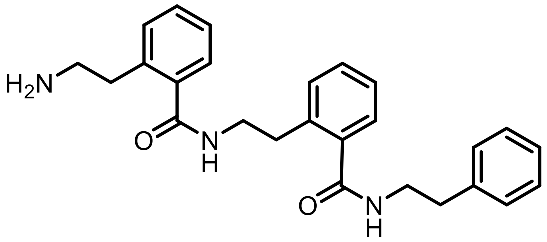


**3**

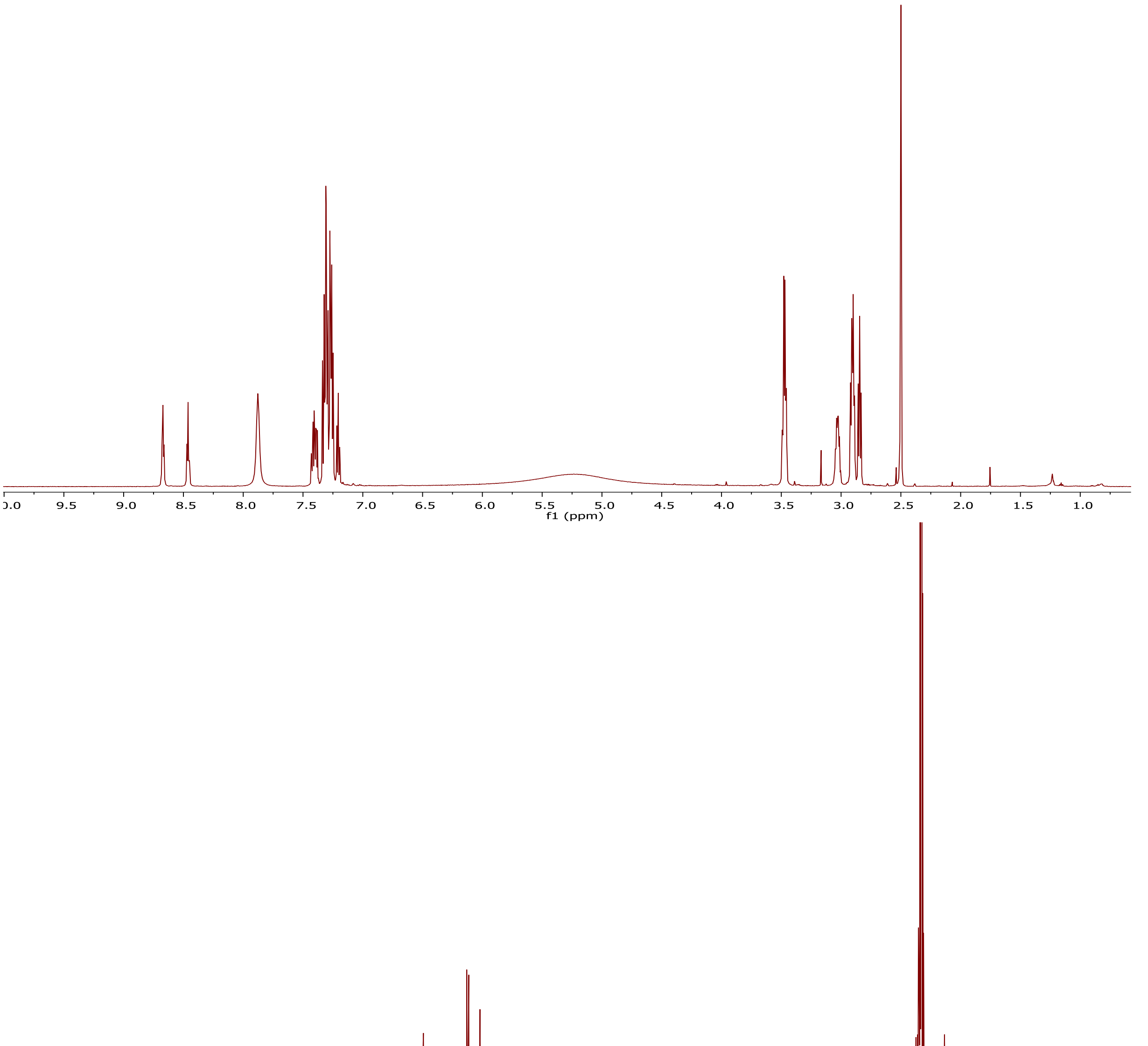


**Figure 1-figure supplement 10**.$^{1}$H NMR (500 MHz; top) and $^{13}$C NMR (125 MHz; bottom) spectra of **3** in DMSO-$d_6$.

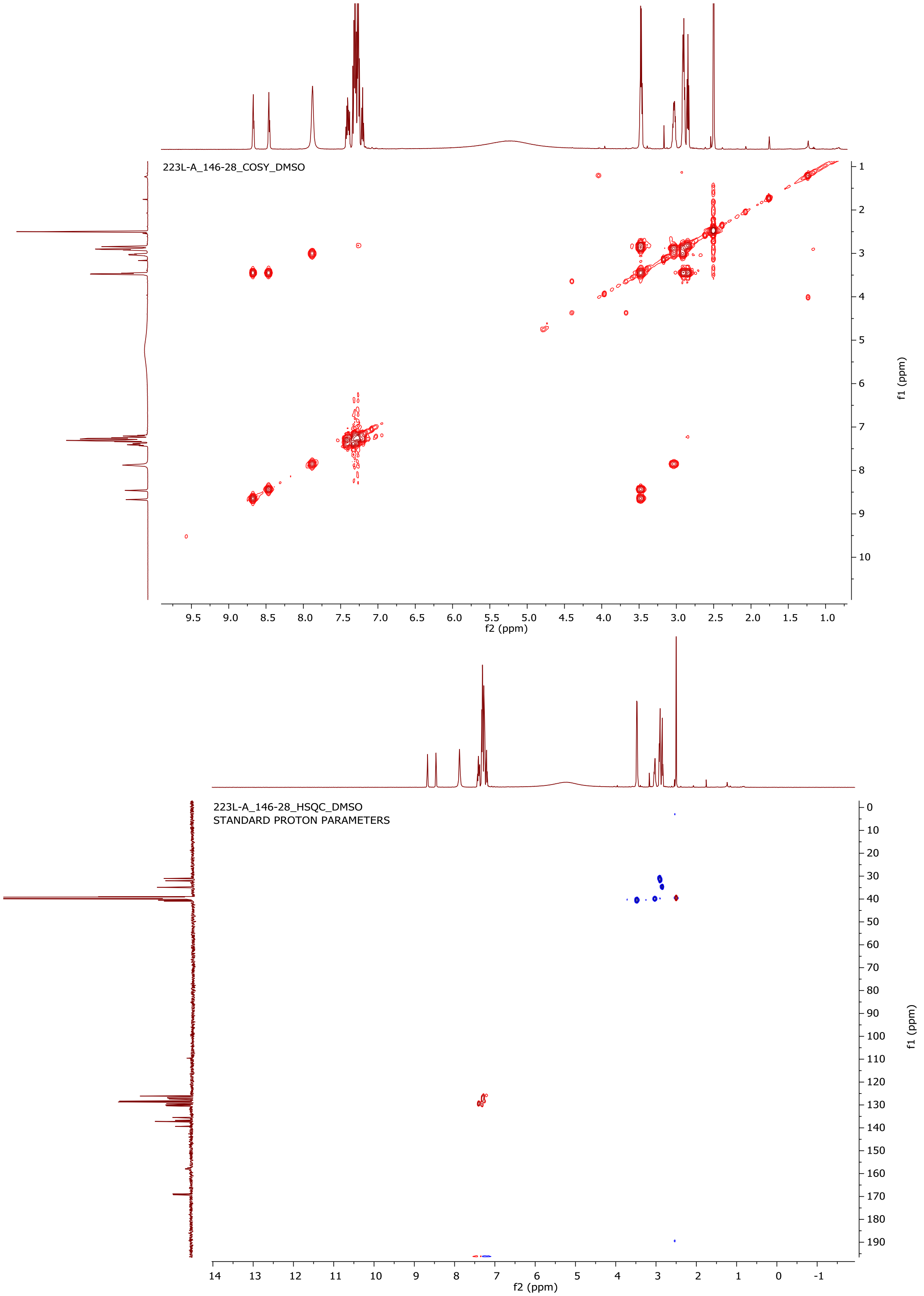


**Figure 1-figure supplement 11**. gCOSY (top) and gHSQC (bottom) spectra of **3** in DMSO-$d_6$.

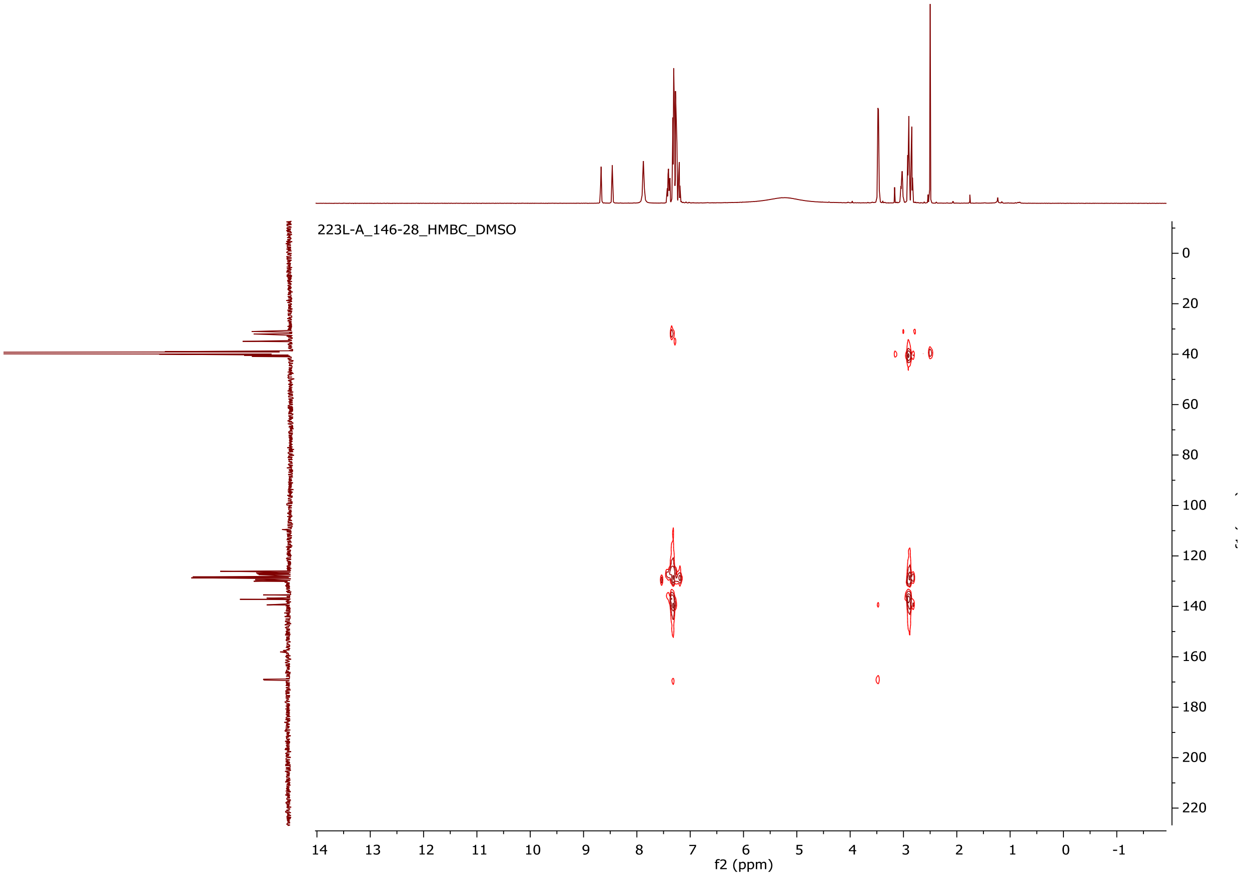

**Figure 1-figure supplement 12**. gHMBC spectrum of **3** in DMSO-$d_6$.

### Elemental Composition Report



#### Single Mass Analysis

Tolerance = 5.0 PPM / DBE: min = -1.5, max = 50.0
Element prediction: Off
Number of isotope peaks used for i-FIT = 3

Monoisotopic Mass, Even Electron Ions
2542 formula(e) evaluated with 7 results within limits (up to 50 best isotopic matches for each mass)
Elements Used:
C: 0-500 H: 0-1000 N: 0-200 O: 0-200 23Na: 0-1

1-146-28_7312019 CE = 6 Cone = 35
1-146-28_7312019 505 (4.488) AM2 (Ar,22000.0,0.00,0.00) 1: TOF MS ES+ 1.66e+006

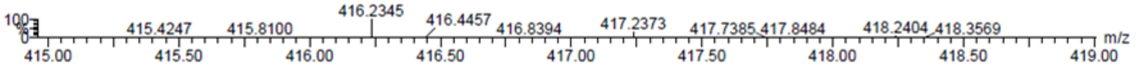


| | | | | | | | | |
|---|---|---|---|---|---|---|---|---|
| Minimum: | | | | -1.5 | | | | |
| Maximum: | | 5.0 | 5.0 | 50.0 | | | | |
| Mass | Calc. Mass | mDa | PPM | DBE | i-FIT | Norm | Conf(%) | Formula |
| 416.2345 | 416.2338 | 0.7 | 1.7 | 13.5 | 519.2 | 0.002 | 99.78 | C26 H30 N3 O2 |
| | 416.2354 | -0.9 | -2.2 | 14.5 | 525.3 | 6.116 | 0.22 | C29 H31 N 23Na |
| | 416.2357 | -1.2 | -2.9 | 0.5 | 530.3 | 11.114 | 0.00 | C14 H34 N5 O9 |
| | 416.2346 | -0.1 | -0.2 | 2.5 | 531.2 | 11.980 | 0.00 | C13 H31 N9 O5 23Na |
| | 416.2359 | -1.4 | -3.4 | 7.5 | 531.7 | 12.544 | 0.00 | C14 H27 N13 O 23Na |
| | 416.2343 | 0.2 | 0.5 | 6.5 | 533.7 | 14.513 | 0.00 | C11 H26 N15 O3 |
| | 416.2330 | 1.5 | 3.6 | 1.5 | 534.0 | 14.781 | 0.00 | C10 H30 N11 O7 |

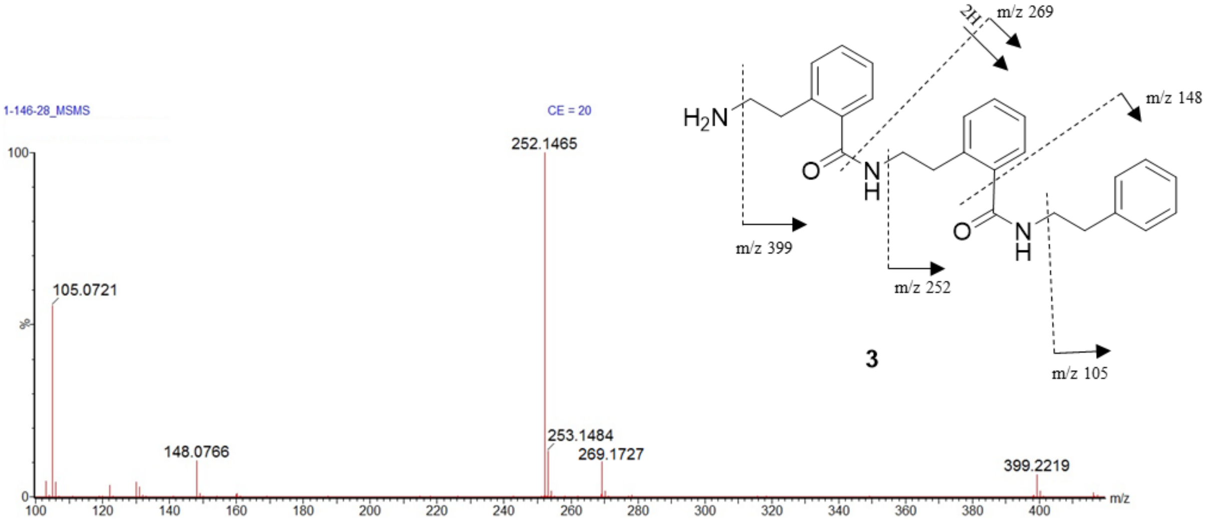


**Figure 1-figure supplement 13**. (+)-HRESITOFMS of compound **3** (top) with predicted elemental composition. (+)-CID MS/MS (20 eV) of *m/z* 416 showing a distinct fragmentation pattern (bottom).

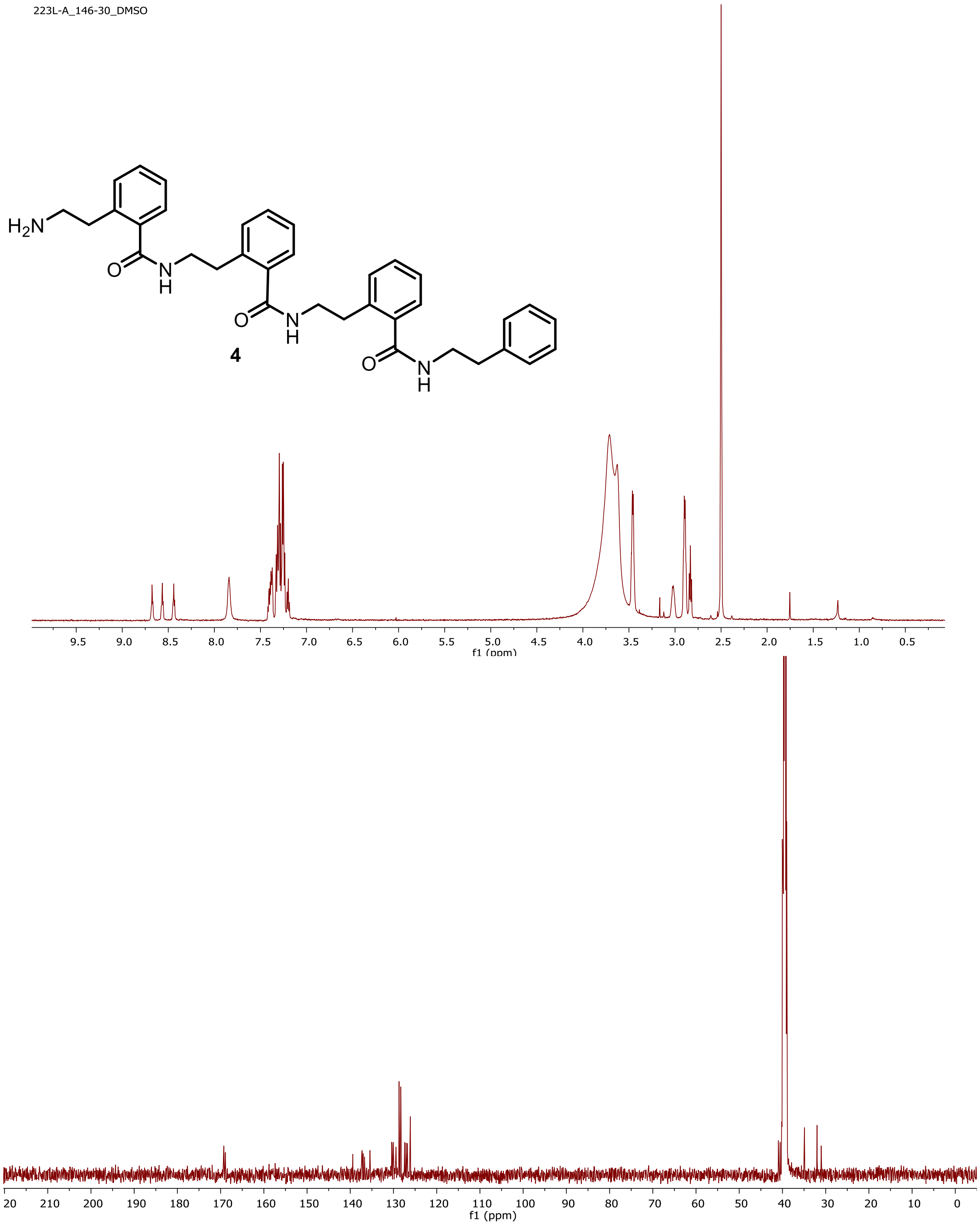


**Figure 1-figure supplement 14**.$^{1}$H NMR (500 MHz; top) and $^{13}$C NMR (125 MHz; bottom) spectra of **4** in DMSO-$d_6$.

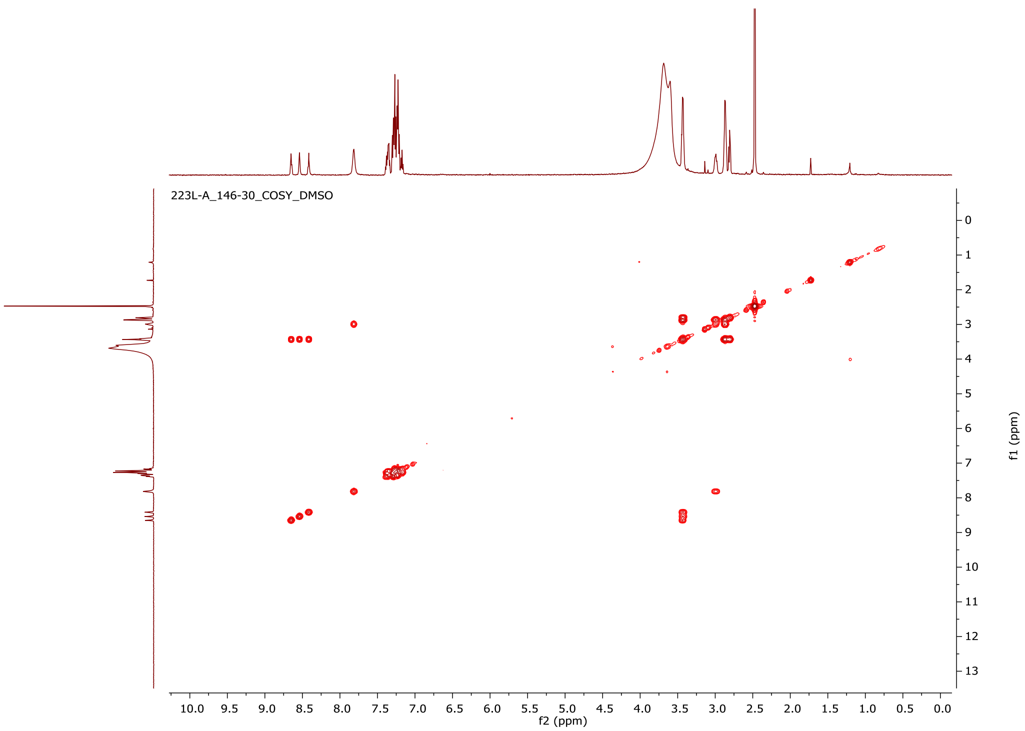


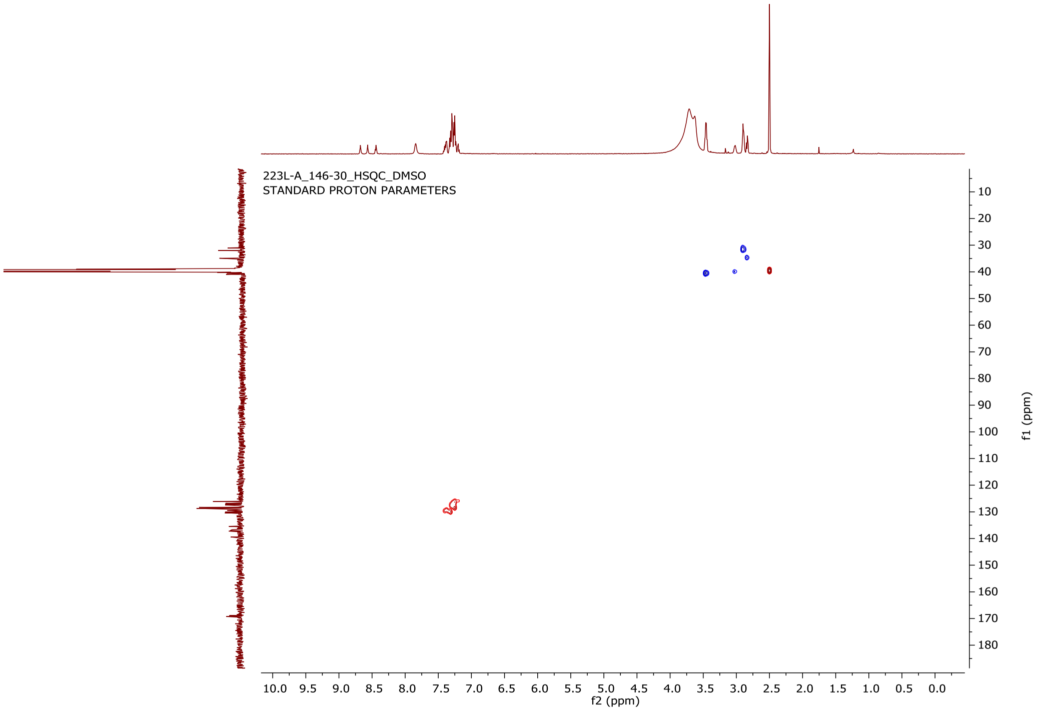


**Figure 1-figure supplement 15**. gCOSY (top) and gHSQC (bottom) spectra of **4** in DMSO-$d_6$.

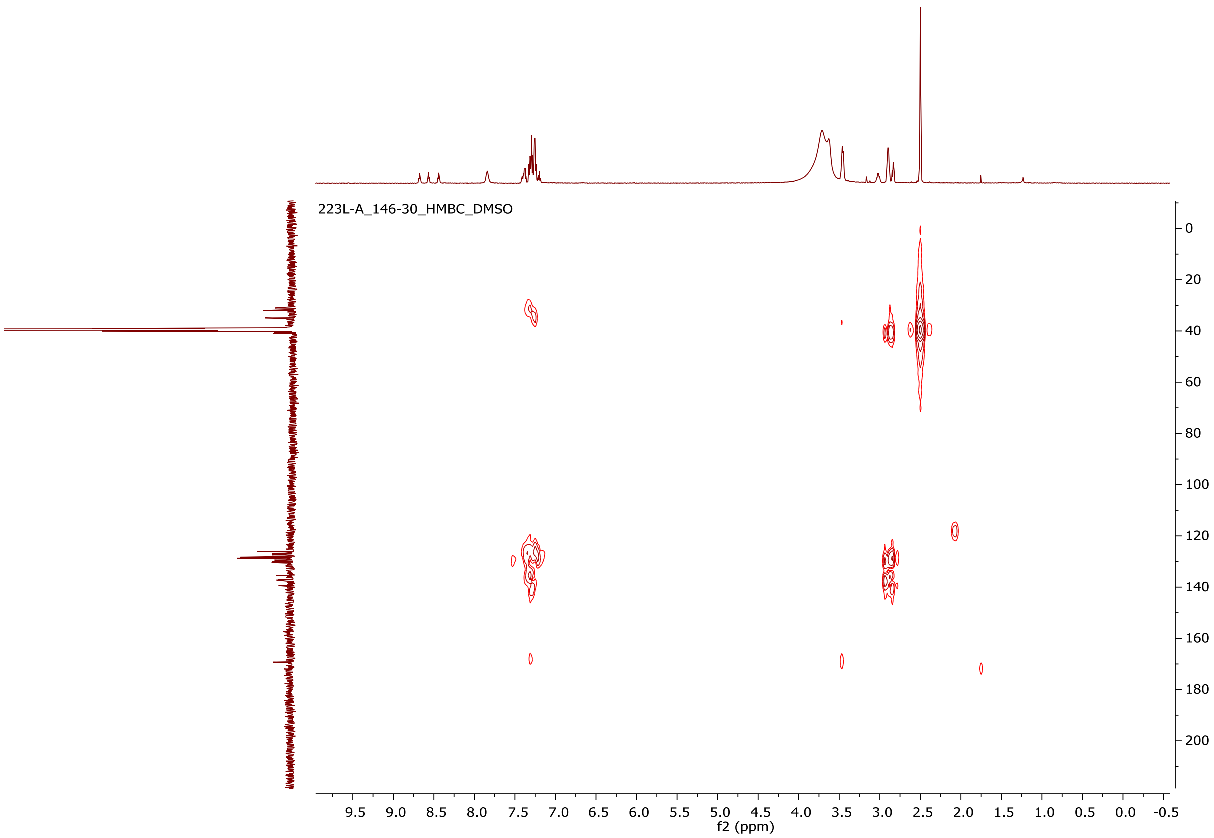


**Figure 1-figure supplement 16**. gHMBC spectrum of **4** in DMSO-$d_6$.

**Elemental Composition Report** **Page 1**

**Single Mass Analysis**
Tolerance = 5.0 PPM / DBE: min = -1.5, max = 50.0
Element prediction: Off
Number of isotope peaks used for i-FIT = 3

Monoisotopic Mass, Even Electron Ions
9721 formula(e) evaluated with 32 results within limits (up to 50 best isotopic matches for each mass)
Elements Used:
C: 0-500 H: 0-1000 N: 0-200 O: 0-200 23Na: 0-1 127I: 0-2

1-146-30_08072019 CE = 6 Cone = 35
1-146-30_08072019 629 (5.574) AM2 (Ar,22000.0,0.00,0.00) 1: TOF MS ES+ 2.56e+006

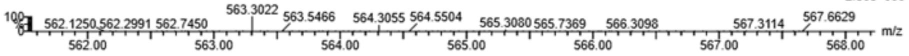


| | | mDa | PPM | DBE | | | | |
|---|---|---|---|---|---|---|---|---|
| Minimum: | | | | -1.5 | | | | |
| Maximum: | | 5.0 | 5.0 | 50.0 | | | | |
| **Mass** | **Calc. Mass** | **mDa** | **PPM** | **DBE** | **i-FIT** | **Norm** | **Conf(%)** | **Formula** |
| 563.3022 | 563.3009 | 1.3 | 2.3 | 13.5 | 555.8 | 0.070 | 93.27 | C34 H43 O7 |
| | 563.3022 | 0.0 | 0.0 | 18.5 | 558.5 | 2.788 | 6.15 | C35 H39 N4 O3 |
| | 563.2998 | 2.4 | 4.3 | 15.5 | 561.0 | 5.260 | 0.52 | C33 H40 N4 O3 23Na |
| | 563.2995 | 2.7 | 4.8 | 19.5 | 563.9 | 8.129 | 0.03 | C31 H35 N10 O |
| | 563.3038 | -1.6 | -2.8 | 19.5 | 564.3 | 8.548 | 0.02 | C38 H40 N2 O 23Na |
| | 563.3027 | -0.5 | -0.9 | 0.5 | 567.1 | 11.394 | 0.00 | C22 H47 N2 O14 |
| | 563.3030 | -0.8 | -1.4 | 7.5 | 567.4 | 11.661 | 0.00 | C22 H40 N10 O6 23Na |
| | 563.3041 | -1.9 | -3.4 | 5.5 | 567.9 | 12.101 | 0.00 | C23 H43 N6 O10 |
| | 563.3017 | 0.5 | 0.9 | 2.5 | 567.9 | 12.137 | 0.00 | C21 H44 N6 O10 23Na |
| | 563.3023 | -0.1 | -0.2 | 0.5 | 568.1 | 12.328 | 0.00 | C21 H49 N8 23Na 127I |
| | 563.3033 | -1.1 | -2.0 | -1.5 | 568.1 | 12.369 | 0.00 | C22 H52 N4 O4 127I |
| | 563.3043 | -2.1 | -3.7 | 12.5 | 568.3 | 12.567 | 0.00 | C23 H36 N14 O2 23Na |
| | 563.3043 | -2.1 | -3.7 | 1.5 | 568.4 | 12.645 | 0.00 | C25 H48 O12 23Na |
| | 563.3027 | -0.5 | -0.9 | 11.5 | 569.4 | 13.682 | 0.00 | C20 H35 N16 O4 |
| | 563.3047 | -2.5 | -4.4 | 3.5 | 569.6 | 13.818 | 0.00 | C23 H48 N8 127I |
| | 563.3014 | 0.8 | 1.4 | 6.5 | 569.9 | 14.179 | 0.00 | C19 H39 N12 O8 |
| | 563.3041 | -1.9 | -3.4 | 16.5 | 569.9 | 14.187 | 0.00 | C21 H31 N20 |
| | 563.3049 | -2.7 | -4.8 | -0.5 | 570.3 | 14.522 | 0.00 | C25 H53 N2 O2 23Na 127I |
| | 563.3006 | 1.6 | 2.8 | -0.5 | 571.3 | 15.521 | 0.00 | C18 H48 N10 O2 127I |
| | 563.3000 | 2.2 | 3.9 | 1.5 | 571.3 | 15.600 | 0.00 | C18 H43 N8 O12 |
| | 563.3017 | 0.5 | 0.9 | 13.5 | 571.5 | 15.738 | 0.00 | C19 H32 N20 23Na |
| | 563.3003 | 1.9 | 3.4 | 8.5 | 572.1 | 16.313 | 0.00 | C18 H36 N16 O4 23Na |
| | 563.3000 | 2.2 | 3.9 | 12.5 | 573.6 | 17.867 | 0.00 | C16 H31 N22 O2 |
| | 563.3046 | -2.4 | -4.3 | -1.5 | 574.3 | 18.507 | 0.00 | C8 H39 N18 O11 |
| | 563.3035 | -1.3 | -2.3 | 0.5 | 575.4 | 19.607 | 0.00 | C7 H36 N22 O7 23Na |
| | 563.3048 | -2.6 | -4.6 | 5.5 | 575.9 | 20.133 | 0.00 | C8 H32 N26 O3 23Na |
| | 563.3019 | 0.3 | 0.5 | -0.5 | 576.6 | 20.866 | 0.00 | C4 H35 N24 O9 |
| | 563.3032 | -1.0 | -1.8 | 4.5 | 576.7 | 20.986 | 0.00 | C5 H31 N28 O5 |
| | 563.3046 | -2.4 | -4.3 | 9.5 | 577.5 | 21.750 | 0.00 | C6 H27 N32 O |
| | 563.3008 | 1.4 | 2.5 | 1.5 | 577.9 | 22.143 | 0.00 | C3 H32 N28 O5 23Na |
| | 563.3022 | 0.0 | 0.0 | 6.5 | 578.5 | 22.783 | 0.00 | C4 H28 N32 O 23Na |
| | 563.3005 | 1.7 | 3.0 | 5.5 | 579.2 | 23.417 | 0.00 | C H27 N34 O3 |

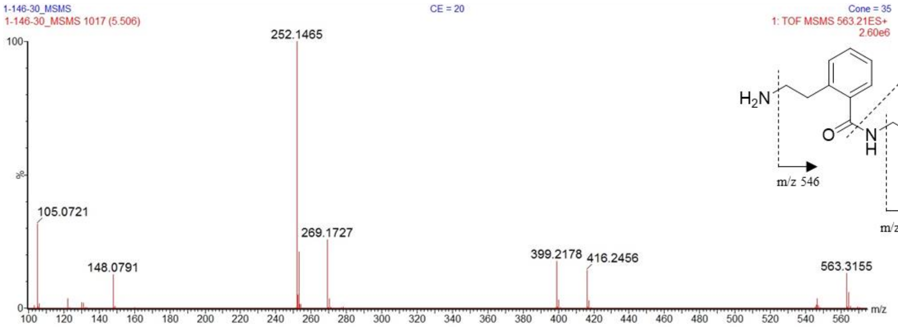


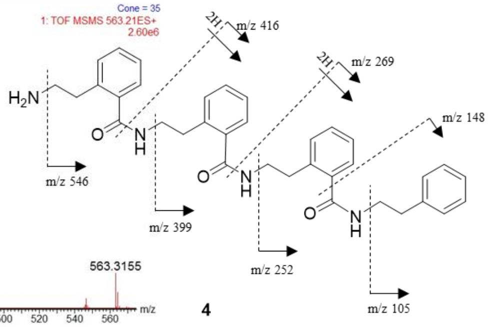


**4**

**Figure 1-figure supplement 17**. (+)-HRESITOFMS of compound **4** (top) with predicted elemental composition. (+)-CID MS/MS (20 eV) of *m/z* 563 showing a distinct fragmentation pattern (bottom).

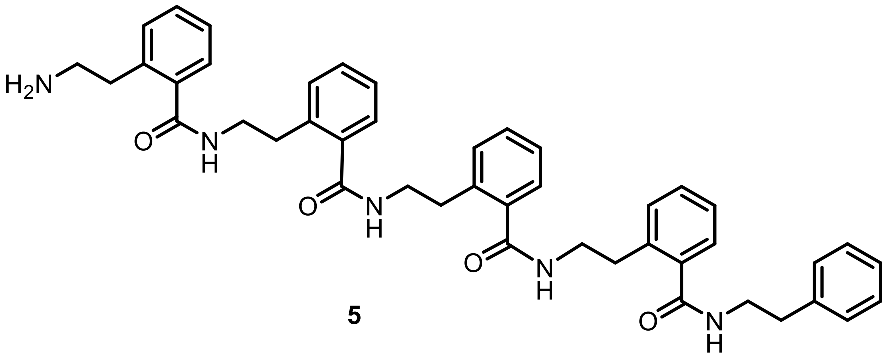


**5**

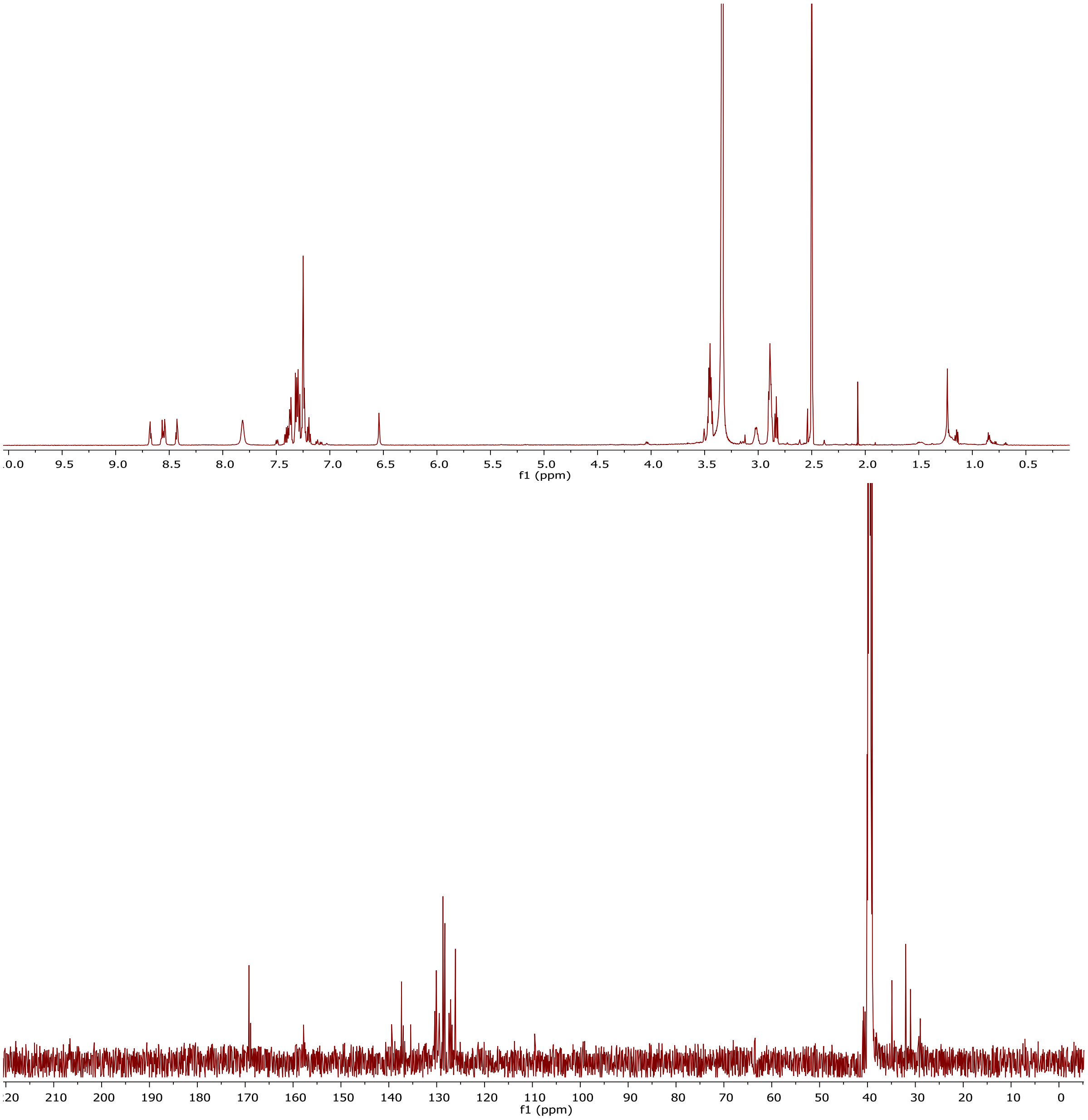


**Figure 1-figure supplement 18**. $^{1}H$ NMR (500 MHz; top) and $^{13}C$ NMR (125 MHz; bottom) spectra of **5** in DMSO-$d_6$.

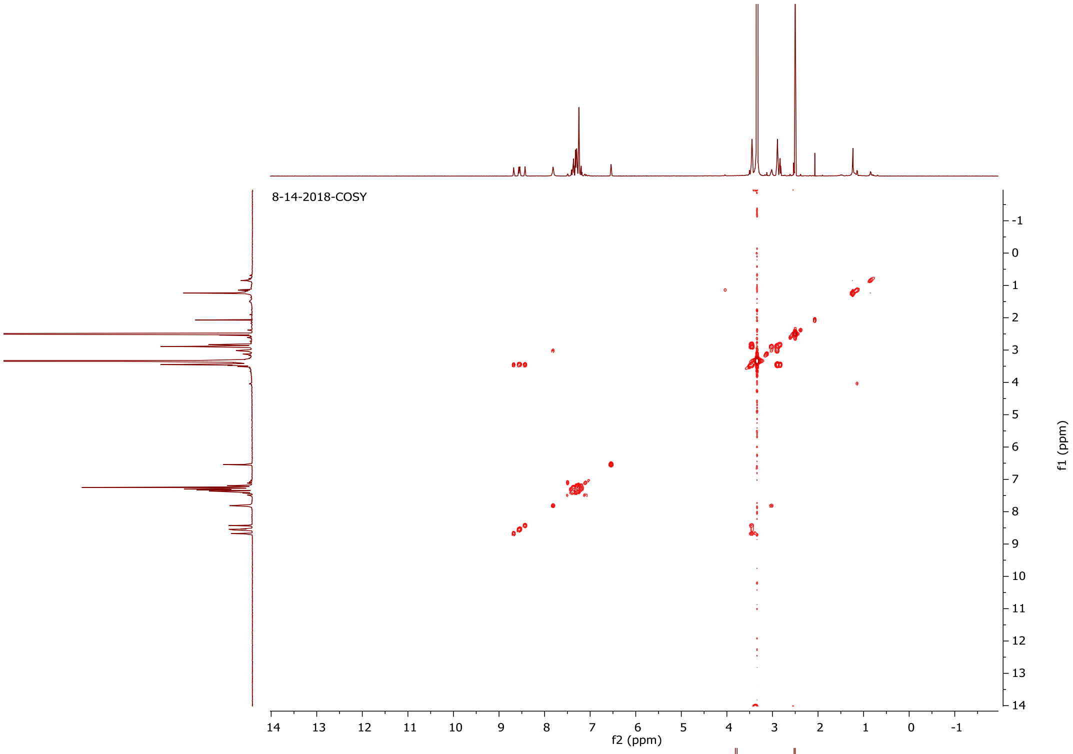


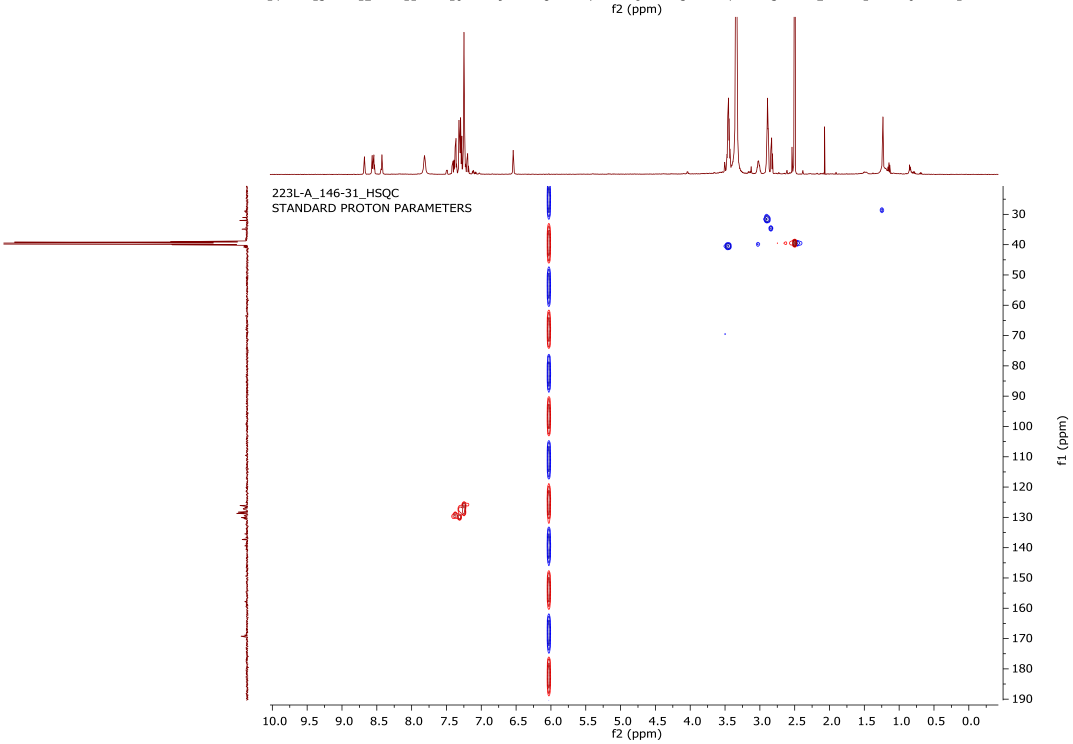


**Figure 1-figure supplement 19**. gCOSY (top) and gHSQC (bottom) spectra of **5** in DMSO-$d_6$.

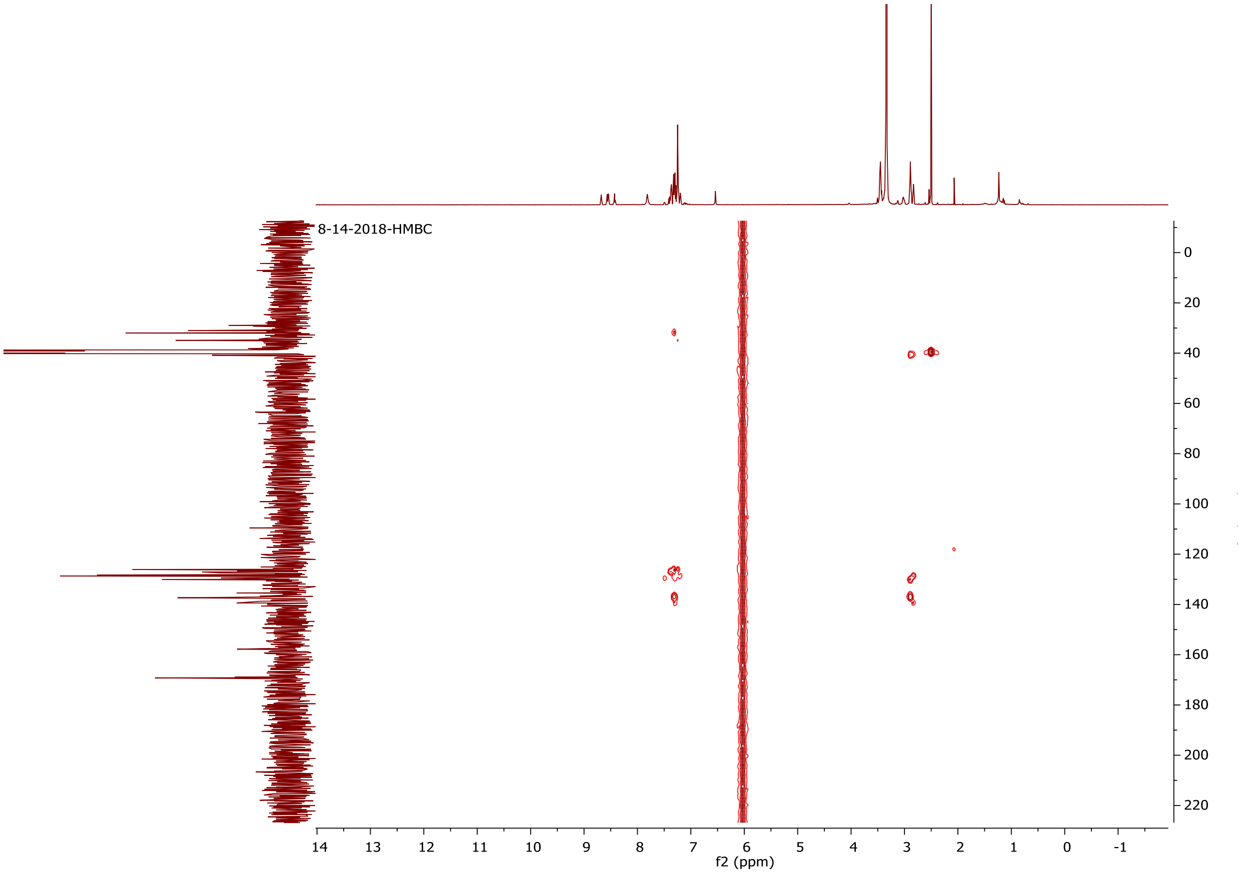


**Figure 1-figure supplement 20.** gHMBC spectrum of **5** in DMSO-$d_6$.

### Elemental Composition Report

Page 1

#### Single Mass Analysis

Tolerance = 5.0 PPM / DBE: min = -1.5, max = 50.0
Element prediction: Off
Number of isotope peaks used for i-FIT = 3

Monoisotopic Mass, Even Electron Ions
11084 formula(e) evaluated with 47 results within limits (up to 50 best isotopic matches for each mass)
Elements Used:
C: 0-500 H: 0-1000 N: 0-200 O: 0-200 23Na: 0-1
1-146-31_7312019
1-146-31_7312021 600 (5.318) AM2 (Ar,22000.0,0.00,0.00)

CE = 6

Cone = 35
1: TOF MS ES+
1.93e+006

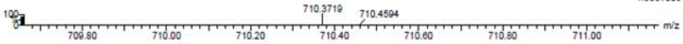


| Mass | Calc. Mass | mDa | PPM | DBE | i-FIT | Norm | Conf(%) | Formula |
|---|---|---|---|---|---|---|---|---|
| Minimum: | | | | -1.5 | | | | |
| Maximum: | | 5.0 | 5.0 | 50.0 | | | | |
| 710.3719 | 710.3724 | -0.5 | -0.7 | -1.5 | 58.6 | 2.739 | 6.46 | C H41 N35 O9 23Na |
| | 710.3716 | 0.3 | 0.4 | -1.5 | 58.9 | 3.082 | 4.59 | C16 H52 N15 O16 |
| | 710.3719 | 0.0 | 0.0 | 5.5 | 59.0 | 3.143 | 4.32 | C16 H45 N23 O8 23Na |
| | 710.3716 | 0.3 | 0.4 | 9.5 | 59.0 | 3.155 | 4.26 | C14 H40 N29 O6 |
| | 710.3735 | -1.6 | -2.3 | 7.5 | 59.2 | 3.346 | 3.52 | H32 N45 O3 |
| | 710.3730 | -1.1 | -1.5 | 3.5 | 59.3 | 3.423 | 3.26 | C17 H48 N19 O12 |
| | 710.3706 | 1.3 | 1.8 | 0.5 | 59.3 | 3.454 | 3.16 | C15 H49 N19 O12 23Na |
| | 710.3714 | 0.5 | 0.7 | 12.5 | 59.3 | 3.466 | 3.12 | C31 H49 N11 O7 23Na |
| | 710.3711 | 0.8 | 1.1 | 5.5 | 59.3 | 3.491 | 3.05 | C31 H56 N3 O15 |
| | 710.3730 | -1.1 | -1.5 | 14.5 | 59.3 | 3.491 | 3.05 | C15 H36 N33 O2 |
| | 710.3725 | -0.6 | -0.8 | 10.5 | 59.4 | 3.495 | 3.03 | C32 H52 N7 O11 |
| | 710.3711 | 0.8 | 1.1 | 16.5 | 59.4 | 3.537 | 2.91 | C29 H44 N17 O5 |
| | 710.3725 | -0.6 | -0.8 | 21.5 | 59.4 | 3.539 | 2.90 | C30 H40 N21 O |
| | 710.3738 | -1.9 | -2.7 | 3.5 | 59.4 | 3.552 | 2.87 | C2 H37 N39 O5 23Na |
| | 710.3706 | 1.3 | 1.8 | 11.5 | 59.4 | 3.563 | 2.83 | C13 H37 N33 O2 23Na |
| | 710.3728 | -0.9 | -1.3 | 6.5 | 59.4 | 3.582 | 2.78 | C34 H57 N O13 23Na |
| | 710.3733 | -1.4 | -2.0 | -0.5 | 59.4 | 3.592 | 2.75 | C19 H53 N13 O14 23Na |
| | 710.3720 | -0.1 | -0.1 | 28.5 | 59.5 | 3.607 | 2.71 | C45 H44 N9 |
| | 710.3733 | -1.4 | -2.0 | 10.5 | 59.5 | 3.623 | 2.67 | C17 H41 N27 O4 23Na |
| | 710.3728 | -0.9 | -1.3 | 17.5 | 59.5 | 3.625 | 2.66 | C32 H45 N15 O3 23Na |
| | 710.3703 | 1.6 | 2.3 | 4.5 | 59.5 | 3.631 | 2.65 | C13 H44 N25 O10 |
| | 710.3722 | -0.3 | -0.4 | 24.5 | 59.5 | 3.640 | 2.62 | C47 H49 N3 O2 23Na |
| | 710.3703 | 1.6 | 2.3 | 15.5 | 59.6 | 3.708 | 2.45 | C11 H32 N39 |
| | 710.3706 | 1.3 | 1.8 | 23.5 | 59.8 | 3.960 | 1.91 | C44 H48 N5 O4 |
| | 710.3701 | 1.8 | 2.5 | 7.5 | 59.9 | 4.086 | 1.68 | C30 H53 N7 O11 23Na |
| | 710.3701 | 1.8 | 2.5 | 18.5 | 60.0 | 4.134 | 1.60 | C28 H41 N21 O 23Na |
| | 710.3748 | -2.9 | -4.1 | 1.5 | 60.1 | 4.248 | 1.43 | C3 H40 N35 O9 |
| | 710.3738 | -1.9 | -2.7 | 15.5 | 60.1 | 4.249 | 1.43 | C33 H48 N11 O7 |
| | 710.3698 | 2.1 | 3.0 | 11.5 | 60.1 | 4.251 | 1.42 | C28 H48 N13 O9 |
| | 710.3743 | -2.4 | -3.4 | 8.5 | 60.2 | 4.344 | 1.30 | C18 H44 N23 O8 |
| | 710.3692 | 2.7 | 3.8 | 6.5 | 60.2 | 4.383 | 1.25 | C12 H41 N29 O6 23Na |
| | 710.3741 | -2.2 | -3.1 | 11.5 | 60.3 | 4.419 | 1.20 | C35 H53 N5 O9 23Na |
| | 710.3690 | 2.9 | 4.1 | -0.5 | 60.3 | 4.458 | 1.16 | C12 H48 N21 O14 |
| | 710.3746 | -2.7 | -3.8 | 4.5 | 60.4 | 4.512 | 1.10 | C20 H49 N17 O10 23Na |
| | 710.3689 | 3.0 | 4.2 | 10.5 | 60.4 | 4.541 | 1.07 | C10 H36 N35 O4 |
| | 710.3751 | -3.2 | -4.5 | 8.5 | 60.4 | 4.555 | 1.05 | C3 H33 N43 O 23Na |
| | 710.3746 | -2.7 | -3.8 | 15.5 | 60.4 | 4.575 | 1.03 | C18 H37 N31 23Na |
| | 710.3696 | 2.3 | 3.2 | 25.5 | 60.5 | 4.640 | 0.97 | C43 H45 N9 23Na |
| | 710.3693 | 2.6 | 3.7 | 18.5 | 60.7 | 4.816 | 0.81 | C43 H52 N O8 |
| | 710.3687 | 3.2 | 4.5 | 2.5 | 60.8 | 4.938 | 0.72 | C29 H57 N3 O15 23Na |
| | 710.3747 | -2.8 | -3.9 | 27.5 | 60.8 | 4.980 | 0.69 | C49 H48 N3 O2 |
| | 710.3687 | 3.2 | 4.5 | 13.5 | 60.8 | 4.989 | 0.68 | C27 H45 N17 O5 23Na |
| | 710.3685 | 3.4 | 4.8 | 6.5 | 60.9 | 5.081 | 0.62 | C27 H52 N9 O13 |
| | 710.3684 | 3.5 | 4.9 | 17.5 | 61.0 | 5.132 | 0.59 | C25 H40 N23 O3 |
| | 710.3752 | -3.3 | -4.6 | 9.5 | 61.0 | 5.136 | 0.59 | C36 H56 N O13 |
| | 710.3752 | -3.3 | -4.6 | 20.5 | 61.0 | 5.177 | 0.56 | C34 H44 N15 O3 |
| | 710.3754 | -3.5 | -4.9 | 16.5 | 61.2 | 5.318 | 0.49 | C36 H49 N9 O5 23Na |

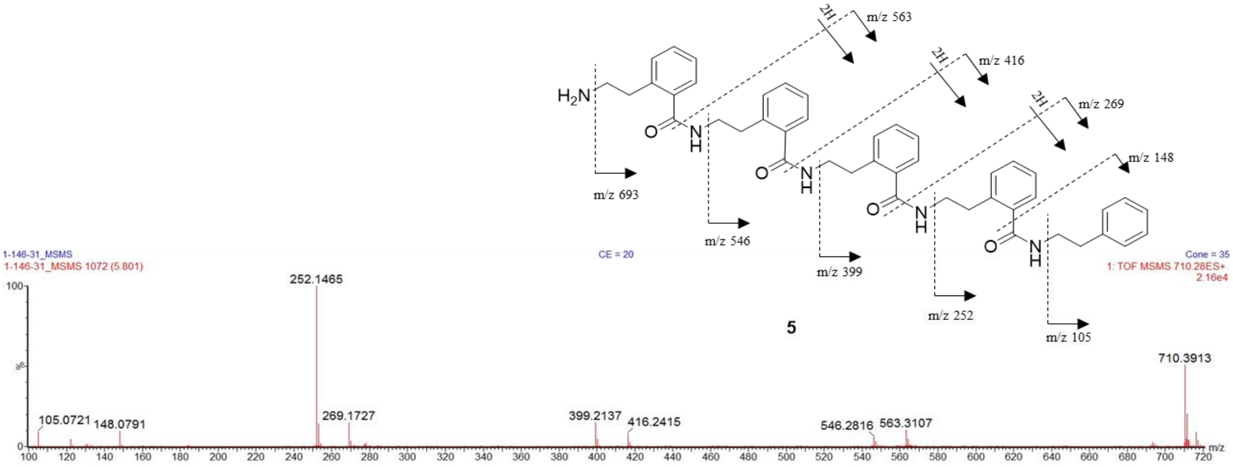


**Figure 1-figure supplement 21**. (+)-HRESITOFMS of compound **5** (top) with predicted elemental composition. (+)-CID MS/MS (20 eV) of *m/z* 710 showing a distinct fragmentation pattern (bottom).

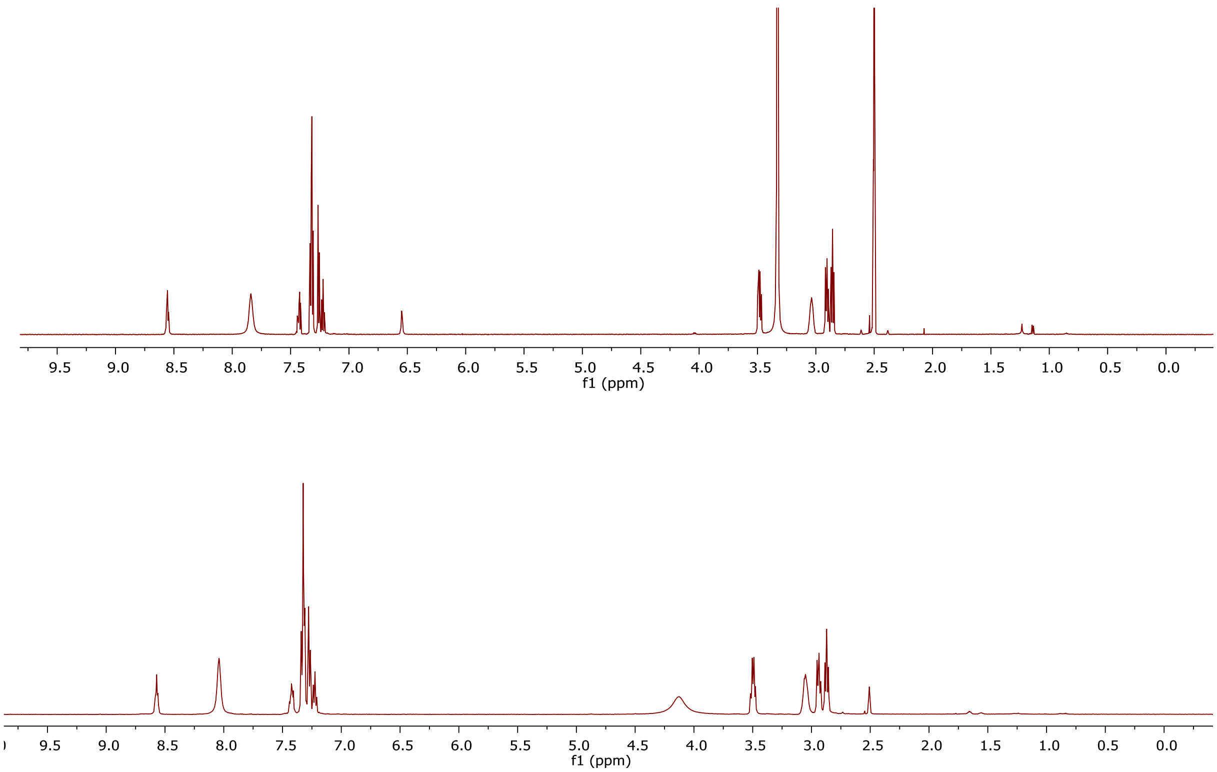


**Figure 1- figure supplement 1**. $^{1}$H NMR (500 MHz) spectrum of **1** (natural; top) and synthetic **1** (bottom) in DMSO-$d_6$.

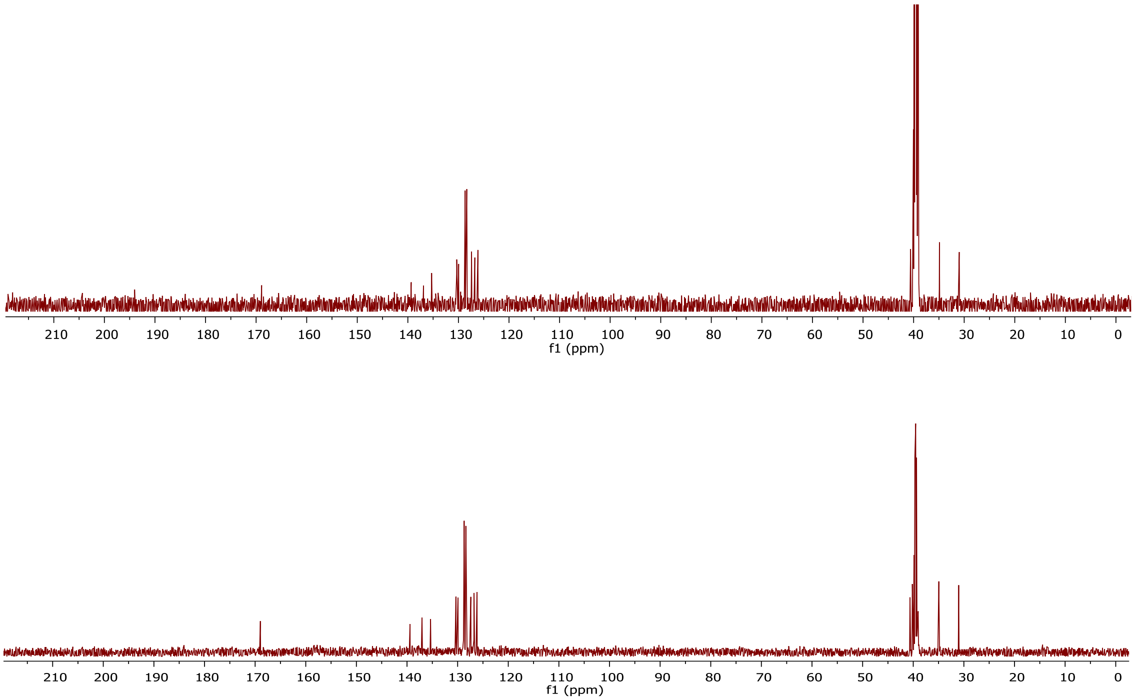


**Figure 1- figure supplement 23**. $^{13}$C NMR (125 MHz) spectrum of **1** (natural; top) and synthetic **1** (bottom) in DMSO-$d_6$.

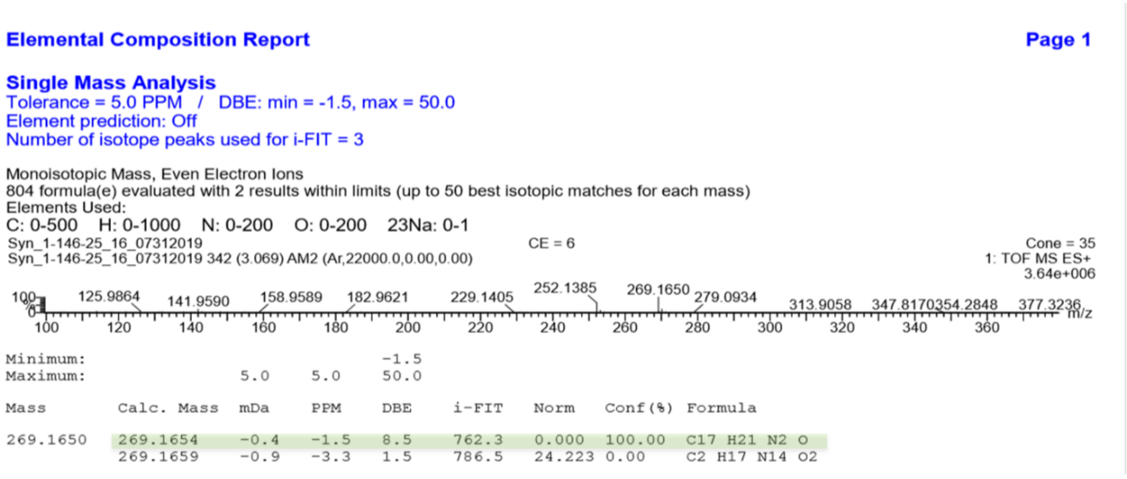

**Elemental Composition Report** **Page 1**

**Single Mass Analysis**
Tolerance = 5.0 PPM / DBE: min = -1.5, max = 50.0
Element prediction: Off
Number of isotope peaks used for i-FIT = 3

Monoisotopic Mass, Even Electron Ions
804 formula(e) evaluated with 2 results within limits (up to 50 best isotopic matches for each mass)
Elements Used:
C: 0-500 H: 0-1000 N: 0-200 O: 0-200 23Na: 0-1
Syn_1-146-25_16_07312019 CE = 6 Cone = 35
Syn_1-146-25_16_07312019 342 (3.069) AM2 (Ar,22000.0,0.00,0.00) 1: TOF MS ES+ 3.64e+006



| Mass | Calc. Mass | mDa | PPM | DBE | i-FIT | Norm | Conf(%) | Formula |
|---|---|---|---|---|---|---|---|---|
| Minimum: | | | | -1.5 | | | | |
| Maximum: | | 5.0 | 5.0 | 50.0 | | | | |
| 269.1650 | 269.1654 | -0.4 | -1.5 | 8.5 | 762.3 | 0.000 | 100.00 | C17 H21 N2 O |
| | 269.1659 | -0.9 | -3.3 | 1.5 | 786.5 | 24.223 | 0.00 | C2 H17 N14 O2 |

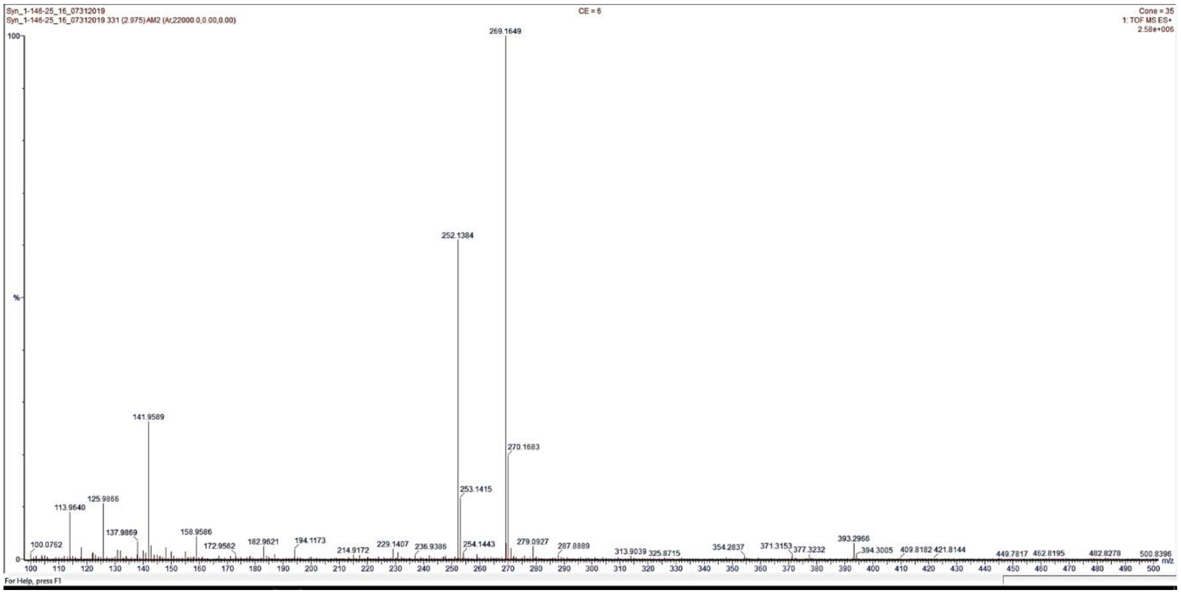



**Figure 1- figure supplement 24**. (+)-HRESITOFMS of synthesized compound **1** with predicted elemental composition.

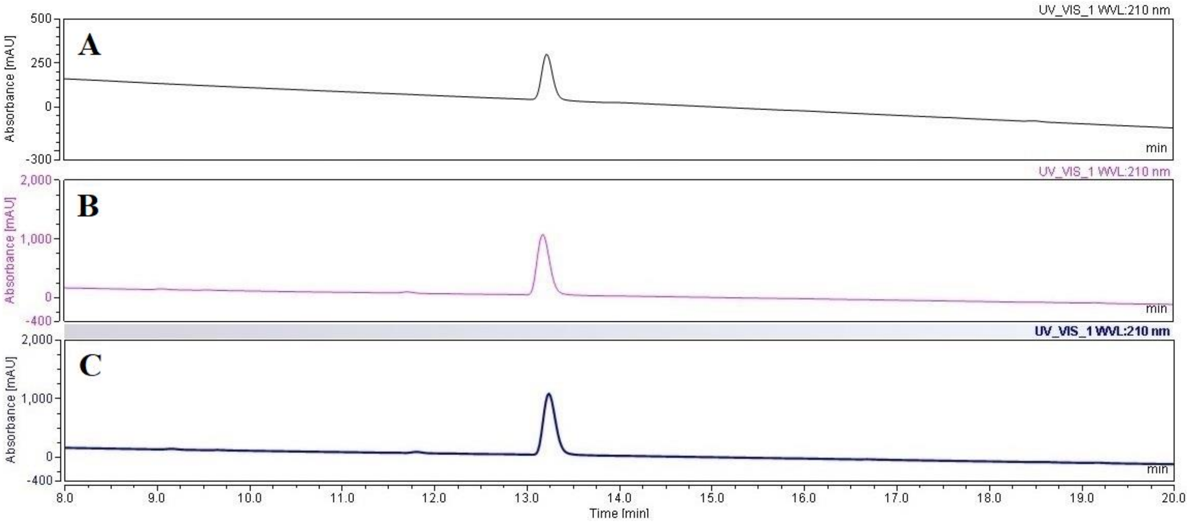


**Figure 1- figure supplement 5**. Chromatographic co-elution profile of natural and synthetic **1**. The HPLC runs were carried out using an analytical Eclipse Plus $C_{18}$ column (3.5 µm; 4.6 x 150 mm) and eluted with a linear gradient from 5% to 100% $CH_3CN$ in $H_2O$ (0.1% TFA) over 24 min at 0.5 mL/min flow rate. (**A**) Compound **1** isolated from *P. forskalii*. (**B**) Synthetic **1**. (**C**) Mixture of natural and synthetic **1**. The UV absorbance was monitored at 210 nm.

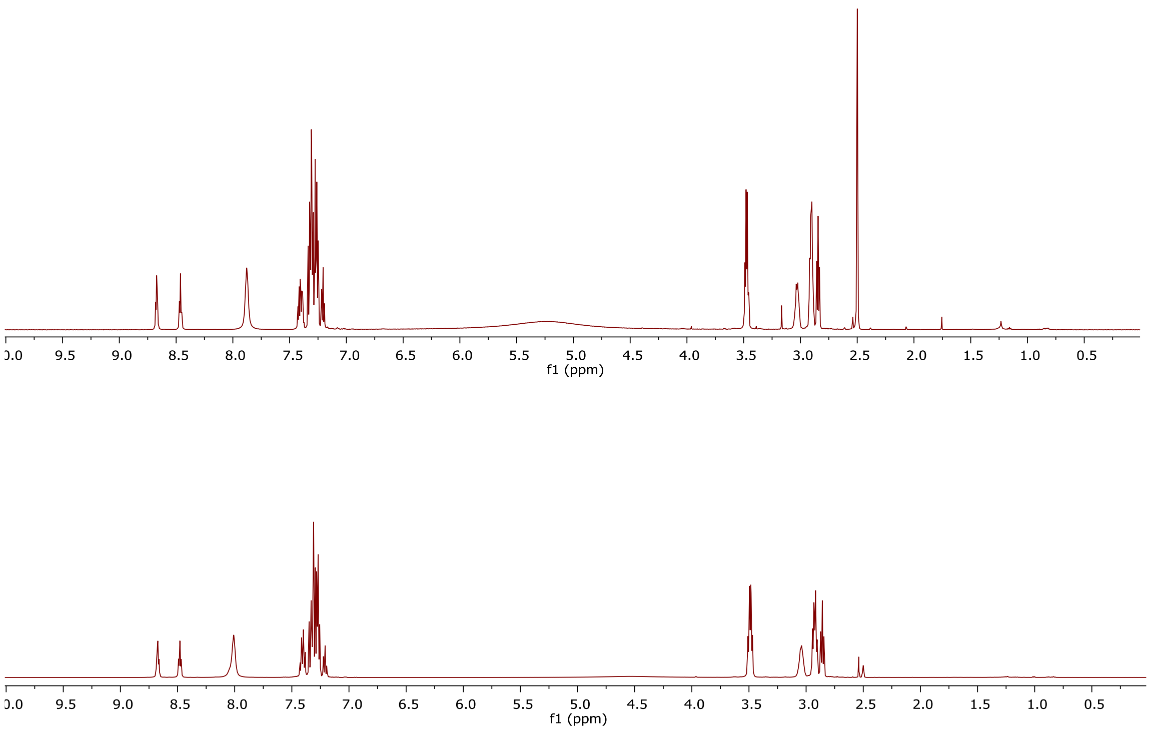


**Figure 1- figure supplement 2**. $^{1}H$ NMR (500 MHz) spectrum of **3** (natural; top) and synthetic **3** (bottom) in DMSO-$d_6$.

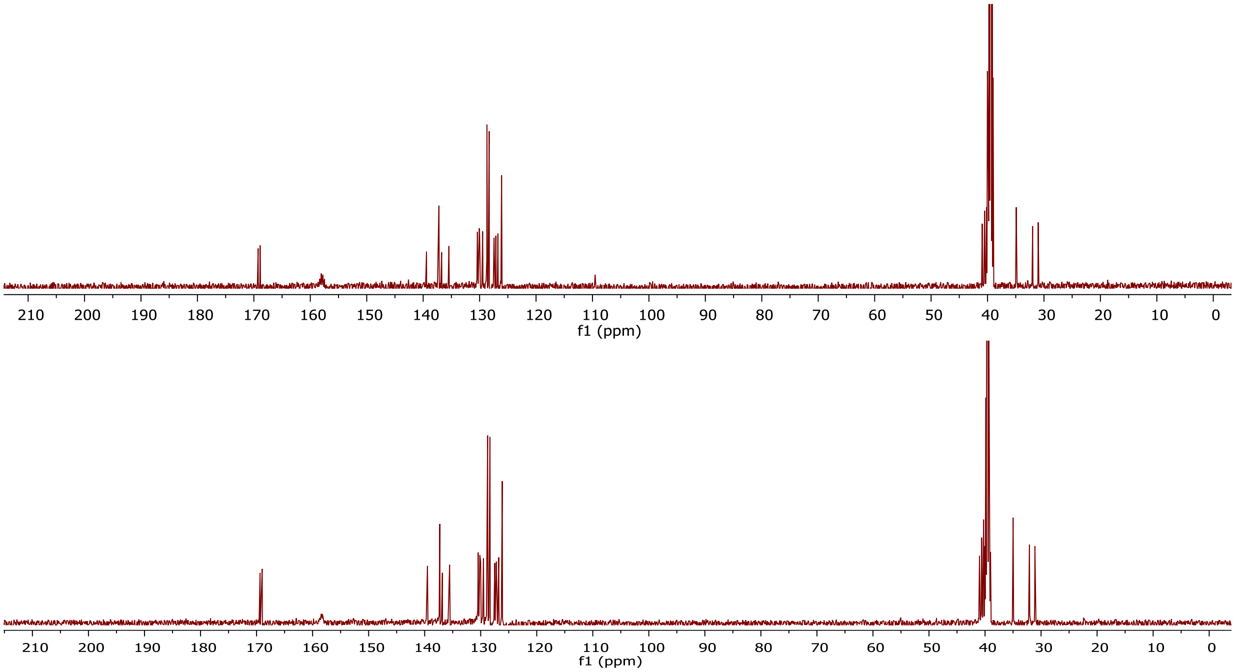


**Figure 1- figure supplement 27**. $^{13}C$ NMR (125 MHz) spectrum of **3** (top) and synthetic **3** (bottom) in DMSO-$d_6$.

### Elemental Composition Report

**Page 1**

#### Single Mass Analysis

Tolerance = 5.0 PPM / DBE: min = -1.5, max = 50.0
Element prediction: Off
Number of isotope peaks used for i-FIT = 3

Monoisotopic Mass, Even Electron Ions
2543 formula(e) evaluated with 7 results within limits (up to 50 best isotopic matches for each mass)
Elements Used:
C: 0-500 H: 0-1000 N: 0-200 O: 0-200 23Na: 0-1

Syn_1-146-28_34_07312019_4 CE = 6 Cone = 35
Syn_1-146-28_34_07312019_4 530 (4.710) AM2 (Ar,22000.0,0.00,0.00) 1: TOF MS ES+
1.55e+006

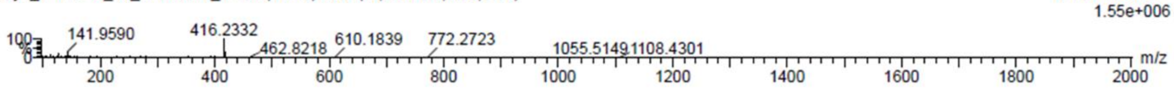


| Mass | Calc. Mass | mDa | PPM | DBE | i-FIT | Norm | Conf(%) | Formula |
|---|---|---|---|---|---|---|---|---|
| Minimum: | | | | -1.5 | | | | |
| Maximum: | | 5.0 | 5.0 | 50.0 | | | | |
| 416.2332 | 416.2338 | -0.6 | -1.4 | 13.5 | 567.5 | 0.000 | 99.96 | C26 H30 N3 O2 |
| | 416.2314 | 1.8 | 4.3 | 10.5 | 575.4 | 7.840 | 0.04 | C24 H31 N3 O2 23Na |
| | 416.2346 | -1.4 | -3.4 | 2.5 | 583.2 | 15.701 | 0.00 | C13 H31 N9 O5 23Na |
| | 416.2330 | 0.2 | 0.5 | 1.5 | 585.2 | 17.715 | 0.00 | C10 H30 N11 O7 |
| | 416.2343 | -1.1 | -2.6 | 6.5 | 585.5 | 18.017 | 0.00 | C11 H26 N15 O3 |
| | 416.2319 | 1.3 | 3.1 | 3.5 | 587.4 | 19.910 | 0.00 | C9 H27 N15 O3 23Na |
| | 416.2316 | 1.6 | 3.8 | 7.5 | 589.5 | 21.950 | 0.00 | C7 H22 N21 O |

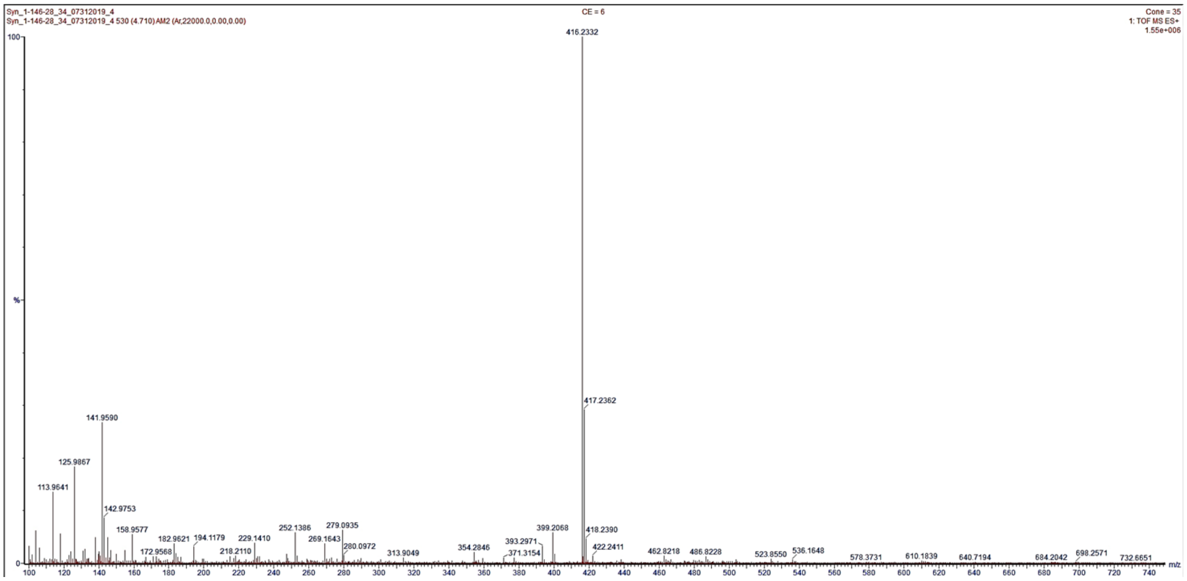


**Figure 1- figure supplement 28**. (+)-HRESITOFMS of synthesized compound **3** with predicted elemental composition.

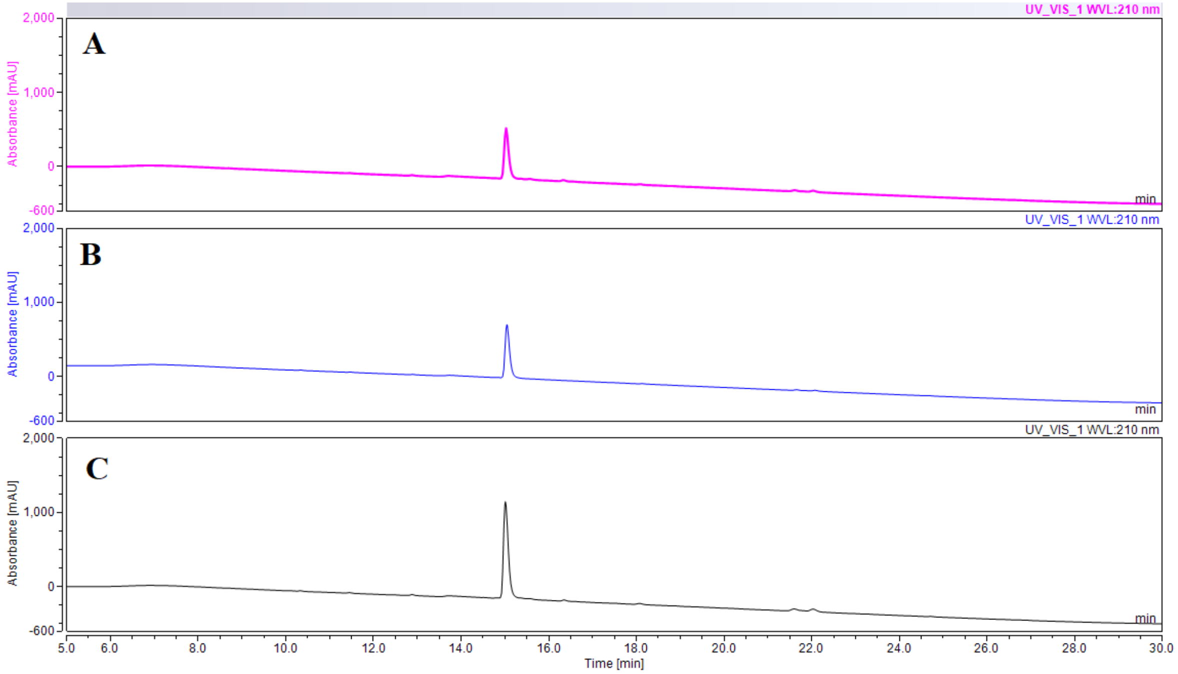


**Figure 1- figure supplement 29**. Chromatographic co-elution profile of natural and synthetic compound **3**. The HPLC runs were carried out using an analytical Eclipse Plus C18 column (3.5 µm; 4.6 x 150 mm) and eluted with a linear gradient from 5% to 100% CH3CN in H2O (0.1% TFA) over 24 min at 0.5 mL/min flow rate. (**A**) Compound **3** isolated from *P. forskalii*. (**B**) Synthetic **3**. (**C**) Mixture of natural and synthetic **3**. The UV absorbance was monitored at 210 nm.

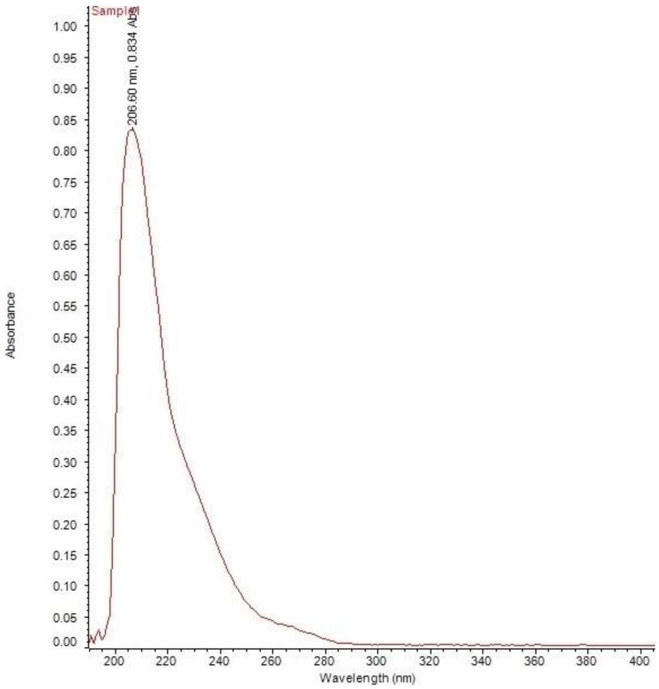


**Figure 1- figure supplement 30**. UV spectrum of **1** in $CH_3OH$.

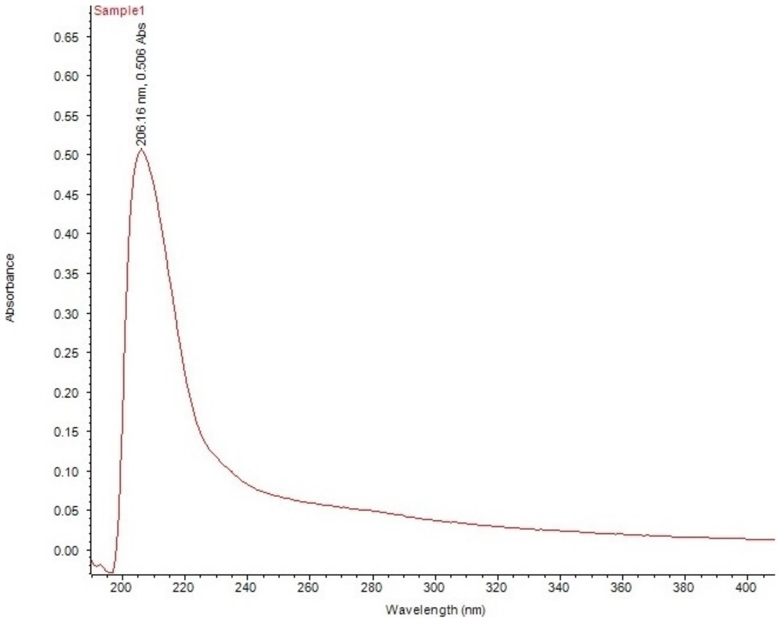


**Figure 1- figure supplement 31**. UV spectrum of **2** in $CH_3OH$.

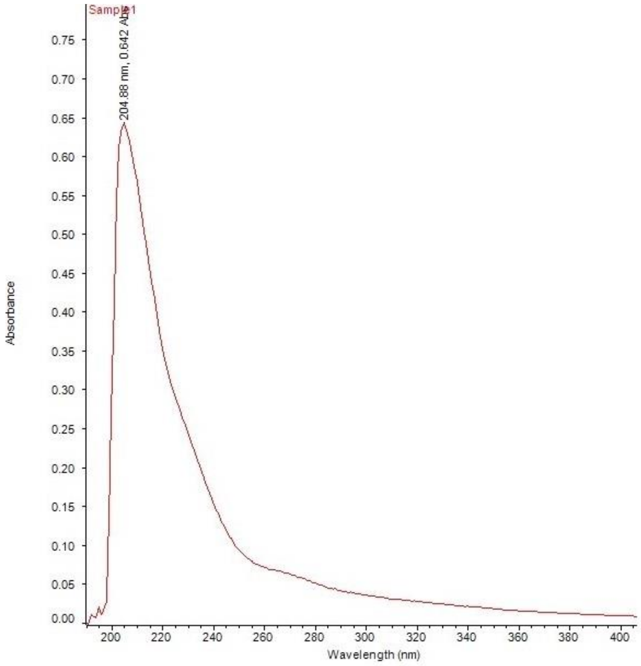


**Figure 1- figure supplement 32**. UV spectrum of **3** in $CH_3OH$.

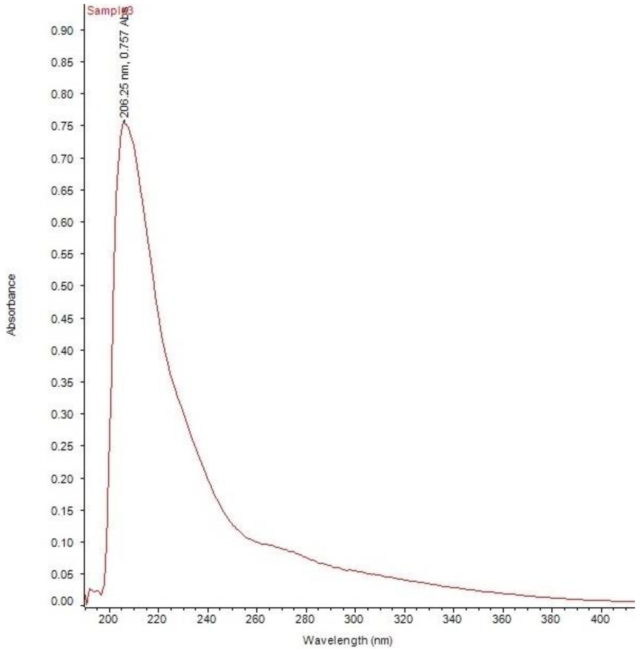


**Figure 1- figure supplement 33**. UV spectrum of **4** in $CH_3OH$.

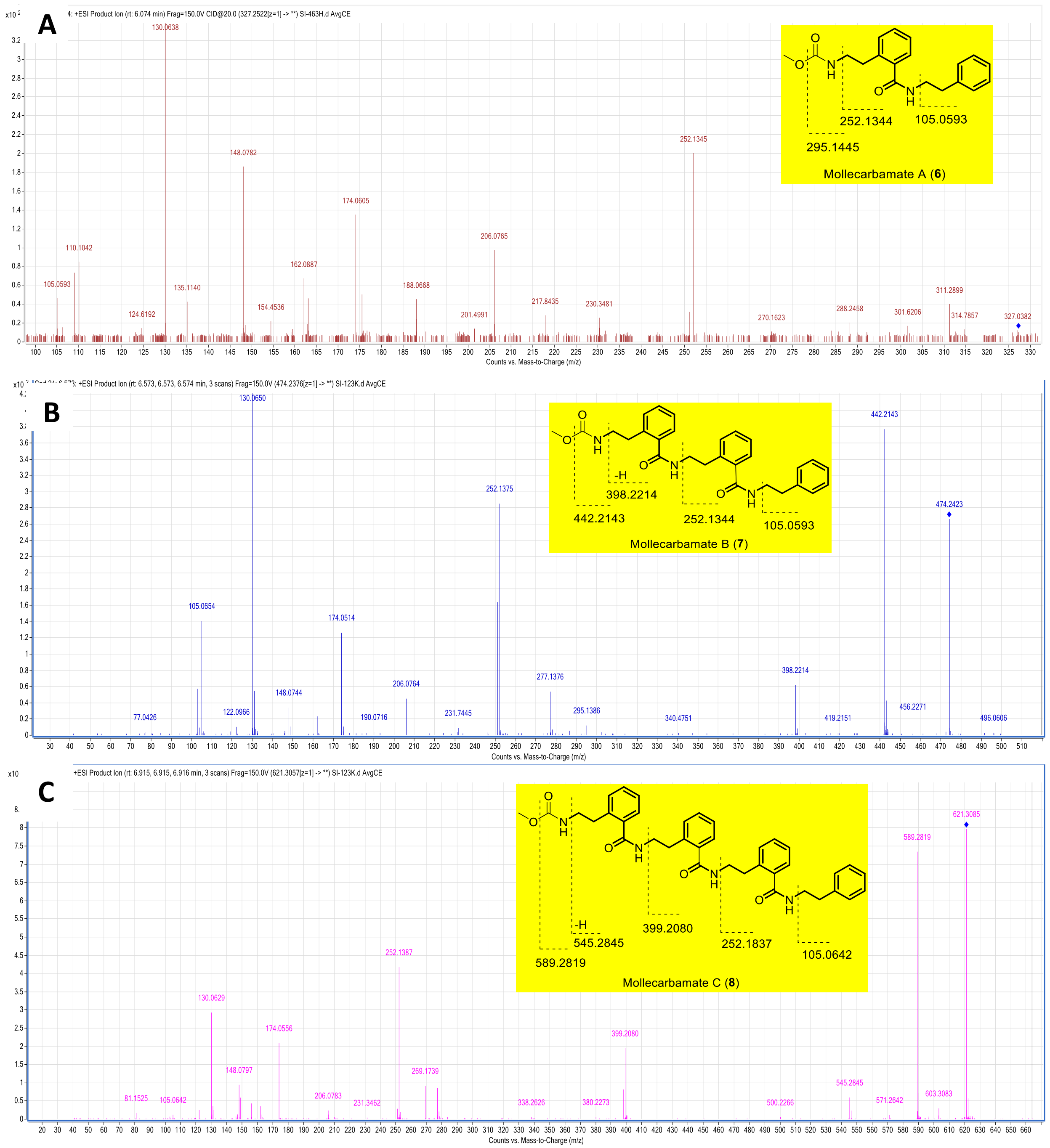


**Figure 2-figure supplement 1**. Chemical structure and MS/MS fragmentation of compounds structurally-related to molleamines identified by metabolomics analysis of *D. molle* specimens. (**A**) Mollecarbamate A. (**B**) Mollecarbamate B. (**C**) Mollecarbamate C.

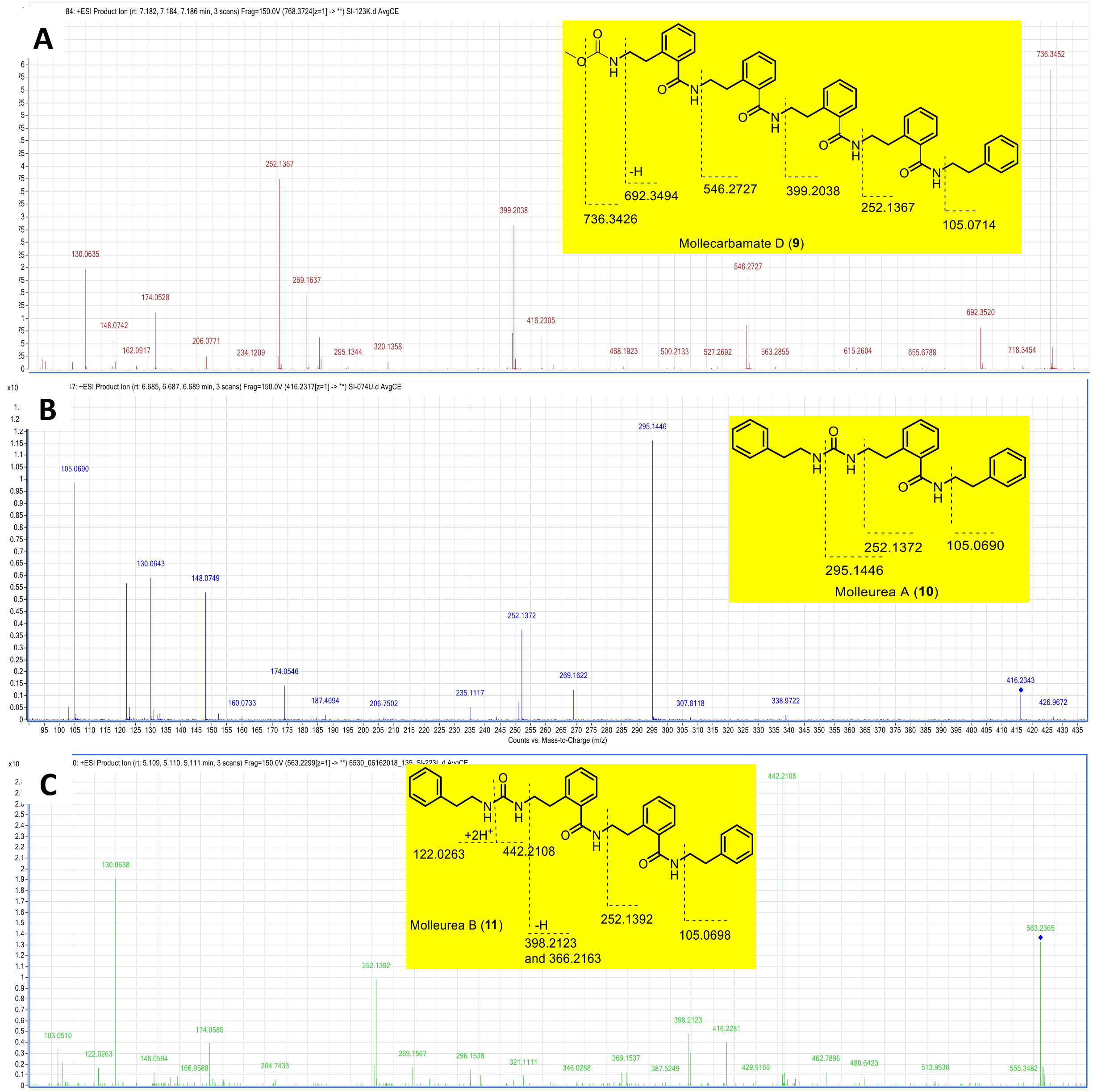


**Figure 2-figure supplement 2**. Chemical structure and MS/MS fragmentation of compounds structurally related to molleamines identified by metabolomics analysis of *D. molle* specimens. (**A**) Mollecarbamate D. (**B**) Molleurea A. (**C**) Molleurea B.

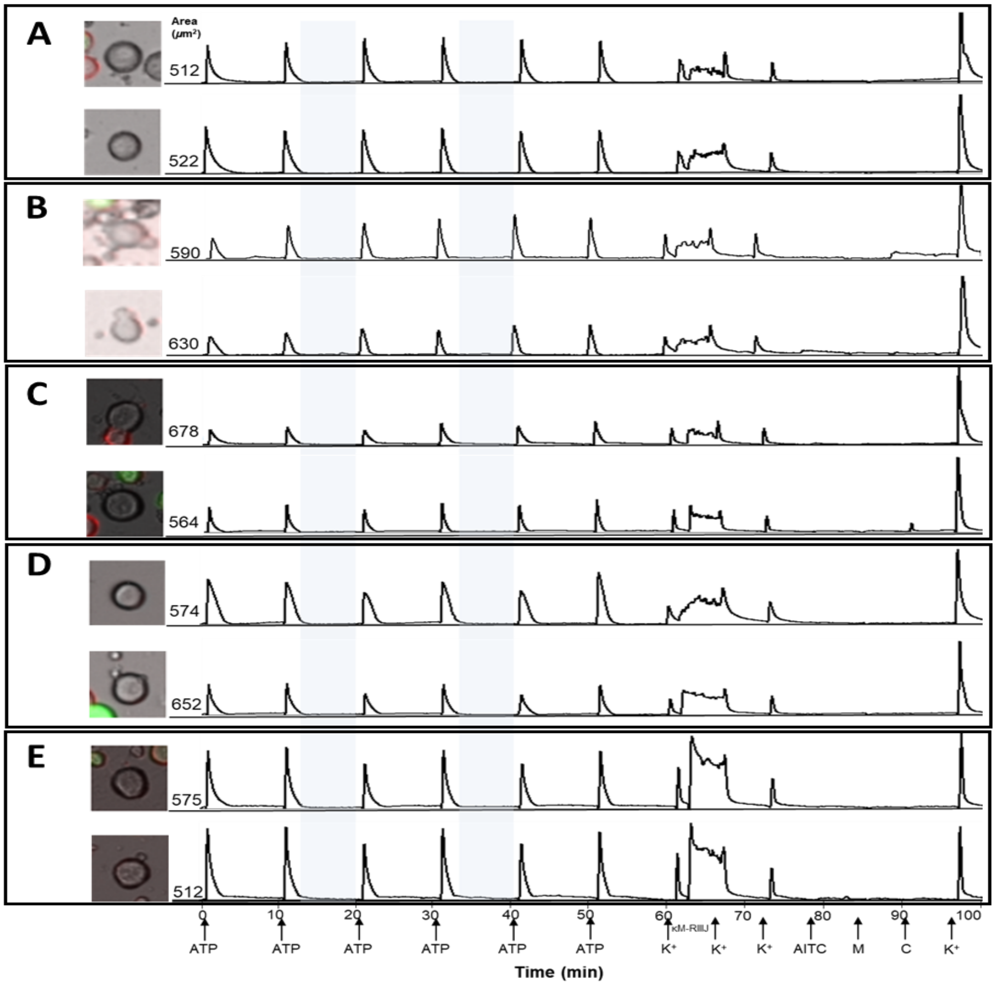


**Figure 3-figure supplement 1**. Constellation pharmacology indicates that compounds **3-5** are selective against L2 DRG neurons, which are Aδ-low threshold mechanoreceptors (LTMRs). Shown above are selected calcium imaging traces from dissociated DRG neurons. Left shows the bright-field image of the cells. The numbers indicate the cross-sectional area of each cell in micrometers$^2$. The y-axis and x-axis (time in minutes) correspond to the relative fluorescence and individual application of treatments for a 15 s duration, respectively. ATP (20 µM) was used to evoke intracellular $Ca^{2+}$ depolarization. Shaded regions indicate the application of the compound (20 µM) in the observation solution. Pharmacological identifiers were sequentially applied toward the end of the experiment to identify different subtypes of DRG neurons including 20 mM KCl ($K^+$), 1 µM κM-RIIIJ, 100 µM allyl isothiocyanate (AITC), 400 µM menthol (M), and 300 nM capsaicin (C). At the end of each experiment, 40 mM of $K^+$ was applied. Responses from L2 neurons that were unaffected following incubation with **(A)** compound **1** and **(B)** compound **2**. Representative traces from L2 neurons showing attenuated responses to depolarization with ATP following incubation with **(B)** compound **3**, **(C)** compound **4**, and **(D)** compound **5**.

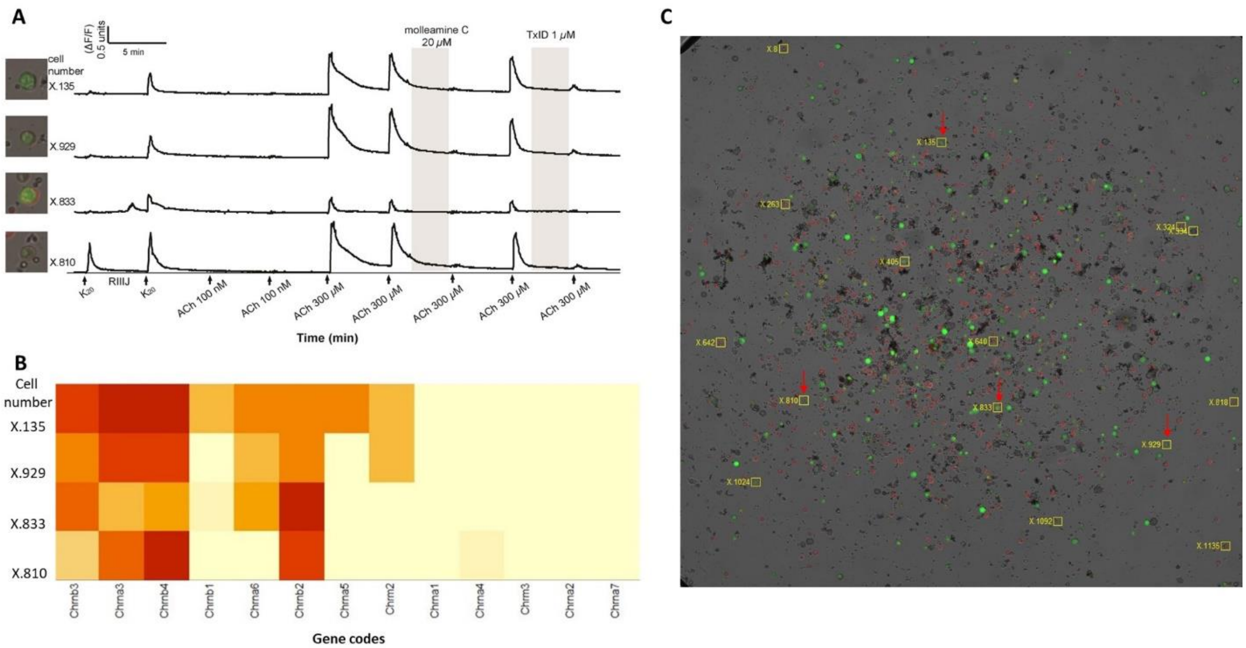


**Figure 3-figure supplement 2.** Constellation pharmacology of molleamine C and transcriptomics analysis of neurons with responses to acetylcholine (ACh) blocked by molleamine C (20 µM) and TxID (1 µM). **(A)** Neurons were classified by the fluorescent labeling, response to potassium ($K_{20,}$ 20 mM) and the response to a 4-minute application of RIIIJ (1 µM). Two applications of acetylcholine (ACh,100 nM) for ~15 s were applied followed by two consecutive ACh (300 µM) were applied at five-minute intervals. In between these applications a control (DRG observation buffer) or compound (**3** at 20 µM or TxID at 1 µM) was applied. **(B)** Neurons shown in panel A were collected, and transcriptomic analysis was done according to procedure described in Methods Section. The heatmap is a representation of the single-cell transcriptomic *CHRN* gene expression where each row is an individual neuron, and each column is a *CHRN* subunit. The heatmap is color-coded based on the level of expression on a logarithmic scale. Darker red colors represent high expression, lighter colors represent lower expression. Trancriptomic data revealed enhanced expression of transcripts for α3(*CHRNA3*), β3(*CHRNB3*), β4(*CHRNA3*), β2(*CHRNB2*) and α6(*CHRNA6*) nicotinic acetylcholine receptor subunits. **(C)** DRG neurons in calcium imaging experiment where the bright-field image is overlaid with fluorescent images obtained from IB4 (red) staining and GFP expression (green). Neurons with ACh response blocked by molleamine C and TxID and analyzed in single-cell transcriptomics are shown and indicated by a red arrow.

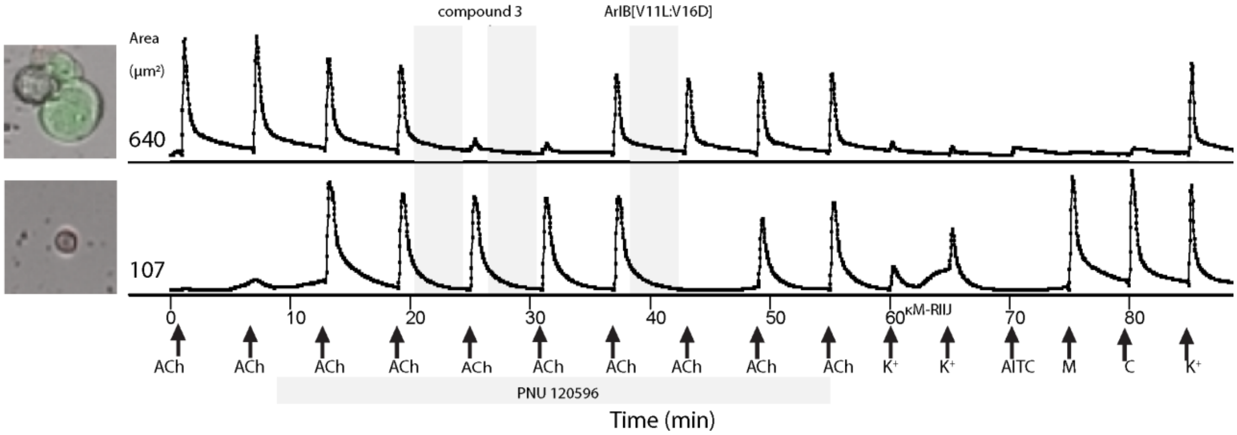


**Figure 3-figure supplement 3**. Molleamine C does not affect α7-nAChRs. The neuron at the top responded to 1 mM ACh before the application of PNU 120596 (10 μM) and served as control for the experiment. The neuron at the bottom responded to ACh only after pre-incubation with PNU 120596, suggesting that it expressed α7-nAChRs. The PNU 120596-enhanced and ACh-induced response from this cell was not blocked by molleamine C (20 μM) but blocked by the selective antagonist of α7-nAChRs, ArIB[V11L;V16D] at 200 nM.

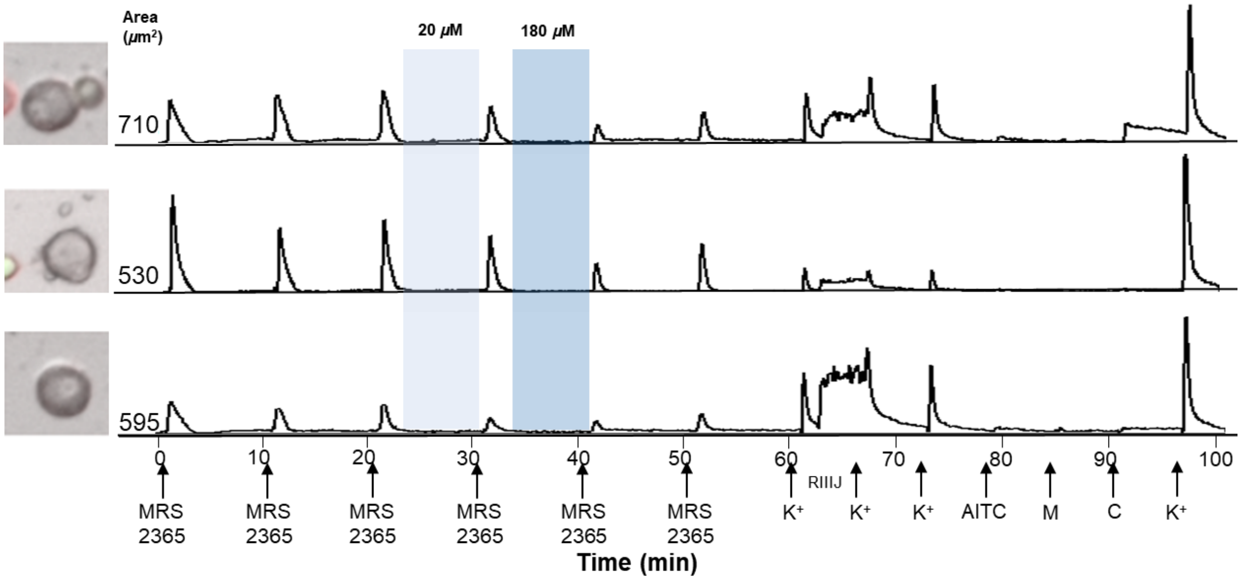


**Figure 6-figure supplement 1**. Compound **3** inhibits responses from L2 neurons after depolarization with a P2Y1 agonist MRS 2365 (100 nM).

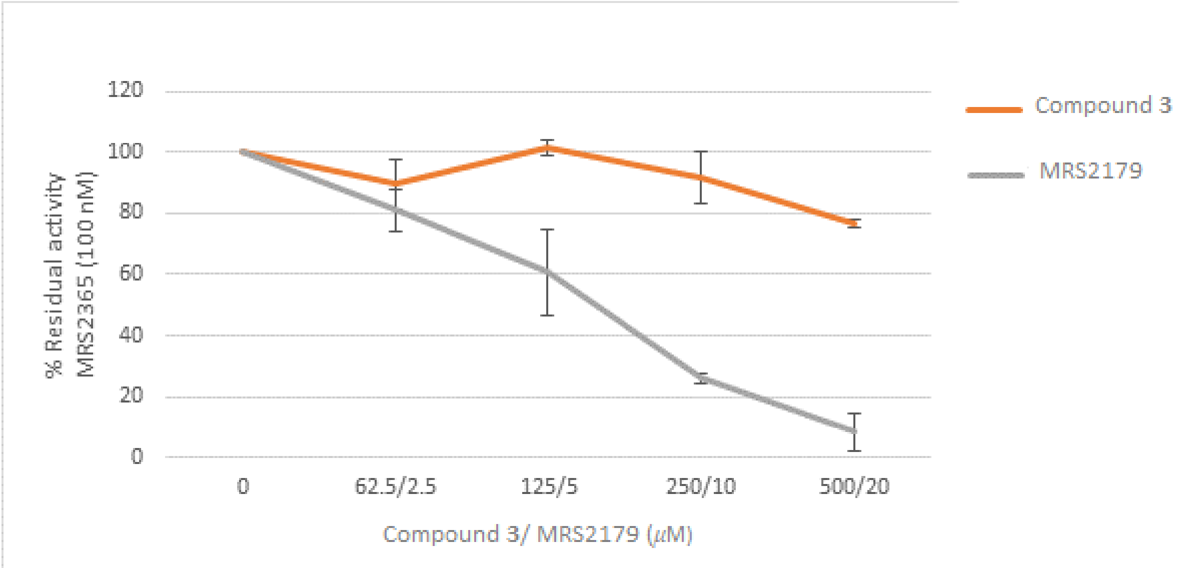


**Figure 6-figure supplement 2**. Compound **3** does not inhibit P2Y1 activity in human embryonic kidney (HEK-293) over-expressing GCaMP6s. Treatment wells were incubated with the P2Y1 antagonist MRS2179 or molleamine C 30 min before assesing P2Y1 activity in the calcium imaging experiments. Data are represented as a percent residual activity of the agonist MRS2365 (100 nM).

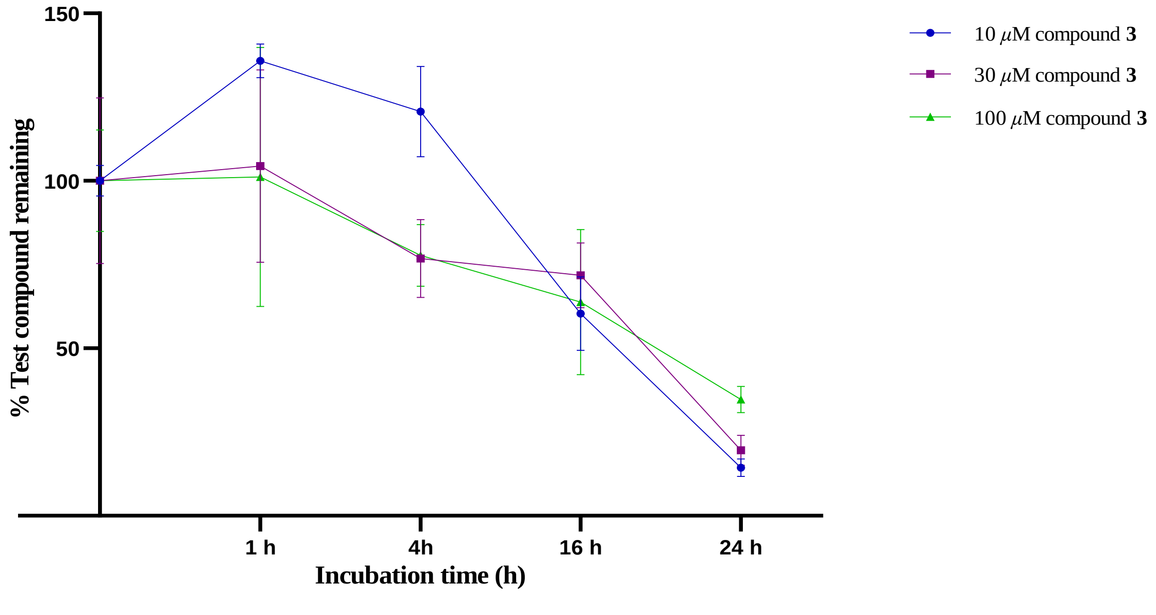


**Figure 7-figure supplement 1**. Stability of compound **3** in mouse plasma over 24 h (data represent mean percentage test compound remaining against time with the error bars representing the standard deviation of three replicate reactions).

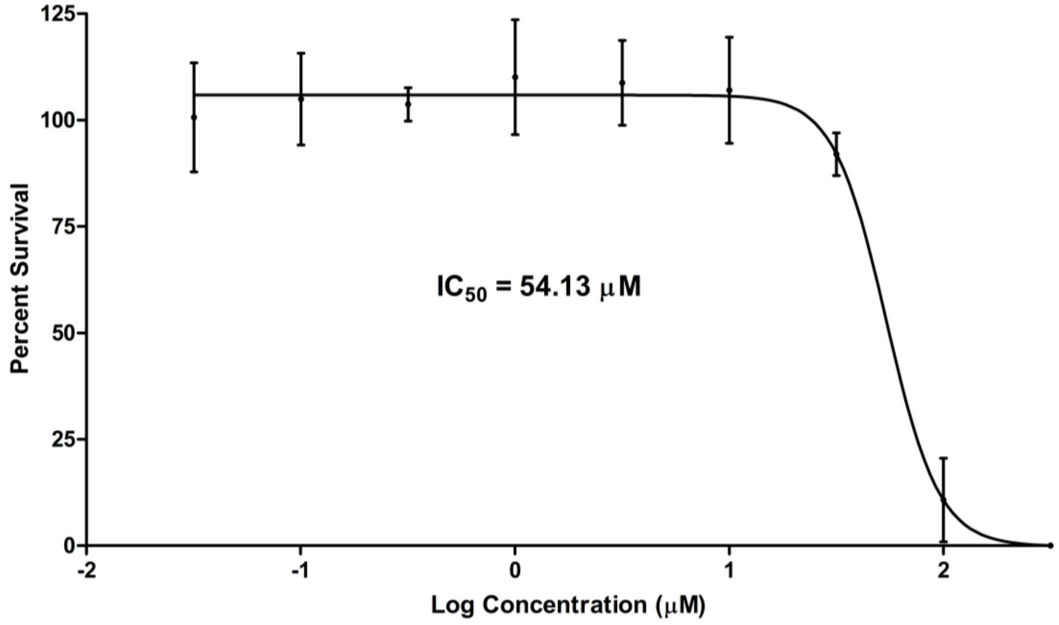


**Figure 7-figure supplement 2**. Cytotoxicity evaluation of compound **3** in *in vitro* MTT assay with human embryonic kidney (HEK-293) cells. The $IC_{50}$ value was calculated using GraphPad Prism 9.0.0 based on a four-point sigmoidal nonlinear regression analysis of cell viability expressed as % cell survival vs log concentration of **3**.

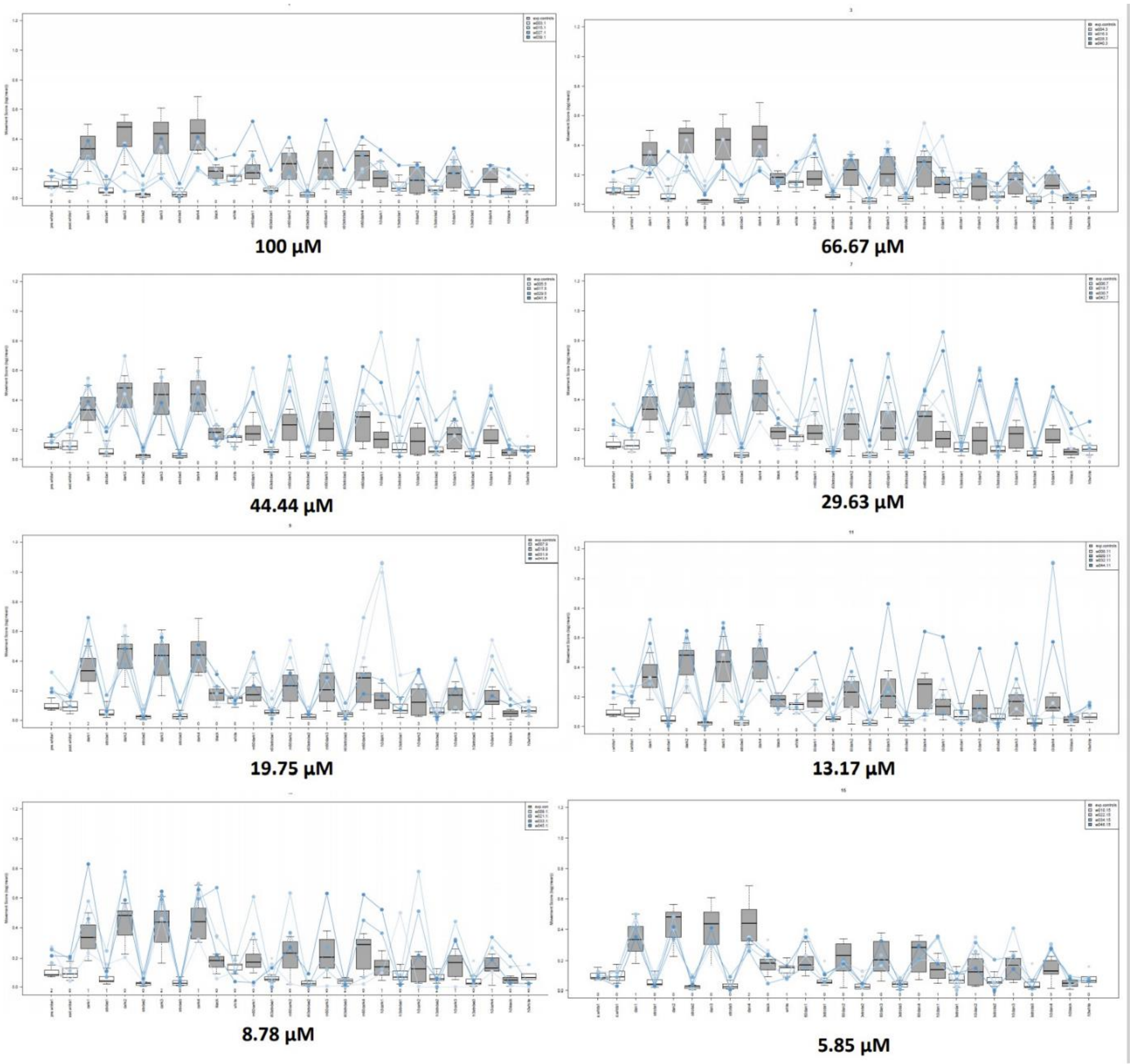


**Figure 7-figure supplement 3**. Zebrafish photomotor-response after exposure to increasing concentrations of compound **3**.

**Figure 1-table supplement 1.** $^1$H NMR (400 MHz) and $^{13}$C NMR (125 MHz) data for compounds **1-5** in DMSO-$d_6$.

| position | **1** | | **2** | | **3** | | **4** | | **5** | |
|---|---|---|---|---|---|---|---|---|---|---|
| | $\delta_C$, type | $\delta_H$, mult. ($J$ in Hz) | $\delta_C$, type | $\delta_H$, mult. ($J$ in Hz) | $\delta_C$, type | $\delta_H$, mult. ($J$ in Hz) | $\delta_C$, type | $\delta_H$, mult. ($J$ in Hz) | $\delta_C$, type | $\delta_H$, mult. ($J$ in Hz) |
| Subunit A | | | | | | | | | | |
| 1 | 40.2, $CH_2$ | 3.03, m | 40.2, $CH_2$ | 3.04, m | 40.2, $CH_2$ | 3.03, m | 40.2, $CH_2$ | 3.01, m | 40.2, $CH_2$ | 3.02, m |
| 2 | 31.0, $CH_2$ | 2.91, t (7.8) | 31.0, $CH_2$ | 2.91, t (7.8) | 31.0, $CH_2$ | 2.91, m | 31.0, $CH_2$ | 2.90, m | 31.0, $CH_2$ | 2.89, m |
| 3 | 136.9, C | | 137.0, C | | 137.2, C | | 137.3, C | | 137.3, C | |
| 4 | 126.8, CH | 7.31, m | 130.3, CH | 7.32, m | 126.8, CH | 7.31, m | 126.8, CH | 7.30[a] | 127.4, CH | 7.30[a] |
| 5 | 130.0, CH | 7.43, m | 130.0, CH | 7.43, m | 130.0, CH | 7.43, m | 130.0, CH | 7.40, m | 130.1, CH | 7.41 td (7.2,2.1) |
| 6 | 127.4, CH | 7.31, m | 126.8, CH | 7.32, m | 127.4, CH | 7.31, m | 127.4, CH | 7.30[a] | 126.8, CH | 7.25[a] |
| 7 | 130.3, CH | 7.34, m | 127.4, CH | 7.32, m | 130.4, CH | 7.32, m | 130.4, CH | 7.30[a] | 128.3, CH | 7.30[a] |
| 8 | 135.4, C | | 135.3, C | | 136.7, C | | 137.2, C | | 137.3, C | |
| 9 | 168.9, C | | 169.1, C | | 169.2, C | | 169.3, C | | 169.2, C | |
| $NH_2$ | | 7.84, brs | | 7.84, brs | | 7.88, brs | | 7.84, brs | | 7.81, brs |
| Subunit B1 | | | | | | | | | | |
| 1 | | | | | 40.9, $CH_2$ | 3.47, td (7.3, 5.2) | 40.9, $CH_2$ | 3.47, m | 41.0, $CH_2$ | 3.46, m |
| 2 | | | | | 32.0, $CH_2$ | 2.90, t (7.3) | 32.0, $CH_2$ | 2.90, m | 32.0, $CH_2$ | 2.89, m |
| 3 | | | | | 137.2, C | | 137.2, C | | 137.3, C | |
| 4 | | | | | 126.1, CH | 7.30, m | 126.1, CH | 7.30[a] | 127.1, CH | 7.30[a] |
| 5 | | | | | 129.4, CH | 7.40, m | 129.5, CH | 7.40[a] | 129.5, CH | 7.39[a] |
| 6 | | | | | 127.1, CH | 7.31, m | 127.2, CH | 7.30[a] | 126.1, CH | 7.25[a] |
| 7 | | | | | 130.1, CH | 7.31, m | 130.1, CH | 7.30[a] | 130.0, CH | 7.30[a] |
| 8 | | | | | 135.4, C | | 136.7, C | | 137.2, C | |
| 9 | | | | | 168.9, C | | 169.3, C | | 169.2, C | |
| NH | | | | | | 8.68, t (5.2) | | 8.68, t (4.8) | | 8.68, t (5.1) |
| Subunit B2 | | | | | | | | | | |
| 1 | | | | | | | 40.8, $CH_2$ | 3.47, m | 40.8, $CH_2$ | 3.46, m |
| 2 | | | | | | | 32.0, $CH_2$ | 2.90, m | 32.0, $CH_2$ | 2.89, m |
| 3 | | | | | | | 137.0, C | | 137.2, C | |
| 4 | | | | | | | 126.1, CH | 7.30[a] | 127.2, CH | 7.30[a] |
| 5 | | | | | | | 129.4, CH | 7.40[a] | 129.4, CH | 7.39[a] |
| 6 | | | | | | | 127.1, CH | 7.30[a] | 126.1, CH | 7.25[a] |
| 7 | | | | | | | 130.1, CH | 7.30[a] | 130.4, CH | 7.30[a] |
| 8 | | | | | | | 135.4, C | | 137.0, C | |
| 9 | | | | | | | 168.9, C | | 169.2, C | |
| NH | | | | | | | | 8.57, t (5.2) | | 8.57, t (5.1) |
| Subunit B3 | | | | | | | | | | |
| 1 | | | | | | | | | 40.8, $CH_2$ | 3.46, m |
| 2 | | | | | | | | | 32.0, $CH_2$ | 2.89, m |
| 3 | | | | | | | | | 137.0, C | |
| 4 | | | | | | | | | 128.7, CH | 7.25[a] |
| 5 | | | | | | | | | 129.4, CH | 7.39[a] |
| 6 | | | | | | | | | 126.1, CH | 7.25[a] |
| 7 | | | | | | | | | 130.1, CH | 7.30[a] |
| 8 | | | | | | | | | 135.4, C | |
| 9 | | | | | | | | | 168.9, C | |
| NH | | | | | | | | | | 8.54, t (5.1) |
| Subunit C | | | | | | | | | | |
| 1 | 40.6 $CH_2$ | 3.48, td (7.4, 5.5) | 41.0, $CH_2$ | 3.41, m | 40.5, $CH_2$ | 3.47 td (7.3, 5.5) | 40.5, $CH_2$ | 3.47, m | 40.5, $CH_2$ | 3.46. m |
| 2 | 34.9, $CH_2$ | 2.86, t (7.4) | 34.2, $CH_2$ | 2.73, t (7.3) | 34.9, $CH_2$ | 2.84 t (7.3) | 34.9, $CH_2$ | 2.83, t (7.2) | 34.9, $CH_2$ | 2.83, t (7.3) |
| 3 | 139.4, C | | 129.5, C | | 139.4, C | | 139.4, C | | 139.4, C | |
| 4 | 128.7, CH | 7.26, d (7.2) | 129.1, CH | 7.04, d (8.4) | 128.3, CH | 7.26, m | 128.7, CH | 7.26[a] | 128.7, CH | 7.25[a] |
| 5 | 128.4, CH | 7.32, m | 114.9, CH | 6.70, d (8.4) | 128.7, CH | 7.25, m | 128.3, CH | 7.25[a] | 128.3, CH | 7.29, d (7.5) |
| 6 | 126.2, CH | 7.22, t (7.2) | 155.7, C | | 126.1, CH | 7.21, t (7.7) | 126.1, CH | 7.20, t (7.2) | 126.1, CH | 7.20 brt (7.30) |
| 7 | 128.4, CH | 7.32, m | 114.9, CH | 6.70, d (8.4) | 128.7, CH | 7.25, m | 128.3, CH | 7.25[a] | 128.3, CH | 7.29, d (7.5) |
| 8 | 128.7, CH | 7.26, d (7.2) | 129.1, CH | 7.04, d (8.4) | 128.3, CH | 7.26, m | 128.7, CH | 7.26[a] | 128.7, CH | 7.25[a] |
| NH | | 8.55, t (5.5) | | 8.51, t (5.7) | | 8.46, t (5.5) | | 8.44, t (5.2) | | 8.43, t (5.7) |
| 6-OH | | | | 9.21, bs | | | | | | |

[a]overlapping signals; chemical shifts were determined from $^1$H-$^{13}$C HSQC correlations.

**Figure 3-table supplement 1**. Census of effects elicited by compounds **1-5** on ATP-induced depolarization in 16 DRG neuronal subtypes screened in calcium imaging experiments. Intracellular calcium ($Ca^{2+}$) depolarization was induced by application of 20 µM ATP. The ability of the compounds (at 20 µM) to modulate the $Ca^{2+}$ signal was scored as either a block (**B**; attenuation of the signal), or amplification (**A**) immediately following compound incubation. Direct effects (**DE**) to the $Ca^{2+}$ baseline with sample incubation were also scored. Each compound was tested twice (except for compound **2**) and for each calcium imaging experiment the compound was applied twice. Only reproducible effects elicited by the compound were scored in each experiment. The effects for each subtype are represented as percentages (number of cells affected/total number of cells x 100) on the table.

| | Neurons | | | L1 | | | L2 | | | L3 | | | L4 | | | L5 | | | L6 | | | G7 | | | G8 | | | G9 | | | G10 | | | R11 | | | R12 | | | R13 | | | N14 | | | N15 | | | N16 | | |
|---|---|---|---|---|---|---|---|---|---|---|---|---|---|---|---|---|---|---|---|---|---|---|---|---|---|---|---|---|---|---|---|---|---|---|---|---|---|---|---|---|---|---|---|---|---|---|---|---|---|---|---|
| Compound | B | A | DE | B | A | DE | B | A | DE | B | A | DE | B | A | DE | B | A | DE | B | A | DE | B | A | DE | B | A | DE | B | A | DE | B | A | DE | B | A | DE | B | A | DE | B | A | DE | B | A | DE | B | A | DE | B | A | DE |
| 1 | 1 | 4 | 0 | 5 | 5 | 0 | 7 | 2 | 0 | 0 | 0 | 0 | 0 | 5 | 0 | 0 | 9 | 0 | 0 | 0 | 0 | 0 | 3 | 0 | 0 | 0 | 0 | 0 | 6 | 0 | 0 | 5 | 0 | 0 | 9 | 3 | 0 | 6 | 0 | 0 | 0 | 0 | 1 | 5 | 0 | 0 | 1 | 1 | 0 | 3 | 1 |
| 2 | 1 | 7 | 0 | 0 | 0 | 0 | 13 | 6 | 0 | 0 | 3 | 0 | 0 | 4 | 0 | 0 | 4 | 0 | 0 | 5 | 0 | 0 | 6 | 0 | 0 | 21 | 0 | 0 | 21 | 0 | 0 | 4 | 0 | 0 | 11 | 0 | 2 | 13 | 1 | 0 | 1 | 0 | 2 | 5 | 0 | 0 | 3 | 0 | 6 | 13 | 0 |
| 3 | 5 | 11 | 0 | 0 | 0 | 0 | 80 | 0 | 0 | 0 | 17 | 0 | 0 | 0 | 0 | 0 | 22 | 0 | 1 | 18 | 0 | 2 | 15 | 0 | 1 | 42 | 1 | 0 | 15 | 0 | 2 | 16 | 0 | 3 | 14 | 4 | 19 | 15 | 0 | 12 | 6 | 0 | 7 | 13 | 0 | 2 | 0 | 1 | 0 | 2 | 2 |
| 4 | 6 | 7 | 0 | 0 | 0 | 0 | 74 | 0 | 0 | 2 | 2 | 0 | 0 | 0 | 0 | 0 | 18 | 0 | 0 | 21 | 0 | 2 | 0 | 0 | 1 | 19 | 0 | 1 | 6 | 0 | 2 | 10 | 0 | 1 | 23 | 1 | 12 | 18 | 0 | 9 | 0 | 0 | 15 | 1 | 0 | 2 | 2 | 0 | 0 | 13 | 5 |
| 5 | 4 | 2 | 0 | 0 | 0 | 0 | 92 | 0 | 0 | 0 | 0 | 0 | 0 | 0 | 0 | 0 | 0 | 0 | 0 | 0 | 0 | 0 | 5 | 0 | 1 | 9 | 0 | 1 | 5 | 0 | 1 | 0 | 0 | 0 | 3 | 0 | 2 | 2 | 2 | 8 | 0 | 0 | 4 | 0 | 0 | 1 | 1 | 0 | 0 | 3 | 0 |

| | Neurons | | | L1 | | | L2 | | | L3 | | | L4 | | | L5 | | | L6 | | | G7 | | | G8 | | | G9 | | | G10 | | | R11 | | | R12 | | | R13 | | | N14 | | | N15 | | | N16 | | |
|---|---|---|---|---|---|---|---|---|---|---|---|---|---|---|---|---|---|---|---|---|---|---|---|---|---|---|---|---|---|---|---|---|---|---|---|---|---|---|---|---|---|---|---|---|---|---|---|---|---|---|---|
| Compound | B | A | DE | B | A | DE | B | A | DE | B | A | DE | B | A | DE | B | A | DE | B | A | DE | B | A | DE | B | A | DE | B | A | DE | B | A | DE | B | A | DE | B | A | DE | B | A | DE | B | A | DE | B | A | DE | B | A | DE |
| 1 | 1 | 6 | 0 | 0 | 0 | 0 | 0 | 0 | 0 | 3 | 0 | 0 | 0 | 0 | 0 | 0 | 0 | 0 | 1 | 0 | 0 | 0 | 13 | 0 | 0 | 26 | 0 | 0 | 11 | 1 | 0 | 1 | 0 | 1 | 7 | 2 | 0 | 6 | 0 | 3 | 0 | 0 | 0 | 2 | 0 | 1 | 1 | 1 | 0 | 9 | 0 |
| 2 | | | | | | | | | | | | | | | | | | | | | | | | | | | | | | | | | | | | | | | | | | | | | | | | | | | |
| 3 | 1 | 6 | 1 | 0 | 0 | 0 | 67 | 0 | 0 | 0 | 0 | 0 | 0 | 0 | 0 | 0 | 7 | 0 | 0 | 0 | 0 | 0 | 0 | 0 | 0 | 20 | 0 | 0 | 15 | 0 | 0 | 2 | 0 | 0 | 9 | 12 | 0 | 7 | 0 | 1 | 0 | 0 | 1 | 0 | 0 | 0 | 0 | 6 | 0 | 0 | 0 |
| 4 | 2 | 2 | 0 | 0 | 0 | 0 | 44 | 0 | 0 | 0 | 0 | 0 | 0 | 0 | 0 | 0 | 3 | 0 | 0 | 0 | 0 | 0 | 2 | 0 | 3 | 3 | 0 | 2 | 1 | 0 | 1 | 3 | 0 | 1 | 4 | 0 | 2 | 0 | 0 | 2 | 2 | 0 | 1 | 3 | 0 | 1 | 3 | 0 | 3 | 2 | 0 |
| 5 | 10 | 4 | 1 | 0 | 0 | 0 | 76 | 0 | 0 | 9 | 0 | 0 | 14 | 0 | 0 | 4 | 0 | 0 | 0 | 0 | 0 | 0 | 8 | 2 | 1 | 22 | 2 | 0 | 16 | 0 | 3 | 7 | 0 | 3 | 5 | 1 | 23 | 0 | 2 | 7 | 0 | 0 | 30 | 0 | 0 | 4 | 1 | 0 | 2 | 3 | 8 |

**B** -block/attenuation **A** -amplification **DE** -direct effects

0% 100%

**Figure 3-table supplement 2.** Census of effects elicited by compounds **1-5** on $K^{+}$-induced depolarization in 16 DRG neuronal subtypes screened in calcium imaging experiments. Intracellular calcium ($Ca^{2+}$) depolarization was induced by application of 30 mM K solution. The ability of the compounds (at 20 µM) to modulate the $Ca^{2+}$ signal was scored as either a block or amplification immediately following compound incubation. Direct effects to the $Ca^{2+}$ baseline with sample incubation were also scored. For each calcium imaging experiment the compound was applied twice. Only reproducible effects elicited by the compound were scored. The effects for each subtype are represented as percentages (number of cells affected/total number of cells x 100) on the table.

| | Neurons | | | L1 | | | L2 | | | L3 | | | L4 | | | L5 | | | L6 | | | G7 | | | G8 | | | G9 | | | G10 | | | R11 | | | R12 | | | R13 | | | N14 | | | N15 | | | N16 | | |
|---|---|---|---|---|---|---|---|---|---|---|---|---|---|---|---|---|---|---|---|---|---|---|---|---|---|---|---|---|---|---|---|---|---|---|---|---|---|---|---|---|---|---|---|---|---|---|---|---|---|---|---|
| Compound | B | A | DE | B | A | DE | B | A | DE | B | A | DE | B | A | DE | B | A | DE | B | A | DE | B | A | DE | B | A | DE | B | A | DE | B | A | DE | B | A | DE | B | A | DE | B | A | DE | B | A | DE | B | A | DE | B | A | DE |
| 1 | 1 | 3 | 0 | 20 | 0 | 0 | 0 | 0 | 0 | 0 | 0 | 0 | 9 | 0 | 0 | 0 | 0 | 0 | 2 | 0 | 0 | 6 | 0 | 0 | 0 | 6 | 0 | 0 | 8 | 2 | 3 | 0 | 0 | 6 | 9 | 0 | 0 | 0 | 0 | 0 | 0 | 0 | 2 | 0 | 0 | 3 | 1 | 1 | 0 | 5 | 0 |
| 2 | 0 | 15 | 0 | 0 | 0 | 0 | 0 | 0 | 0 | 0 | 50 | 0 | 0 | 0 | 0 | 0 | 8 | 0 | 0 | 29 | 0 | 0 | 32 | 0 | 0 | 22 | 0 | 0 | 22 | 0 | 0 | 7 | 0 | 0 | 7 | 0 | 0 | 25 | 0 | 1 | 8 | 0 | 0 | 10 | 0 | 0 | 6 | 2 | 0 | 0 | 0 |
| 3 | 0 | 1 | 0 | 0 | 0 | 0 | 0 | 0 | 0 | 0 | 0 | 0 | 0 | 0 | 0 | 3 | 3 | 3 | 0 | 9 | 2 | 0 | 7 | 0 | 0 | 0 | 0 | 0 | 0 | 0 | 0 | 0 | 0 | 0 | 0 | 0 | 0 | 0 | 0 | 2 | 0 | 0 | 0 | 0 | 0 | 0 | 2 | 0 | 0 | 1 | 0 |
| 4 | 0 | 4 | 0 | 0 | 0 | 0 | 0 | 0 | 0 | 0 | 0 | 0 | 0 | 0 | 0 | 0 | 0 | 0 | 0 | 0 | 0 | 0 | 0 | 0 | 0 | 7 | 0 | 0 | 8 | 0 | 0 | 0 | 0 | 0 | 6 | 0 | 0 | 0 | 0 | 0 | 2 | 0 | 0 | 1 | 0 | 0 | 2 | 0 | 0 | 12 | 0 |
| 5 | 0 | 12 | 0 | 0 | 17 | 0 | 0 | 0 | 0 | 0 | 0 | 0 | 0 | 0 | 0 | 0 | 23 | 0 | 0 | 5 | 0 | 0 | 0 | 0 | 0 | 44 | 1 | 0 | 38 | 0 | 0 | 28 | 0 | 0 | 19 | 0 | 0 | 0 | 0 | 0 | 2 | 0 | 0 | 1 | 0 | 0 | 9 | 1 | 0 | 4 | 4 |

**B** -block/attenuation **A** -amplification **DE** -direct effects

0% 100%

**Figure 6-table supplement 1**. Census of effects elicited by **3** on MRS2365-induced depolarization in 16 DRG neuronal subtypes screened in two calcium imaging experiments. Intracellular calcium ($Ca^{2+}$) depolarization was induced by application of 100 mM MRS2365 solution (a specific P2Y1 agonist). The number of cell responses to MRS2365 that were blocked by **3** (20 μM and 180 μM) were scored as % block responses.

| | Neurons | L1 | L2 | L3 | L4 | L5 | L6 | G7 | G8 | G9 | G10 | R11 | R12 | R13 | N14 | N15 | N16 |
|---|---|---|---|---|---|---|---|---|---|---|---|---|---|---|---|---|---|
| Cells | 1826 | 2 | 21 | 40 | 20 | 46 | 29 | 38 | 262 | 242 | 113 | 136 | 269 | 309 | 165 | 62 | 48 |
| % responded | 3.0 | 0.0 | 100.0 | 20.0 | 5.0 | 2.2 | 3.4 | 0.0 | 0.8 | 0.4 | 3.5 | 0.7 | 1.1 | 0.0 | 4.2 | 0.0 | 0.0 |
| % Block at 20 uM | 1.2 | 0.0 | 71.4 | 2.5 | 0.0 | 0.0 | 0.0 | 0.0 | 0.0 | 0.0 | 1.8 | 0.0 | 0.0 | 0.0 | 1.2 | 0.0 | 0.0 |
| % Block at 180 uM | 1.3 | 0.0 | 85.7 | 2.5 | 0.0 | 0.0 | 0.0 | 0.0 | 0.0 | 0.0 | 1.8 | 0.0 | 0.0 | 0.0 | 1.2 | 0.0 | 0.0 |

| | Neurons | L1 | L2 | L3 | L4 | L5 | L6 | G7 | G8 | G9 | G10 | R11 | R12 | R13 | N14 | N15 | N16 |
|---|---|---|---|---|---|---|---|---|---|---|---|---|---|---|---|---|---|
| Cells | 1296 | 3 | 20 | 22 | 19 | 46 | 26 | 23 | 188 | 90 | 140 | 62 | 72 | 455 | 80 | 29 | 13 |
| % responded | 2.2 | 0.0 | 75.0 | 0.0 | 5.3 | 2.2 | 4.0 | 0.0 | 1.6 | 0.0 | 0.0 | 0.0 | 0.0 | 0.0 | 4.2 | 0.0 | 0.0 |
| % Block at 20 uM | 1.0 | 0.0 | 65.0 | 0.0 | 5.3 | 0.0 | 0.0 | 0.0 | 0.0 | 0.0 | 0.0 | 0.0 | 0.0 | 0.0 | 1.2 | 0.0 | 0.0 |
| % Block at 180 uM | 1.0 | 0.0 | 75.0 | 0.0 | 5.3 | 0.0 | 0.0 | 0.0 | 0.0 | 0.0 | 0.0 | 0.0 | 0.0 | 0.0 | 1.2 | 0.0 | 0.0 |

0% 100%
